# Transcriptional milestones in *Dictyostelium* development

**DOI:** 10.1101/2021.05.27.445976

**Authors:** Mariko Katoh-Kurasawa, Karin Hrovatin, Shigenori Hirose, Amanda Webb, Hsing-I Ho, Blaž Zupan, Gad Shaulsky

## Abstract

Development of the social amoeba *Dictyostelium discoideum* begins by starvation of single cells and ends in multicellular fruiting bodies 24 hours later. These major morphological changes are accompanied by sweeping gene expression changes, encompassing nearly half of the 13,000 genes in the genome. To explore the relationships between the transcriptome and developmental morphogenesis, we performed time-series RNA-sequencing analysis of the wild type and 20 mutant strains with altered morphogenesis. These strains exhibit arrest at different developmental stages, accelerated development, or terminal morphologies that are not typically seen in the wild type. Considering eight major morphological transitions, we identified 1,371 milestone genes whose expression changes sharply between two consecutive transitions. We also identified 1,099 genes as members of 21 regulons, which are groups of genes that remain coordinately regulated despite the genetic, temporal, and developmental perturbations in the dataset. The gene annotations in these milestones and regulons validate known transitions and reveal several new physiological and functional transitions during development. For example, we found that DNA replication genes are co-regulated with cell division genes, so they are co-expressed in mid-development even though chromosomal DNA is not replicated at that time. Altogether, the dataset includes 486 transcriptional profiles, across developmental and genetic conditions, that can be used to identify new relationships between gene expression and developmental processes and to improve gene annotations. We demonstrate the utility of this resource by showing that the cycles of aggregation and disaggregation observed in allorecognition-defective mutants involve a dedifferentiation process. We also show unexpected variability and sensitivity to genetic background and developmental conditions in two commonly used genes, *act6* and *act15*, and robustness of the *coaA* gene. Finally, we propose that *gpdA* should be used as a standard for mRNA quantitation because it is less sensitive to genetic background and developmental conditions than commonly used standards. The dataset is available for democratized exploration without the need for programming skills through the web application dictyExpress and the data mining environment Orange.

## Introduction

*Dictyostelium discoideum* is a social amoeba that lives in the soil and feeds on bacteria (Kessin 2001). The vegetative stage of life is mainly solitary, although predation can be cooperative (Rubin et al. 2019). Upon starvation, the cells begin to cooperate in a developmental process that leads to fruiting body formation. In the first few hours of starvation, the cells produce and secrete cyclic adenosine monophosphate (cAMP), which is used as an extracellular chemoattractant and intracellular second messenger, mainly through activation of cAMP-dependent protein kinase (PKA) at various developmental stages (Loomis 1998; Ritchie et al. 2008). The starving cells begin to aggregate as they form ripples and stream toward aggregation centers, forming loose aggregates of about 50,000 cells each (Gomer et al. 2011). They then differentiate into two major cell types, prespore and prestalk, in a spatially independent manner and eventually sort into two main compartments as a tight aggregate is formed. The next morphological transition is tip formation, when some prestalk cells move to the top of the tight aggregate. The tip leads elongation of the aggregate into a finger-like structure that can topple over and form a migrating slug. After a few more hours, the slug erects itself and its posterior shortens and widens as the structure assumes a Mexican hat shape. The prestalk cells at the tip of the Mexican hat vacuolize and encase themselves in a cell wall, pushing each other down through the prespore cell mass toward the solid substrate (Loomis 1975). The emerging stalk lifts the structure off of the substrate as the prespore cells encapsulate and desiccate themselves to become spores during the culmination process. Development ends with a mature fruiting body that is about 1 mm tall, 24 h after the onset of starvation (Kessin 2001).

*Dictyostelium* morphogenesis is readily amenable to genetic analysis. Early studies showed that mutations can abrogate the aggregation stage, leading to stable variants that grow well but fail to aggregate or develop further (Sussman 1952). Subsequent studies discovered dozens of mutant strains that exhibited arrest at every possible morphological stage, as well as strains that exhibited altered developmental timing and even morphologies that are not seen in the wild type (Loomis 1975). Many mutations that have profound effects on development have little or no effect on cell growth during the vegetative stage (Loomis 1978). These early studies were bolstered by subsequent observations that many of the vegetative genes are not expressed during development and many developmental genes are not expressed during growth (Parikh et al. 2010). These findings and the fact that *Dictyostelium* cells are usually haploid (Sussman and Sussman 1962) have led to the discovery of numerous developmental genes and mutations that arrest development at various stages (Kuspa 2006).

Dictoystelid development is accompanied by evolutionarily conserved changes in morphology and transcription (Parikh et al. 2010; Glockner et al. 2016; Schilde et al. 2016). Previous studies have shown that the *D. discoideum* transcriptome is a quantitative phenotype that can be used to predict developmental stages and to infer epistatic relationships (Van Driessche et al. 2005). They also suggested correlations between the transcriptome and morphogenesis (Good et al. 2003; Cai et al. 2014; Katoh-Kurasawa et al. 2016). Nevertheless, transcriptional and morphological changes occur in bursts that are not always coordinated. For example, some of the biggest changes in gene expression occur before aggregation and visible morphogenesis begin, whereas the vast morphological changes of culmination are accompanied by only a few transcriptional changes (Rosengarten et al. 2015).

Our goal in this study was to explore the relationship between morphogenesis and the transcriptome. We chose 20 mutant strains that exhibit various developmental abnormalities as a set of genetic and developmental perturbations, and analyzed them at various times, as temporal perturbations. We hypothesized that links between the transcriptome and morphogenesis would identify markers of specific morphological transitions. We also sought groups of co-regulated genes whose coordinate regulation is robust against the three perturbations. Our final goal was to make this large dataset available for easy exploration by the research community on the web-based application dictyExpress (Stajdohar et al. 2017) and in the data mining environment Orange (Demsar et al. 2013). Neither of these exploration options require programming skills.

## Results

### Transcriptome signatures match morphological progression during development

To explore the relationship between morphogenesis and gene expression, we compared the developmental transcriptomes of the wild type (AX4) to 20 strains that represent genetic perturbations in different developmental processes. AX4 and the mutant strains belong to seven morphological groups: wild type (WT; AX4 and MybBGFP), culmination defective (cul def/cud; *gtaI^−^*, *gtaG^−^*, *cudA^−^*, *dgcA^−^*, and *ecmA:Rm*), tight-aggregate arrest (tag arrest; *tagB*^−^ and *comH^−^*), aggregationless (agg-; *gtaC^−^*, *acaA^−^*, *mybB^−^*, and *amiB^−^*), precocious development (precocious/prec; *pkaC^oe^*, and *pkaR^−^*), small fruiting body (small FB/sFB; *pkaC^oe^* in two adenylate cyclase knockout strains), and tight-aggregate/loose-aggregate disaggregation (disagg; *tgrB1^−^*, *tgrB1^−^/tgrC1^−^*, *tgrC1^−^*, and *gbfA^−^*) (Table S1). We developed the cells, photographed them, and performed RNA-seq analysis on all the strains at similar time intervals. The morphological stages of representative strains are shown in Fig. 1. In the wild type, cell populations appear morphologically unchanged for the first six hours of starvation. At eight hours, they appear as streams and loose aggregates, followed by tight aggregates at 12 h, slugs at 16 h, culminants at 20 h, and fruiting bodies at 24 h (Fig. 1). Most of the culmination-defective strains develop indistinguishably from the wild type until the slug stage, where they arrest and fail to exhibit further morphological changes for the next eight hours (cul def, Fig. 1). Cells of the tag-arrest group develop to the tight aggregate stage at 12 h and arrest there (tag arrest, Fig. 1), and cells of the aggregationless group do not exhibit gross morphological changes (agg-, Fig. 1).

**Fig. 1.**
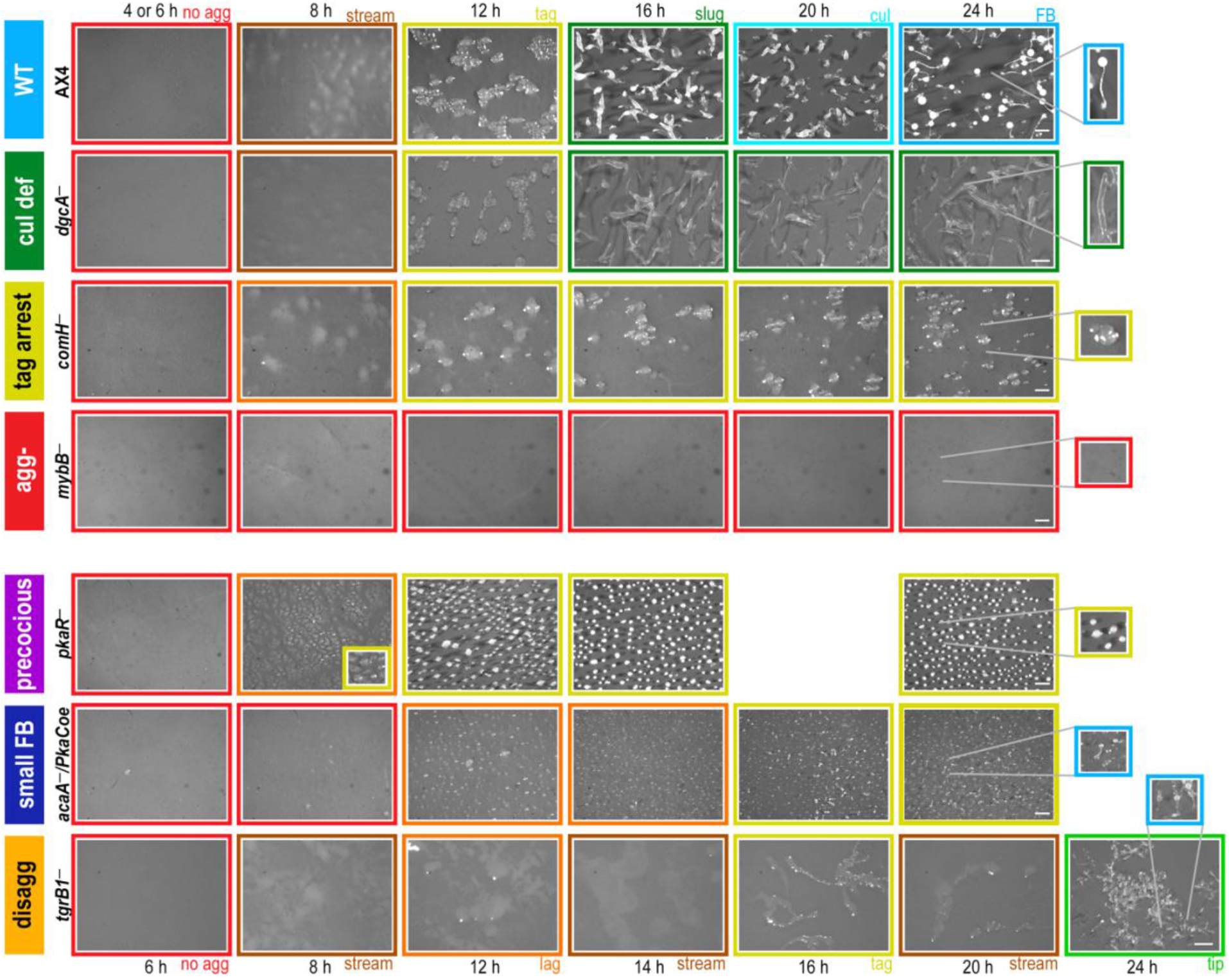
Developmental morphologies in different phenotype groups. Cells were starved and developed on black nitrocellulose filters. The phenotype groups and strain names are indicated on the left: wild type (WT, light blue), culmination defective (cul def, dark green), tight aggregate arrest (tag arrest, dark yellow), aggregationless (agg-, red), precocious development (precocious, violet), small fruiting body (small FB, dark blue), and tight-aggregate/loose-aggregate disaggregation (disagg, orange). Developmental structures were photographed at the times (hours) indicated above the panels with the exception of the disaggregation group in which times are indicated below. The frame colors indicate the most representative morphological stage: no aggregation (red), rippling/streams (brown), loose aggregate (orange), tight aggregate (dark yellow), tipped aggregate (light green), slug/first finger (dark green), culminant (cyan) and fruiting body (light blue). Bars = 0.5 mm. The most advanced developmental structures are shown in the 2-fold magnified images. Images brightness was adjusted according to the variable color of the nitrocellulose filters.

Unlike mutants that exhibit developmental arrests with morphologies that are equivalent to wild-type stages, some mutants exhibit new and/or dynamic terminal morphologies. The precocious mutants exhibit accelerated morphogenesis at early stages, followed by a tight aggregate arrest (*pkaR^−^*, Fig. 1) or precocious culmination (*pkaC^oe^*, Supplemental Fig S1), both associated with spore production. The small fruiting body mutants exhibit sparse aggregation and a mix of stages at the end of development, including mostly tight aggregates and occasional small fruiting bodies (small FB, Fig. 1). The disaggregation mutants initiate aggregation like the wild type, but begin to exhibit cycles of aggregation and disaggregation at 8-12 h of development. Some disaggregation strains (e.g. *tgrB1^−^*) exhibit a mix of terminal morphologies, including loose aggregates, tight aggregates, and a few small fruiting bodies (disagg, Fig. 1), whereas others (e.g. *tgrC1^−^*) appear as loose aggregates at the end of development (Dynes et al. 1994).

This set of strains and time points represents genetic and temporal perturbations that result in various morphological stages. To test whether each morphological stage is accompanied by a unique transcriptional profile, we analyzed replicate samples by RNA-seq. Altogether, the dataset includes 486 transcriptional profiles. The multidimensional scaling (MDS) plot in Fig. 2A shows the developmental series of the respective transcriptomes in each strain. Individual strain trajectories are shown separately in Supplemental Fig S2A. The wild-type (AX4 and MybBGFP) transcriptomes follow a nearly linear path in the direction of the x-axis, suggesting that this direction largely represents normal developmental progression in time (cyan, Fig. 2A and Supplemental Fig S2A). In the agg- group, all the transcriptomes clustered close to the wild type at 0 h (red, Fig. 2A and Supplemental Fig S2A), suggesting that the aggregationless transcriptomes are similar to the wild-type pre-aggregation transcriptome and that they do not change much after starvation. The fact that four different mutations and 12 different time points exhibit similar transcriptome signatures supports the idea that the morphological stage matches the transcriptional phenotype.

**Fig. 2.**
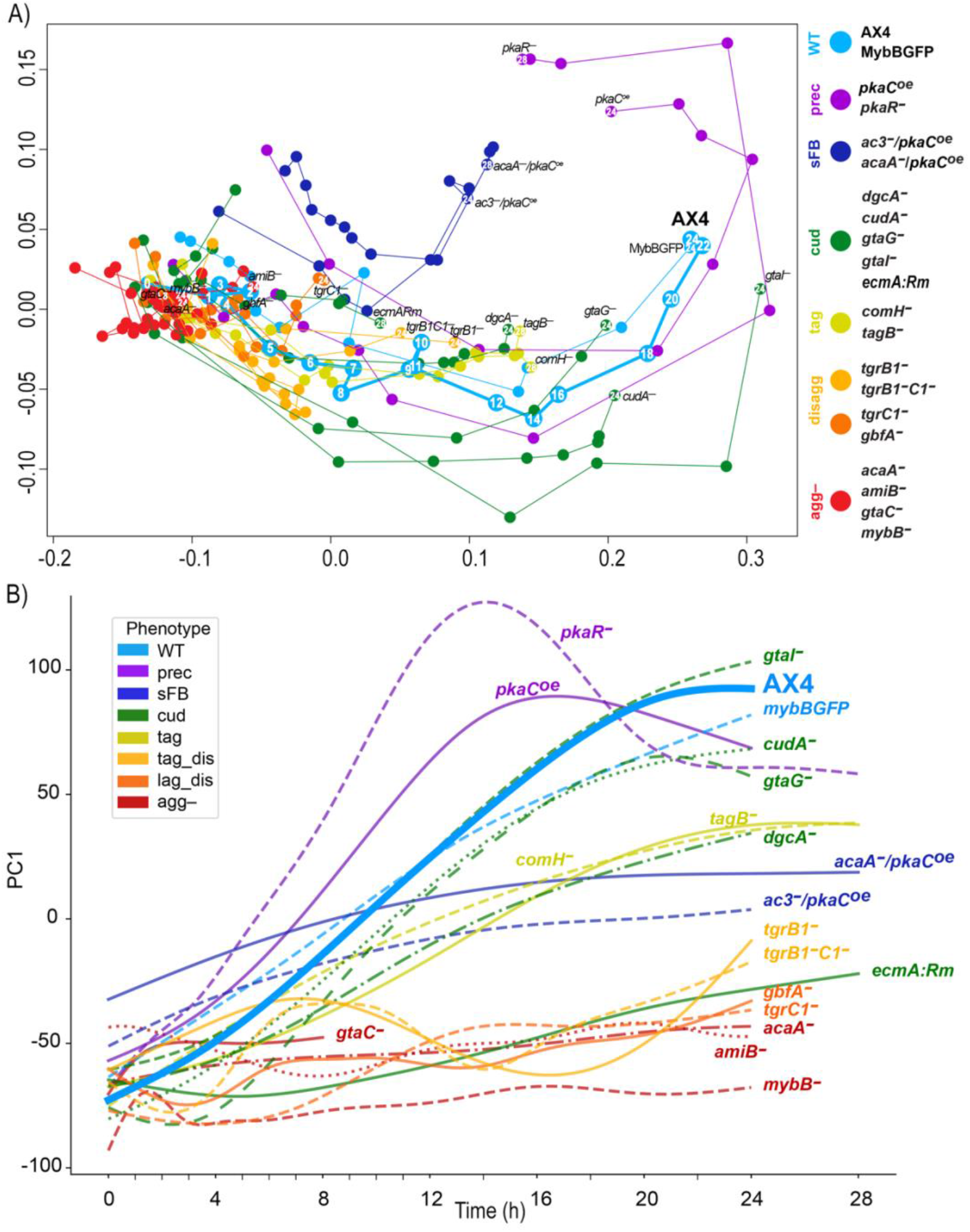
Characteristic transcriptome patterns of different phenotype groups. We analyzed the transcriptomes of the developing cells across time by RNA-seq and used dimensionality reduction techniques to compare the different strains. A) In the MDS plot, each circle represents the average transcriptome of 2-7 replicates at a certain time point of a strain. The arbitrary units of the x and y axes reflect the distances between the transcriptomes such that the proximity between the circles approximates their similarity (adjacent circles are similar to one another). The circle and line colors represent the phenotype group of the strain and the strain name is indicated next to the last time point, which is indicated inside the circle (hours). The circles are connected in temporal order. The phenotype groups are wild type (WT, light blue), precocious development (prec, violet), small fruiting body (sFB, dark blue), culmination defective (cud, dark green), tight aggregate arrest (tag, dark yellow), tight aggregate / loose aggregate disaggregation (disagg, light and dark orange, respectively) and aggregationless (agg-, red). The strain names in each phenotype group are indicated next to the terminal time point. Developmental time points are indicated in each of the AX4 circles and in the last time point of each strain. B) We performed PCA and plotted PC1 (y-axis, arbitrary units) against time (x-axis, hours) of each strain. PC1 accounts for 30.4% of the variation. The strain names are indicated in the plot and the color represents the phenotype group (see legend). The wild type (AX4) is shown as a thick line and bold text.

The transcriptional trajectories of the disaggregation strains also started near the wild-type transcriptome at 0 h and followed a path similar to the wild type for the first few hours. They then seemed to halt around the 8 h-time point of the wild type (dark and light orange, Fig. 2A and Supplemental Fig S2A), which matches the loose aggregate stage. Similarly, the tight aggregate mutant transcriptomes progressed like the wild type until the 12 h samples (yellow, Fig. 2A and Supplemental Fig S2A), which is the tight aggregate stage. The culmination-defective strains, *gtaG^−^*, *dgcA^−^*, and *cudA*^−^ (green, Fig. 2A and Supplemental Fig S2A), followed a near wild-type trajectory that halted around the AX4 16 h transcriptome, which matches the slug stage. Nevertheless, the culmination-defective strain *ecmA:Rm* exhibited an arrest around the 8 h wild-type stage, and the *gtaI*^−^ trajectory departed rather broadly from the wild type, mainly along the y-axis (green, Fig. 2A and Supplemental Fig S2A).

The precocious mutants, *pkaR*^−^ and *pkaC^oe^* (purple, Fig. 2A and Supplemental Fig S2A), exhibited large intervals between adjacent time points along the x-axis, consistent with the accelerated developmental phenotype. Moreover, their transcriptomes did not converge on the terminal transcriptomes of the wild type, but continued to diverge, mainly along the y-axis. One of the major differences between *pkaR*^−^ and *pkaC^oe^* was that *pkaR*^−^ exhibited a near wild-type pattern at 0 h, whereas *pkaC^oe^* was different, matching the 0 h-time points of the other two sFB strains that also overexpress *pkaC*. The developmental transcriptomes of the sFB strains, *acaA^−^*/*pkaC^OE^* and *ac3^−^*/*pkaC^OE^* (blue, Fig. 2A and Supplemental Fig S2A), exhibited shorter distances between adjacent time points compared to the wild type and an arrest along the x-axis around 12 h, which is the tight-aggregate stage, consistent with their major terminal morphologies. Altogether, these observations suggest that the transcriptome profile is largely consistent with the morphological stage with a few exceptions that are mostly related to PKA mis-regulation.

Our dataset is somewhat skewed because most of the mutants arrest development before the fruiting body stage. This skew could have affected the MDS visualization, so we employed an additional method to visualize the data. We performed principal component analysis (PCA) on the wild-type (AX4) data and used the PC1 embeddings to transform the data from the remaining strains. We then plotted the PC1 trendlines against time together with the AX4 data (Fig. 2B) and separately for each strain (Supplemental Fig S2B). The AX4 trendline showed a largely direct correlation between PC1 and time, suggesting that areas above the trendline indicate accelerated development and areas below it indicate attenuated development. Indeed, the embedded PC1 values of the agg- group were relatively constant over time and largely below the AX4 graph, consistent with the inability of the strains to aggregate and develop (red, Fig. 2B and Supplemental Fig S2B). The trendlines of the tag-arrest group showed a direct correlation between time and embedded PC1, similar to AX4, but their slopes were shallower than AX4 and they plateaued near the 12 h value of AX4 (yellow, Fig. 2B and Supplemental Fig S2B), suggesting that developmental progression in this group slowed down after aggregation began, and finally ceased at the tight aggregate stage. This transcriptional trend matches well with the morphological progression of this group (Fig. 1). The trendlines of the disaggregation groups (light and dark orange, Fig. 2B and Supplemental Fig S2B) oscillated between the values of AX4 at 8 h and at 2-3 h, suggesting that the transcriptional state of these strains was reversed after about 8 h of development and resumed after about 14 h. The trendlines of the culmination-defective group varied among strains, but most of them were below the AX4 line. Those of *gtaG*^−^ and *cudA*^−^ plateaued around the 16 h value of AX4, which corresponds to the slug stage (green, Fig. 2B and Supplemental Fig S2B). The trendlines of the precocious mutants (purple, Fig. 2B and Supplemental Fig S2B) were above the AX4 line up to 12-16 h, when they peaked. Afterwards, these transcriptomes dipped below the AX4 trendline. The small fruiting body mutants (blue, Fig. 2B and Supplemental Fig S2B) exhibited shallow slopes that crossed the AX4 line at 8-10 h, suggesting an early start during aggregation and slow progress afterwards. Overall, the PC1 values corresponded well to the morphological progression, confirming and extending the MDS analysis conclusions.

The most prominent exception to this rule was the *pkaR*^−^ strain, which exhibited a tight-aggregate terminal morphology (Fig. 1) while exhibiting transcriptional profiles that were much more advanced than the other tag mutants, resembling and exceeding the profiles of the wild type (Fig. 2). Mutations in *pkaR* accelerate development and uncouple spore and stalk cell differentiation from morphogenesis (Abe and Yanagisawa 1983). Morphological examination of the *pkaR*^−^ tight aggregates revealed many spores and occasional vacuolized stalk cells as early as 16 h, about eight hours before the wild type. In AX4, spores and stalk cells were not observed at 16 h, as expected, but both were abundant later. The *pkaR*^−^ spores seemed compromised, as suggested by their phase-dim appearance (Supplemental Fig S3A and B). These findings suggest that, although gross morphology was disrupted in the *pkaR*^−^ mutant, fine morphogenesis and precocious differentiation were still consistent with the transcriptome profiles. Therefore, the transcriptional patterns are good predictors of developmental stage.

### Milestone genes define developmental stage boundaries

Transcripts that undergo sharp changes during developmental transitions can be thought of as developmental milestones. To identify such transcripts, we performed combined differential expression analyses on the wild-type transcriptome data. First, we annotated each sample in the AX4 dataset based on the most abundant morphological stage. We then carried out differential expression analysis to identify transcripts that exhibited significant differences between two consecutive developmental stages. We selected transcripts whose trajectories changed only once or twice during development to eliminate highly fluctuating genes. Altogether, we found 1,371 milestone genes at eight stage transitions (Fig. 3). Finally, we performed gene-set enrichment analysis on the milestone groups to identify potential common functions (Supplemental File S1).

**Fig. 3.**
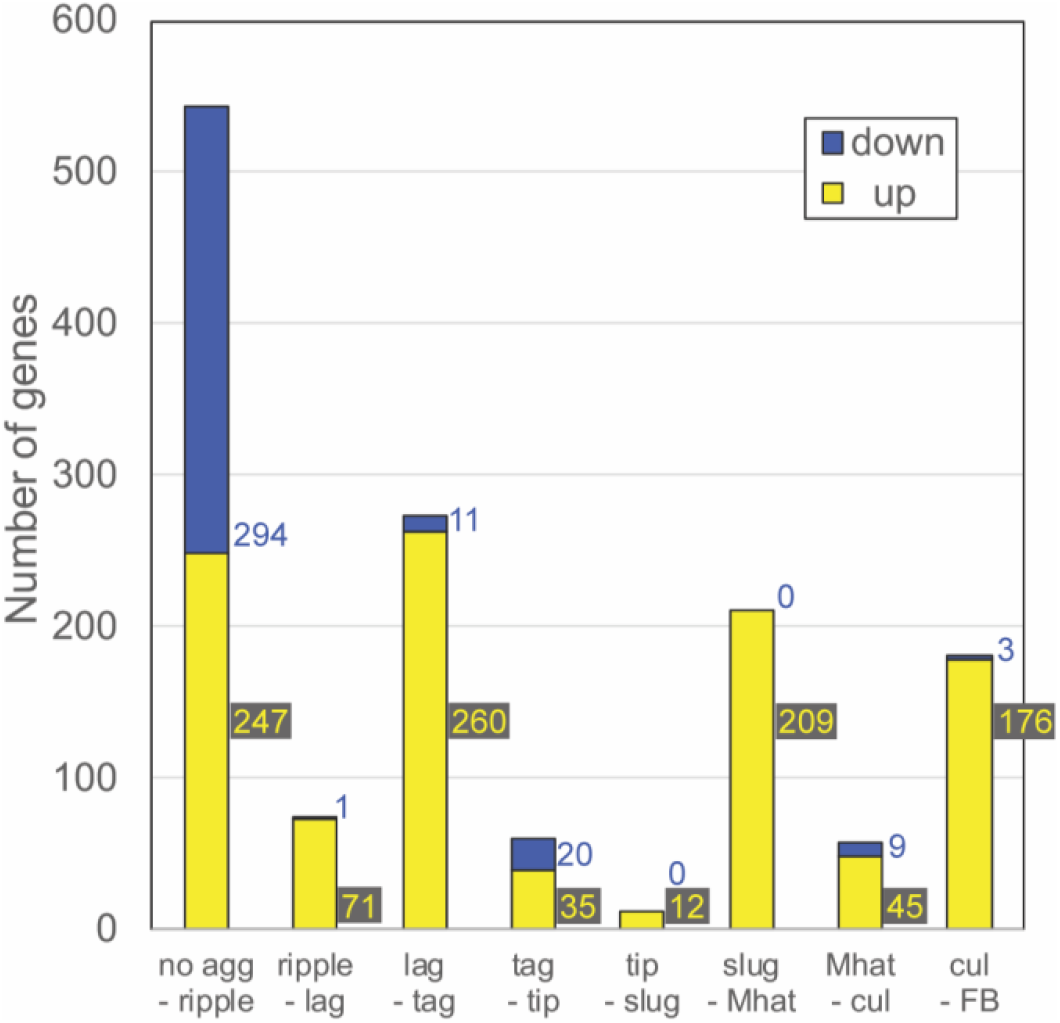
The number of milestone genes at each developmental transition. We performed differential expression analysis between samples across each developmental transition in AX4 (WT) development and selected transcripts that exhibited significant change at each transition. The graph shows the number of differentially expressed genes at each stage transition in AX4 (WT). Down-regulated genes (blue): mRNA abundance at the early stage is significantly greater than mRNA abundance at the later stage. Up-regulated genes (yellow): mRNA abundance at the early stage is significantly lower than mRNA abundance at the later stage. The stage transitions are indicated below each bar. Colored numbers next to the bars indicate the number of genes in each ‘down’ and ‘up’ category.

Most of the milestone genes were up-regulated except for the transition from no aggregation to ripples (Fig. 3). This transition was marked by the largest number of milestone genes: 294 genes were down-regulated and 247 genes were up-regulated. This finding is consistent with the largest developmental transition in gene expression (Parikh et al. 2010), which corresponds to the cessation of growth and the onset of aggregation. About 200 up-regulated milestone genes and almost no down-regulated genes were found at the transitions from loose aggregate to tight aggregate (lag-tag), slugs to Mexican hats (slug-Mhat), and culminant to fruiting body (cul-FB) (Fig. 3). The other transitions (ripple-lag, tag-tip, tip-slug, and Mhat-cul) were accompanied by small numbers of milestone genes (Fig. 3).

In the no agg-ripple group, the down-regulated genes were largely associated with metabolic/biosynthesis pathways (Supplemental File S1). The lag-tag transition was highly enriched in DNA replication, chromosome segregation, and mismatch repair annotations as well as some *srfA* induced genes (sig), cAMP-pulse induced genes, and prespore (psp) genes (Supplemental File S1). The tag-tip down-regulated group was enriched in cell adhesion, cytoskeleton organization, and signal transduction genes, and the up-regulated group included members of the 57-aa protein family, gtaG-dependent short proteins, cAMP-pulse induced genes, and prestalk (pst) genes (Supplemental File S1). The tip-slug up-regulated group included hssA/2C/7E family genes (Supplemental File S1). The slug-Mhat group was highly enriched in gtaG-dependent short proteins and psp genes (Supplemental File S1), and some hssA/2C/7E family genes. The psp genes were down-regulated in the subsequent Mhat-cul group (Supplemental File S1). The cul-FB up-regulated group was enriched in sugar metabolism, differentiation, and morphogenesis genes (Supplemental File S1). The variable numbers of milestone genes at different developmental transitions (Fig. 3) confirm and extend previous findings (Rosengarten et al. 2015). This variability suggests that some morphological transitions are accompanied by more gradual changes in gene expression, which would not be consistent with the milestone definition, and that some morphological transitions do not require large changes in gene expression. The gene-set enrichment analysis (Supplemental File S1) is consistent with known functional changes during development (Kessin 2001; Parikh et al. 2010). Details about the gene sets described above are provided in Supplemental File S5, section 5.1. and Supplemental File S8.

### Milestone gene response to genetic perturbation of development

The milestone genes were identified in the AX4 data as genes whose transcript abundance is sharply increased or decreased during a transition between two consecutive stages. We predicted that they would not exhibit sharp transcriptional changes in mutants that fail to undergo the respective morphological transitions. To test the prediction, we plotted the expression patterns as heatmaps across the mutant strains (Fig. 4). Indeed, the milestone genes were expressed similarly to the AX4 patterns in the mutant strains until the respective developmental arrests, with a few interesting exceptions. The down-regulated transcripts in the no agg-ripple milestone exhibited a marked reduction in abundance in most phenotype groups and no reduction in the aggregationless mutants, as expected. The exceptions were the small fruiting body mutant *acaA/pkaC^oe^* and the disaggregation strain *gbfA^−^*. Reduction was observed in *tgrC1*^−^ and the culmination-defective strain *ecmA:Rm*, but to a lesser extent. The up-regulated transcripts in the no agg-ripple milestone genes were sharply induced in most of the strains but not in the aggregationless mutants, as expected. Notable exceptions were the small fruiting body mutant *acaA/pkaC^oe^*, the culmination-defective strain *ecmA:Rm*, and the disaggregation strain *gbfA*^−^ (Fig. 4). One common feature of these strains is reduced synchronicity during development. It is possible that the presence of un-aggregated cells contributed to the transcriptional profile such that the expression of the milestone genes appeared attenuated.

**Fig. 4.**
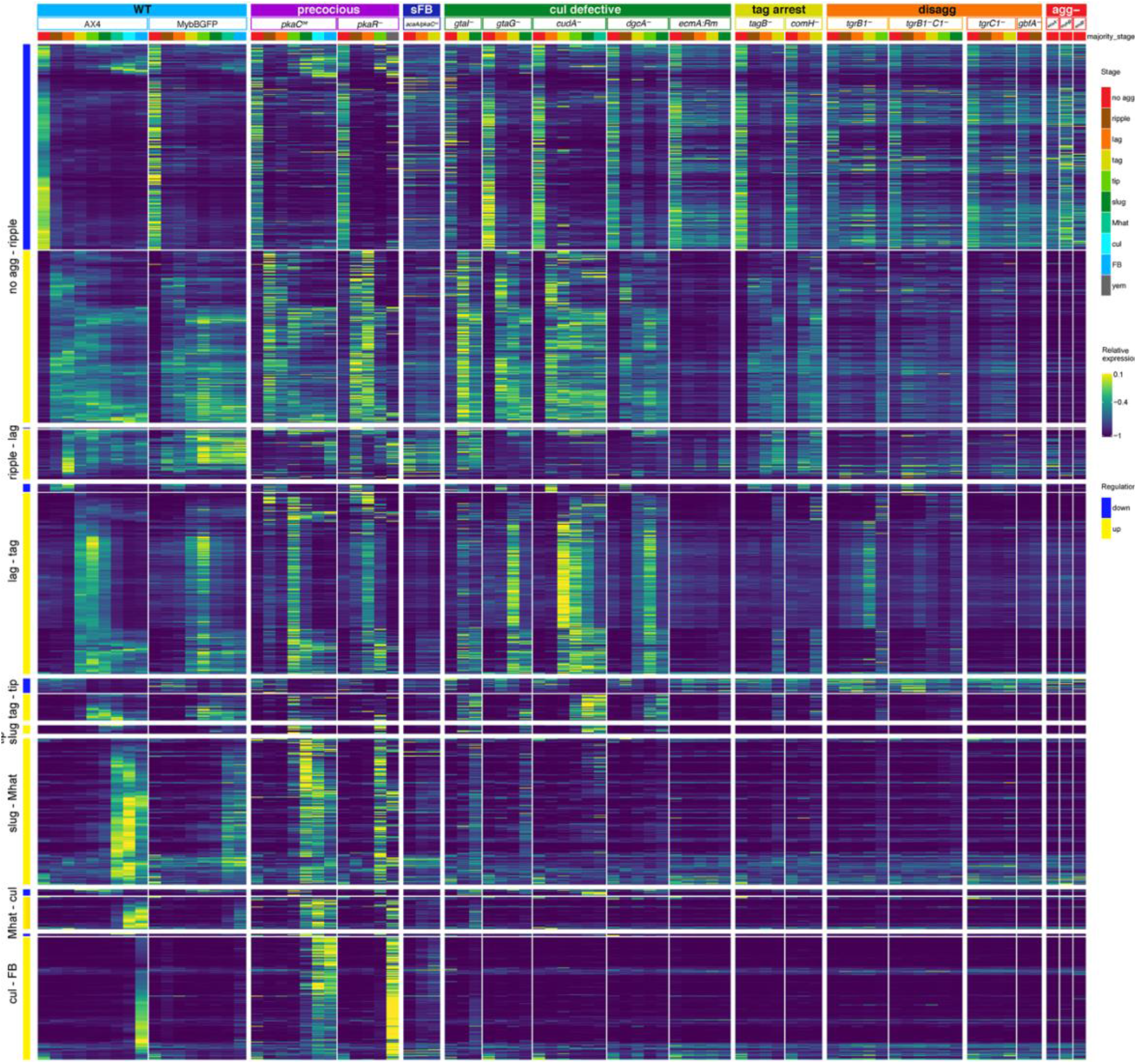
Transcriptional milestones mark developmental transitions. The heatmaps show the mRNA abundance in the indicated 19 strains, where each row represents a gene and each column represents expression data averaged across the multiple samples annotated with a common majority stage in the strain. The yellow-blue colors represent relative mRNA abundances (Relative expression scale on the right). Stage transitions are indicated on the left. The genes are ordered based on the direction of their regulation patterns (blue – down, yellow – up). Phenotype groups, strain names and annotated morphological stages are indicated above the heatmap blocks. The phenotype groups are: wild type (WT, light blue), precocious development (precocious, violet), small fruiting body (sFB, dark blue), culmination defective (cul defective, dark green), tight aggregate arrest (tag, dark yellow), tight aggregate / loose aggregate disaggregation (disagg, orange), and aggregationless (agg-, red). The morphological stage color represents the majority morphology as indicated in the legend on the right: no aggregation (no_agg, red), rippling/stream (ripple, brown), loose aggregate (lag, orange), tight aggregate (tag, dark yellow), tipped aggregate (tip, light green), slug/first finger (slug, dark green), Mexican hat (Mhat, jade green), culmination (cul, cyan), fruiting body (FB, light blue) and yellow mound (yem, dark gray).

The ripple-lag milestone genes were largely upregulated during the respective transitions except for the aggregationless and disaggregation strains, as expected. We observed an unexpected late induction in the precocious *pkaC^oe^* strain and lack of induction in the *gtaG^−^*, *cudA^−^*, and *dgcA*^−^ culmination-defective strains. The expression patterns in *pkaR^−^*, *ecmA:Rm*, and *acaA/pkaC^oe^* were also unexpected (Fig. 4), suggesting that expression of these milestone genes is affected by PKA.

The lag-tag milestone transcripts were largely upregulated during the tight aggregate stage in most strains and did not exhibit upregulation in the disaggregation and aggregationless mutants, which could not progress to the tight aggregate stage. Interestingly, however, many of the lag-tag milestone genes were not induced in the tight-aggregate arrest mutants (Fig. 4). Curiously, most of those genes were related to cell cycle progression (Supplemental File S1). This finding suggests that preparation for cell division is a key event in the progression from tight aggregates to the next stage.

The tag-tip, slug-Mhat, Mat-cul, and cul-FB milestone genes behaved largely as expected, failing to upregulate in the respective mutants (Fig. 4). The precocious mutants exhibited early rises in the accumulation of these transcripts, consistent with their accelerated morphogenesis. Overall, these findings indicate good matching between the transcriptional changes and the respective morphological transitions.

### Developmental regulons

The *D. discoideum* genome does not include large segments of co-regulated genes, like most other eukaryotic genomes (Parikh et al. 2010; Rosengarten et al. 2015). Nevertheless, many eukaryotic genes that have common functions are regulated by common transcriptional mechanisms. Regulons, which are groups of co-regulated genes, should be robust to various types of perturbation. Our dataset is ideal for regulon definition because it contains three major perturbations – development, which causes evolutionarily conserved changes in gene expression over time (Parikh et al. 2010); time, which affects the cells regardless of developmental progression; and mutations that perturb the developmental process and the transcriptome (Van Driessche et al. 2005). Therefore, we searched the dataset for groups of genes that remain co-regulated despite changes in development, time and genotype.

First, we considered individual strains. We searched for co-regulated gene pairs among the nearest neighbors by selecting transcripts that had at least one close neighbor that exhibited a similar expression pattern. Then, we considered individual transcripts by counting the number of strains in which the transcript was selected. Finally, we selected the top 1,099 transcripts based on the number of close neighbors in multiple strains. The leftmost section of Fig. 5 shows heatmaps of 13 clusters (letters, A-M) that were found by considering only the developmental perturbations in AX4. Transcripts in cluster A exhibit high abundance in vegetative cells and low abundance after starvation. Clusters B through L exhibit peak abundance at progressively later stages, and cluster M includes transcripts that peak at the terminal fruiting body stage.

**Fig. 5.**
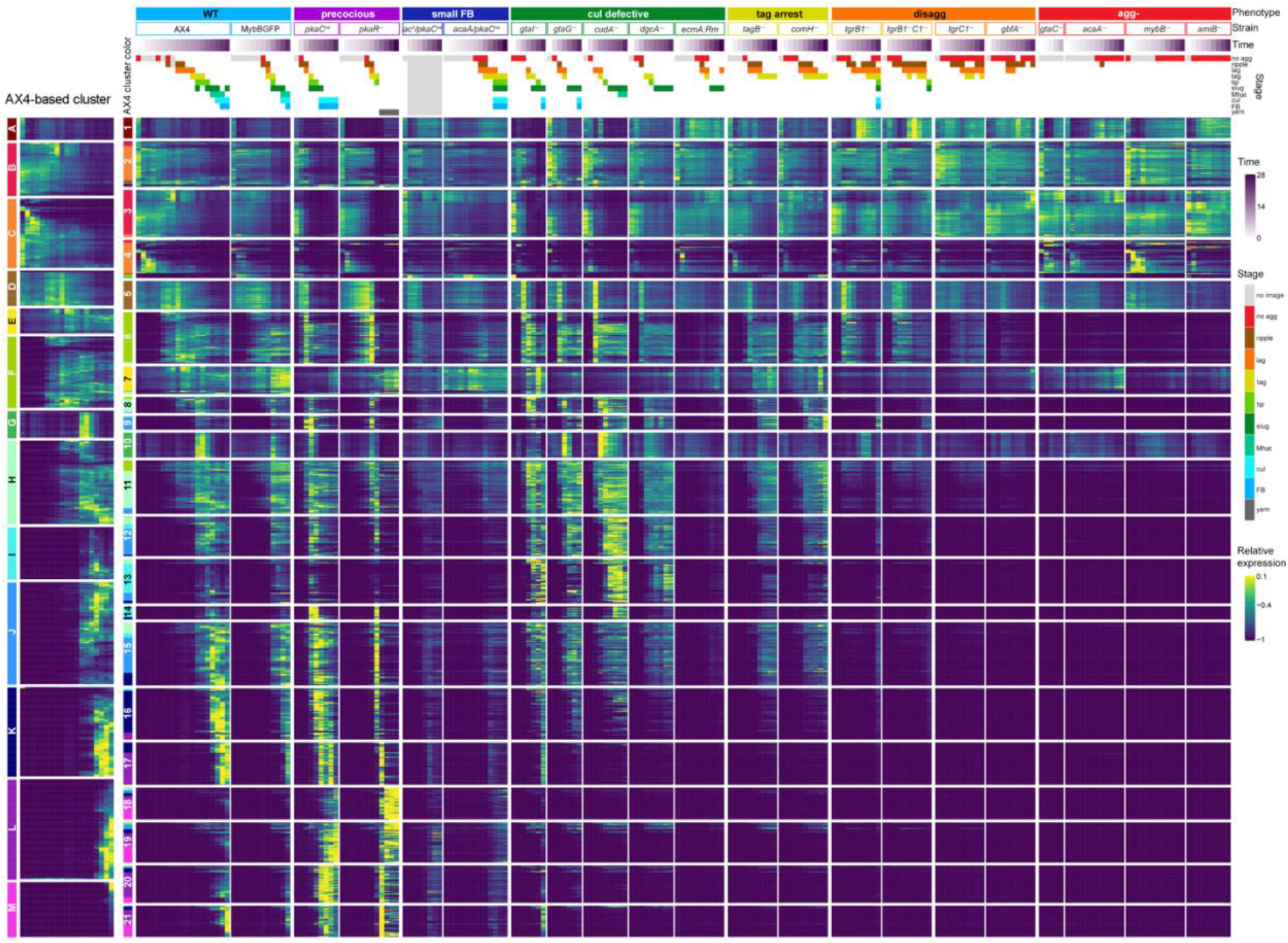
Developmental regulons are robust to temporal and genetic perturbations. We searched for genes whose expression patterns were similar despite changes in developmental time and strain genotype. The leftmost heatmap shows 13 clusters of genes that were co-expressed in the wild type AX4 during development. The clusters were assigned a letter (A to M) and a color as indicated on the left of the AX4-based column. We then re-clustered these genes based on their expression patterns in all 21 strains. The resulting 21 regulons (numbered boxes with AX4-based assignment colors, left) are shown as heatmaps. Each row in the heatmaps represents a gene, each column represents a time point and the entry colors represent relative mRNA abundances (as indicated in the relative expression scale on the right). Phenotype groups, strain names, developmental time (purple gradation, see scale on the right) and annotated morphological stages (see color legend on the right) are indicated above the heatmaps. The phenotype groups are: wild type (WT, light blue), precocious development (precocious, violet), small fruiting body (small FB, dark blue), culmination defective (cul defective, dark green), tight aggregate arrest (tag, dark yellow), tight aggregate / loose aggregate disaggregation (disagg, orange), and aggregationless (agg-, red). The morphological stages are: no aggregation (no agg, red), rippling/stream (ripple, brown), loose aggregate (lag, orange), tight aggregate (tag, dark yellow), tipped aggregate (tip, light green), slug/first finger (slug, dark green), Mexican hat (Mhat, jade green), culmination (cul, cyan), fruiting body (FB, light blue) and yellow mound (yem, dark gray). Light gray indicates that no image was captured (no image, light gray).

Considering all the other strains introduced genetic and temporal perturbations into the analysis, and refined the AX4 regulons into 21 clusters (numbered clusters, Fig. 5). Only three clusters remained largely unperturbed by the mutations; clusters 1, 7 and 10 are nearly identical to AX4 clusters A, E, and G, respectively. The most prevalent annotations in cluster 1 are related to ribosome biogenesis; cluster 7 includes cell death, RNA degradation, and morphogenesis annotations; and cluster 10 is enriched in cell division annotations (Supplemental File S2). AX4 cluster C was re-classified into clusters 2 and 4, mainly due to different expression patterns in the aggregationless strains (Fig. 5). The transcripts in cluster 2 were not down-regulated in the aggregationless strains, whereas the transcripts in cluster 4 were down-regulated, like in AX4. The gene-term enrichment analysis showed that oxidative phosphorylation was more enriched in cluster 2, whereas endocytosis and cytoskeletal protein binding were more enriched in cluster 4 (Supplemental File S2), suggesting that regulon refinement was aligned with improved classification. AX4 clusters H, I, and J were also re-classified into 2-3 clusters each, mainly clusters 8, 9, and 11-15, based on the differences in the culmination defective and precocious mutant strains (Fig. 5). Two annotations, *sig/sigN* genes and pst genes, were enriched in cluster 9 but not in clusters 12 and 15, all of which were in AX4 cluster J (Supplemental File S2). Comparison between clusters 13 and 14, which derive from AX4 cluster I, shows that the annotations *hssA/2C/7E* family and gtaG-dependent short proteins were enriched in both, but the annotations 57-aa protein family and pst gene were not enriched in cluster 14 (Supplemental File S2). Cluster L was broken into clusters 17, 20, and 21, whose expression patterns differed in the precocious strains (Fig. 5), and cluster M was distributed into clusters 18 and 19 due to the differences in the *pkaC*^oe^ strain (Fig. 5). Thus, by analyzing the data of all strains, we were able to obtain more detailed clusters. The enriched terms are detailed in Supplemental File S2. Overall, transcripts in any given cluster exhibited very similar expression patterns in AX4 and in the seven classes of mutants. The respective gene annotations indicated common or complementary functions in many cases. This finding suggests that regulons might have specific functions in the progression between developmental stages and each regulon is probably regulated by a unique combination of transcriptional mechanisms.

### Dedifferentiation signatures in the disaggregation mutant strains

The disaggregation strains exhibited reversal of morphogenesis (Fig. 1) and of transcriptional patterns, as their PC1 values oscillated over time (Fig. 2 and Supplemental Fig S4). These oscillations suggested that the cells have experienced cycles of differentiation and dedifferentiation. To examine this possibility, we searched the transcriptome dataset for disaggregation-related genes by performing differential expression analysis using *tgrB1*^−^ and *tgrB1^−^C1*^−^ as disaggregation representatives and AX4, *tagB^−^*, and *comH*^−^ as the non-disaggregation reference. The first peak of PC1 values of both *tgr* strains was around 8 h (Fig. 6A and Supplemental Fig S4). After that peak, the PC1 trendline regressed to a pattern similar to earlier developmental stages. We therefore selected transcripts that were differentially expressed before and after 8 h in both *tgr* strains and, separately, in each of the other strains. Then, we compared those transcripts and selected genes that were significantly upregulated at 6-8 h or 8-12 h in both *tgr* strains individually, but not in the other strains. The search yielded 72 transcripts at 6-8 h and 218 transcripts at 8-12 h. The heatmap in Fig. 6B shows the expression patterns of the 8-12 h set in all the strains, and Supplemental File S3 shows the gene-set enrichment analyses of both groups of transcripts. The selected transcripts were sharply up-regulated mainly in the *tgrB1*^−^ and *tgrB1^−^C1*^−^ strains at 8-12 h, while exhibiting low or constant levels in the other strains (Fig. 6B). The gene-enrichment analysis revealed lysine degradation, tryptophan metabolism, lipid binding, and membrane organization among the 6-8 h gene set (Supplemental File S3). The 8-12 h set was enriched in ribosome biogenesis and nucleolus annotations (Fig. 6C and Supplemental File S3). Many of these transcripts were also up-regulated during the pre-aggregation stage in the wild type (Fig. 6B), suggesting that the disaggregating cells may have returned to an earlier developmental stage.

**Fig. 6.**
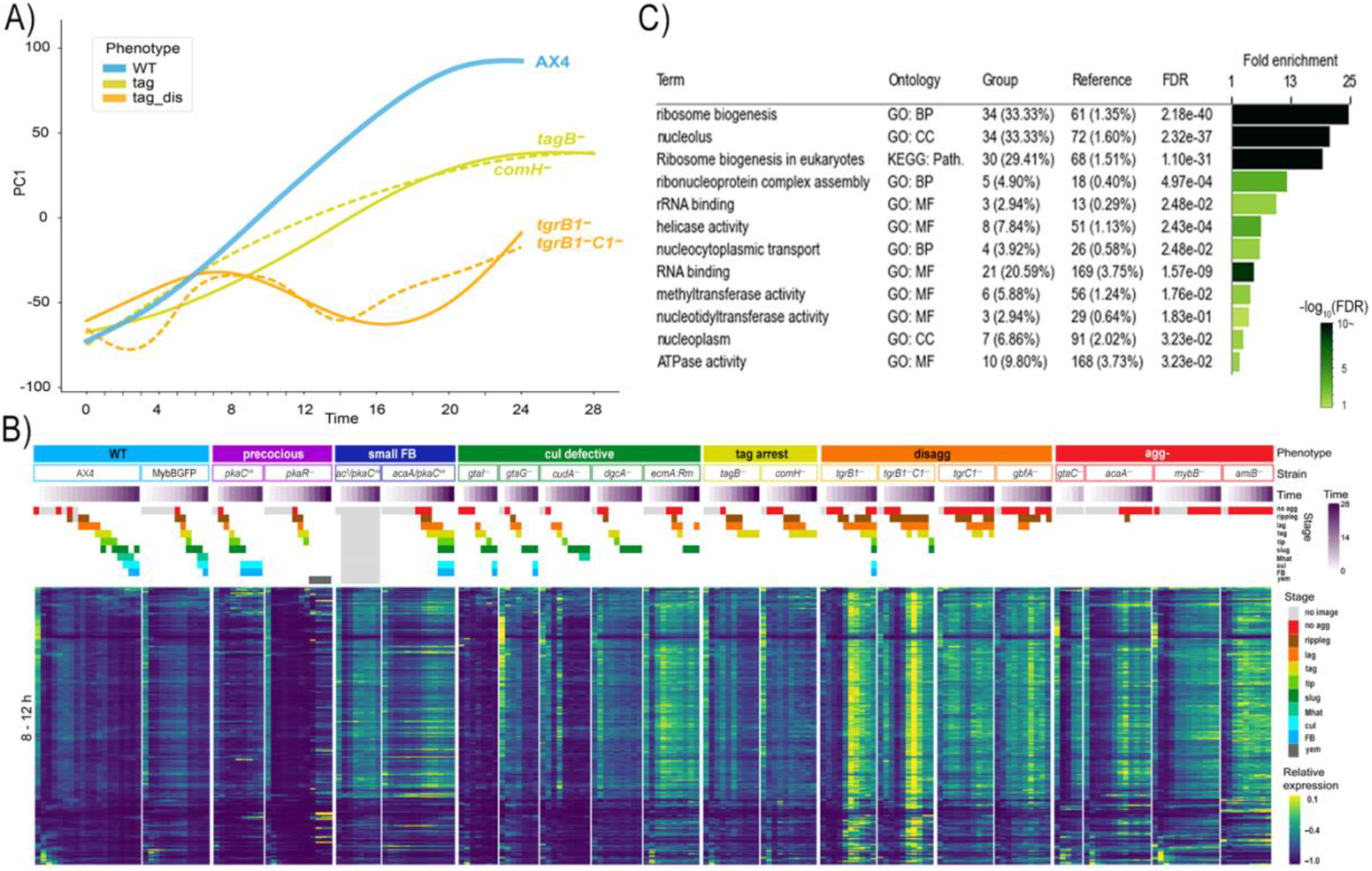
Transcriptome changes during disaggregation. We plotted PC1, which explains 30.4% of the variance, (y-axis, arbitrary units) against time (x-axis, hours) of the disaggregation strains (tag_dis, light orange), tag arrested strains (tag, dark yellow) and AX4 (WT, light blue). The graphs are a subset of Fig. 2. B) Up-regulated genes in both *tgrB1*^−^ and *tgrB1^−^C1*^−^ but not in *comH^−^*, *tagB*^−^ and AX4 were selected by differential expression analysis of the 8 to 12 h samples. The changes of mRNA abundance of these genes are shown as heatmaps in all 21 strains. Phenotype groups, strain names, developmental time (purple gradation, see scale on the right) and morphological stages (see color legend on the right) are indicated above the heatmaps. The phenotype groups are: wild type (WT, light blue), precocious development (precocious, violet), small fruiting body (small FB, dark blue), culmination defective (cul defective, dark green), tight aggregate arrest (tag, dark yellow), tight aggregate / loose aggregate disaggregation (disagg, orange), and aggregationless (agg-, red). The morphological stages are: no aggregation (no agg, red), rippling/stream (ripple, brown), loose aggregate (lag, orange), tight aggregate (tag, dark yellow), tipped aggregate (tip, light green), slug/first finger (slug, dark green), Mexican hat (Mhat, jade green), culmination (cul, cyan), fruiting body (FB, light blue) and yellow mound (yem, dark gray). C) The table shows gene-set enrichments among the up-regulated genes. The bar size shows the fold enrichment and the color (see scale) represents the FDR (false discovery rate, hypergeometric test). GO: Biological process (BP), Cellular component (CC), Molecular function (MF), KEGG: pathway (Path.)

Reversal of development in *D. discoideum* can occur through the orderly process of dedifferentiation (Katoh et al. 2004). We therefore compared the disaggregation transcriptome to a published dedifferentiation dataset (Nichols et al. 2020). We first preprocessed the published RNA-seq data to reduce pipeline effects and performed differential expression analysis, identifying 360 dedifferentiation-related transcripts that were enriched in several terms related to ribosomal biogenesis (Supplemental File S3). Moreover, the disaggregation and the dedifferentiation gene sets shared 70 genes, including most of the U3 small nucleolar ribonucleoprotein (snoRNP) genes that are involved in pre-rRNA processing (Supplemental File S3). Interestingly, all 30 genes in regulon 1 (Fig. 5) were included in the disaggregation gene set and 15 of them were included in the dedifferentiation set. Many of them are U3 snoRNPs and other rRNA processing-related genes. G-protein-coupled receptor genes were enriched in the disaggregation gene set as well, but not in the dedifferentiation gene set (Supplemental File S3). The overlap between the disaggregation and dedifferentiation gene sets was statistically significant (p-val: 9.25e-55, hypergeometric test) (Supplemental File S3), suggesting that the disaggregation phenotype includes dedifferentiation steps.

### Common gene-expression tools and standards

Several promoters are being used in *D. discoideum* expression vectors, mainly the actin promoters *act6* and *act15* (Knecht et al. 1986) and the recently adopted *coaA* promoter (Paschke et al. 2018). We tested the effects of genetic perturbation and temporal and developmental progression on the mRNA abundance of *act6*, *act15*, and *coaA* during development in our data set and in published data (Rosengarten et al. 2015), in which AX4 cells were starved in buffer suspension and subjected to cAMP pulses (Fig. 7A). We found that *act6* mRNA abundance was rather low in vegetative cells (0 h), higher during starvation in most strains, and very low after aggregation. The post-starvation increase was suppressed by cAMP-pulses and PKA-C overexpression. The *act6* mRNA abundance changed nearly 100-fold during AX4 filter development and exhibited more than 350-fold difference between the lowest level in the cAMP-pulse experiment and the highest level in AX4 filter development (Fig. 7A). The vegetative mRNA abundance of *act15* was a bit higher than *act6*, but it also varied widely during development and between strains. The *act15* mRNA abundance changed 10-fold during AX4 development, 5-fold in the *pkaC^OE^* mutant, and as much as 40-fold between the highest level in AX4 and the lowest level in *pkaC^OE^*. It was also generally reduced by cAMP-pulses and PKA-C overexpression and increased in the absence of cAMP in the *acaA*^−^ mutant (Fig. 7A). The mRNA abundance of *coaA* was 8-28 fold higher than the two actin mRNA abundance in vegetative cells, accounting for its improved utility in gene expression vectors, but it also exhibited great variability between strains and developmental stages. Moreover, *coaA* abundance exhibited as much as a 5-fold change during AX4 development and a 33-fold difference between the highest point in *pkaC^OE^* and the lowest point in *pkaR^−^*. Nevertheless, unlike the actin genes, *coaA* mRNA abundance was not suppressed by cAMP pulses or PKA-C overexpression (Fig. 7A).

**Fig. 7.**
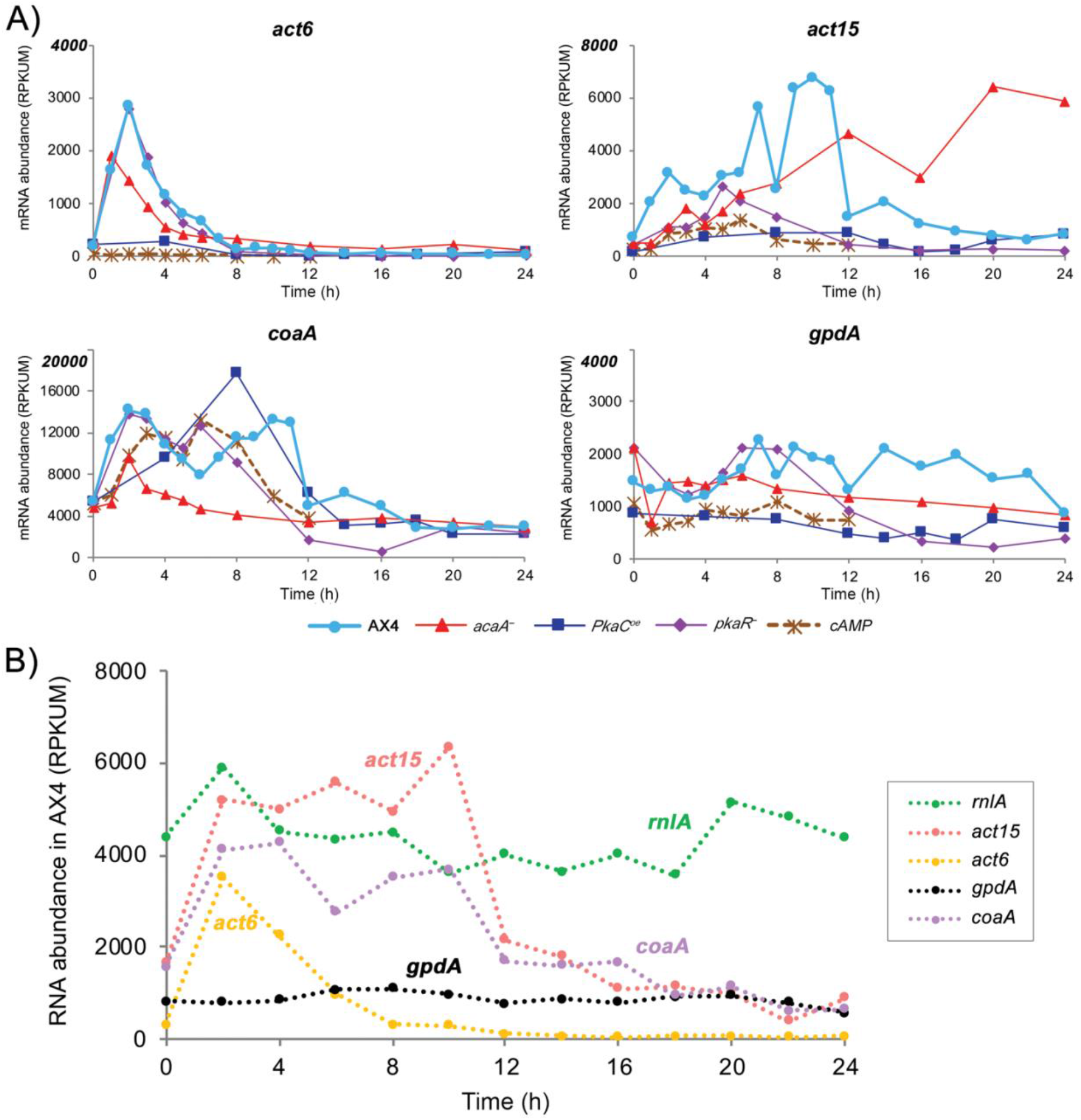
The expression patterns of common genetic markers. The mRNA abundance trajectories of the genes *act6*, *act15*, *coaA* and *gpdA* (y-axis, RPKUM) are plotted against developmental time (x-axis, hours). A) Data are from our data set (normal filter development) for strains AX4, *acaA^−^*, *pkaC^oe^* and *pkaR*^−^ and from AX4 cells developed in suspension with cAMP pulses (Rosengarten et al. 2015). The gene names are indicated above each panel and the strains/conditions are indicated below the panels (cAMP – cells pulsed with cAMP in suspension). Note that the y-axes are different between the panels. B) Data are from AX4-development RNA samples that were enriched by ribosomal RNA depletion (Rosengarten et al. 2017). Gene names are shown on the right.

Seeking a transcript that exhibits relatively high and constant mRNA abundance levels revealed *gpdA*, which encodes the *D. discoideum* glyceraldehyde-3-phosphate dehydrogenase (GAPDH; Fig. 7A). GAPDH is a common RNA quantification standard in other systems, whereas *rnlA* (1G7) has been a common standard in *D. discoideum* (Early and Williams 1988). *rnlA* is a mitochondrial ribosome RNA component, which is not polyadenylated, so it is not included in our dataset. We therefore compared *rnlA* expression to that of *gpdA* and the other genes in a published RNA-seq dataset that did not rely on enrichment of polyadenylated RNA (Rosengarten et al. 2017). Fig. 7B shows that *rnlA* RNA abundance was indeed higher than *gpdA*, but it fluctuated more during AX4 development.

## Discussion

We performed RNA-seq analysis on 20 developmental mutants and compared them to the wild type. We chose strains that exhibit similar developmental defects due to disruption of different pathways to increase the probability that the transcriptional changes would correlate with the morphological stage rather than the genetic manipulation. For example, the four aggregationless mutations affect different pathways but result in the same gross morphology. Some of the other mutations arrest development at typical wild-type stages, including pre-aggregation, tight aggregates, and fingers. Others exhibit precocious development, allowing distinction between the effects of time and morphology, and some exhibit morphologies not seen in the wild type, such as small fruiting bodies and disaggregation. Differences between the developmental conditions and analysis pipelines were minimal, to reduce experimental and computational effects on the outcome. Developmental morphology and transcriptome patterns change over time within each strain, and genetic perturbations cause changes between strains. These facts allowed us to evaluate the genetic effects by comparing data between strains and to evaluate the developmental/temporal effects by comparing data between time points within each strain. Most importantly, the genetic and temporal factors were considered to be distinct perturbations, which allowed us to test the relationship between morphology and the transcriptome.

Our data indicate coupling between morphogenesis and transcriptional changes. The most convincing evidence is the similarity between the transcriptomes of mutants that exhibit common morphologies, such as the aggregationless mutants, irrespective of time. The mutations that cause these phenotypes affect different pathways and there is no theoretical reason to assume that their transcriptomes should be similar, especially after prolonged incubation. Likewise, the differences between the mutant groups strongly indicate that time alone is not a major factor. In fact, the morphological stage is the most likely determinant in the similarity between transcriptional stages across samples, with a few notable exceptions. In the precocious *pkaR*^−^ mutant, gross morphology indicated an arrest at the tight aggregate stage whereas the transcriptome indicated progression to the wild-type equivalent of culmination. In this case, examination of fine morphology revealed that cell differentiation into spores and stalks was more related to the transcriptome than gross morphology. Another exception was observed in strains that overexpress the PKA catalytic subunit *pkaC* or the mutated regulatory subunit *pkaRm*. In those strains, the transcriptomes at the initial time points seemed shifted to the right along the x-axis and up along the y-axis in the MDS plots. This property is probably due to overexpression of the PKA subunits rather than any overexpression from the G418-selection vectors because it was not seen in the *mybB-GFP* strain, which also carries the G418-selection vector.

Previous studies of wild-type development indicated that the transcriptional changes occur in leaps and lulls – bursts of change separated by periods of relatively few changes. The annotations of the genes that underwent changes during these leaps were correlated well with known developmental processes (Rosengarten et al. 2015). The milestones we identified here are different because they include only sharp transcriptional transitions that occur at the boundary between two morphological stages. They were also visualized in the mutant strains, a step that was not included in the previous studies. Nevertheless, the overall picture that emerges from these milestones is similar to that found in the previous study (Rosengarten et al. 2015). The initial developmental milestones include down-regulation of metabolism pathways and up-regulation of signaling and cell morphology pathways. The finding that only the first milestone was accompanied by a large group of down-regulated genes does not mean that genes are not down-regulated at later stages. It only means that downregulation at later stages is not as sharp as at the onset of development. It is important to notice that RNA-seq measures mRNA abundance, which is a function of RNA synthesis and stability. Therefore, it is possible that mRNA degradation is slower or less effective at later stages of development.

Another significant milestone includes the simultaneous induction of cell division and DNA replication genes during the loose-aggregate to tight-aggregate transition, which is the subject of some controversy. Early studies showed that *D. discoideum* develops well without a significant change in cell number (Sussman and Sussman 1960), but later studies suggested that prespore cells replicate DNA and divide during development (Zimmerman and Weijer 1993). Subsequent studies, using molecular resolution techniques, showed that chromosomal DNA is not replicated during development. Instead, the cells replicate their mitochondrial DNA and undergo mitosis, reducing their chromosomal DNA content from a G2-phase equivalent to a G1-phase equivalent (Shaulsky and Loomis 1995; Chen et al. 2004). More recently, live imaging techniques were used to argue again in favor of cell division and DNA replication during development (Muramoto and Chubb 2008), but these studies did not explain how replication might occur without incorporation of nucleotides into the chromosomal DNA. Although the problem has not yet been resolved, we may have found a possible explanation for the controversy. Our data show that cell division and DNA replication genes are sharply and contemporaneously induced during development, which confirms and extends previous findings that used different mutants (Strasser et al. 2012). In addition, we observed that the tag-arrest strains do not exhibit elevated expression of these genes, even though they pass the respective morphological transition. Moreover, the *pkaR*^−^ mutant, which exhibits tag arrest but proceeds to complete cell differentiation, exhibits up-regulated expression of these genes as well. Altogether, these findings suggest the existence of a mechanism that co-regulates cell division and DNA replication genes in *Dictyostelium*, unlike budding yeast (Haase and Wittenberg 2014). During vegetative growth, this co-regulation couples the processes of cell division and DNA replication, possibly due to the lack of a G1-phase in the *Dictyostelium* cell cycle (Strasser et al. 2012). During development, however, cell division is probably essential for progression past the tight aggregate stage. Induction of the cell division genes causes an incidental co-induction of the DNA replication genes even though chromosomal DNA replication does not follow (Shaulsky and Loomis 1995; Chen et al. 2004).

The notable absence of prespore genes from the lag-tag milestone is consistent with the fact that these genes continue to accumulate during later stages of development. The tag-tip milestone includes down-regulation of cytoskeleton and cell-adhesion genes, consistent with the major changes in tissue organization that accompany cell-type sorting and tip formation. The tip-slug transition is low in milestone genes, which is surprising considering the major changes that must accompany that transition. This transition is either accompanied by more gradual changes that did not satisfy the milestone definition, or by earlier changes that set the stage for the transition. Subsequent milestones include late prespore gene induction at the slug-Mhat transition, and sugar metabolism and morphogenesis genes at the cul-FB transition. These transcriptional milestones were concordant with the mutation data in that most of them did not occur in mutants that failed to undergo the respective morphological transitions.

Genes that function in common cellular processes or structures are often clustered in operons in prokaryotes and in a few eukaryotes (Blumenthal 2004), but not in *D. discoideum* (Eichinger et al. 2005). Nevertheless, the *D. discoideum* genome includes regulons, which are groups of co-expressed genes that are useful predictors of function (Booth et al. 2005). Our dataset revealed many groups of genes that remained co-regulated despite genetic and temporal/developmental perturbations. Many of the regulons encode components of common protein machines (Alberts 1998), including ribosome structure, ribosome assembly, oxidative phosphorylation, proteasome, cytoskeleton elements, and cytoskeleton assembly. These findings validate our approach and suggest that regulons could be used to predict gene function. Indeed, 640 of the 1,099 regulon genes are either not annotated or partly annotated in dictyBase (Basu et al. 2013; Fey et al. 2019). Deeper examination of a few examples suggests that the regulons could predict gene function rather accurately. Regulon 1 (Fig. 5) contains 30 genes, 6 of which are unnamed and 12 of which are annotated as ribosome biosynthesis genes. One gene, DDB_G0271272, is annotated in dictyBase as being similar to yeast pol5 and human mybBP1A-binding protein. Based on the regulon annotation, DDB_G0271272 is probably involved in rRNA transcription, similar to the yeast homolog. Likewise, regulon 5 (Fig. 5) contains 41 genes, 3 of which are unnamed and 29 of which are annotated as proteasome subunit genes. One gene, DDB_G0291003, is automatically annotated in dictyBase as a putative protein DOA1, which plays a role in protein ubiquitination, sorting, and degradation in other organisms, a good fit with the proteasome annotation.

The similarity between the published dedifferentiation patterns (Nichols et al. 2020) and the disaggregation mutants illustrates the utility of our dataset for discovery and comparison across experiments. It also shows that the disaggregation process that results from lack of proper TgrB1-TgrC1 signaling is a dedifferentiation process. This finding also supports the hypothesis that TgrB1-TgrC1 signaling is a developmental checkpoint (Dynes et al. 1994; Benabentos et al. 2009; Hirose et al. 2015) and suggests that failure to pass this checkpoint leads to cycles of dedifferentiation and re-differentiation.

Our dataset has a few other practical implications. Firstly, the choice of promoters for ectopic gene expression in *D. discoideum* should be considered carefully. We recommend using *coaA*-based vectors (Paschke et al. 2018) rather than *act6* or *act15*-based vectors (Knecht et al. 1986; Manstein et al. 1995; Veltman et al. 2009). The transition to *coaA*-based vectors could be greatly facilitated by GoldenBraid (Kundert et al. 2020). In some cases, it may be worthwhile to reevaluate conclusions from experiments that involved gene expression under actin promoters, especially if PKA and cAMP signaling were involved. Secondly, we suggest using *gpdA* as a standard for RNA quantification, especially if poly-A selection of mRNA is used. The commonly used *rnlA* (1G7) (Early and Williams 1988) is not poly-adenylated in *D. discoideum*, but it comes through poly-A purifications as a contaminant because of its high abundance, which makes it unsuitable as a standard.

The data presented here represent a large collection of transcriptomes that can be used by the research community for further exploration. We suggest two convenient options in addition to downloading the data from the public repository and analyzing them in-house. The simplest way is to view and analyze the data on dictyExpress, which is a web-based platform for data exploration and comparison with other published datasets (Stajdohar et al. 2017) (Supplemental File S4a). This platform does not require any programming skills and it is conveniently linked to dictyBase (Fey et al. 2019). Data analysis can be extended by using the bioinformatics component in Orange, a visual programming system that is also linked to dictyExpress and dictyBase and facilitates data mining without scripting skills (Demsar et al. 2013) (Supplemental File S4b). The data can also be used to identify and test gene regulatory elements such as promoters and transcription factors, compare a gene of interest to genes in the regulon and milestone lists, and help in annotating poorly characterized genes. We recommend comparing future RNA-seq datasets to the ones presented here.

## Methods

### Cell culture, strain maintenance, development and spore collection

All the *Dictyostelium discoideum* strains were derivatives of AX4 (Knecht et al. 1986) as detailed in Supplemental Table S1. Growth and developmental conditions are described in Supplemental File S6.

### RNA-seq

We collected the cells from one nitrocellulose filter at each time point of two to seven independent developmental series, extracted total RNA and performed poly(A) selection twice as described (Katoh-Kurasawa et al. 2016). cDNA library multiplexing and mapping were done as described (Miranda et al. 2013). The data were deposited in GEO (accession numbers GSE152851). Additional information is provided in Supplemental File S6.

### Multi-dimensional scaling and principal component analysis

Dimensionality reduction with multi-dimensional scaling (MDS) was performed on the averaged RPKUM data across independent replicates at each developmental time point in 21 strains. About 3% of all protein-coding genes (12,828) were not expressed under any of the conditions sampled in all 21 strains. Using MDS (R-function: cmdscale), we visualized relative distances between each whole-transcriptome pair among all samples (Santhanam et al. 2015) in 2-dimensional space. We used all RPKUM data and Spearman’s correlation (SC) to calculate the distance (D = 1-SC).

We performed PCA on the preprocessed AX4 RPKUM data, except for genes non-expressed at any developmental time point in AX4. The PC1 embedding was obtained with the scikit-learn (v0.22.2) Python library (Pedregosa et al. 2011). The preprocessed data of other strains were transformed with the AX4-based embedding. For each strain, a generalized additive model (GAM) was fit to PC1 embedded samples across time with the pygam (v0.8.0) Python library (Servén and Brummitt 2018). Parameters of GAM (number of splines and smoothing parameter) were selected for each strain with grid search using ‘leave one timepoint out’ cross-validation. The prediction quality was compared with mean-squared error.

### Selection of milestone genes

To find genes with variable expression trajectories across developmental stages, we used AX4 samples with developmental stage annotations based on morphological analysis of images taken at different developmental times. When the image contained multiple developmental stages, the majority morphology was chosen as the stage annotation. Then, using the stage annotation, we performed differential expression analyses with the DESeq2 (v1.26.0) R library (Love et al. 2014) and ImpulseDE2 (v1.10.0) R library (Fischer et al. 2018) as a combinational approach. Milestone genes were defined according to two criteria; 1) the gene was significantly differentially expressed between two stages based on DESeq2 and 2) it was significantly DE across stages based on ImpulseDE2 with assigned transition time between the two stages. The DESeq2 analysis was performed for every pair of neighboring stages using the AX4 samples in case–control mode. The later stage was used as the case and the earlier as the control. The padj was recalculated over all the tests for all neighboring stage pairs using Benjamini-Hochberg correction on the pval from DESeq2. We then used absolute lFC ≥ 2 and padj ≤ 0.01 to select differentially expressed genes and classified them as up- or down-regulated. The ImpulseDE2 model was used to fit AX4 data, using stage annotations converted to consecutive integers in developmental order as time-points. We ran the analysis in ‘case-only’ mode to identify genes whose trajectories change at stage boundaries once or twice during development, using padj threshold = 0.001. The parameters of the fitted models were used to obtain neighboring stages when the transitions of the gene profile occurred. The genes significantly differentially expressed across neighboring stages were defined as milestone genes. Heatmaps were made with the ComplexHeatmap package (Gu et al. 2016), with expression data averaged across multiple samples that were annotated as the same stage in each strain, except for the two strains whose images were not captured. Additional details of the computational analysis are provided in Supplemental File S5 (Computational methods). A complete list of the milestone genes is available in Supplemental File S7.

### Regulon extraction

We extracted co-regulated gene pairs as regulon candidates in individual strains to avoid bias toward strains with more samples. We preprocessed RPKUM data of the genes whose expression was non-NULL in a strain and used the k-nearest neighbors (kNN) descent algorithm, PyNNDescent (v0.3.3) with cosine similarity (Dong et al. 2011), to obtain the top 300 nearest neighbors of each gene. We counted the number of strains in which the gene had at least one nearest neighbor above the strain-specific threshold (the 30^th^ percentile of the similarities to the closest neighbors in the strain), and we also specified a gene-specific N threshold (capped at 18) by the number of strains in which the gene was expressed. When the gene’s expression at any time points in a strain reached 10% of the 99^th^ percentile of the expression in all samples, we determined that the gene was expressed highly enough in the strain. If the gene had one nearest neighbor in at least N strains, we kept it as a regulon candidate.

Using the strain-specific and the gene-specific thresholds, we selected 1,099 regulon candidate genes and assigned them into regulons with Louvain clustering analysis on preprocessed expression data with data mining software Orange (v3.26) (Demsar et al. 2013). Regulon clustering was performed on the AX4 expression data or all strains data. Additional details are provided in Supplemental File S5 (Computational methods). A complete list of the regulon genes is available in Supplemental File S7.

### Analysis of disaggregation genes

To select genes that are related to the disaggregation process in the *tgr* strains, we extracted genes upregulated around the first peak-time of PC1 values in *tgrB1*^−^ and *tgrB1^−^C1^−^*, but not in AX4, *tagB*^−^ or *comH^−^*. Differentially expressed genes between pairs of time points 6 and 8 h, and 8 and 12 h in each strain were extracted with DESeq2. We then subtracted the genes that were upregulated for AX4, *tagB*^−^ or *comH*^−^ strains from the genes that were upregulated for *tgrB1*^−^ and *tgrB1^−^C1*^−^ strains. The DESeq2 results were optimized for padj threshold 0.01. A gene was considered upregulated if lFC ≥ 1.32 and padj ≤ 0.01. A complete list of the disaggregation genes is available in Supplemental File S7.

To characterize the selected disaggregation genes, we compared disaggregation genes with the dedifferentiation genes which are upregulated during early dedifferentiation process. The dedifferentiation genes were obtained from published dedifferentiation RNA-seq data (Nichols et al. 2020). The published data included an experiment, in which cells were disaggregated and incubated in nutrient medium to induce dedifferentiation, and a control in which the disaggregated cells were incubated in non-nutrient buffer. We downloaded the published RNA-seq data (GSE144892) and prepared their RPKUM data by the same procedure as ours. Dedifferentiation genes were selected based on a DESeq2 comparison between the ‘dedifferentiation’ samples at 0.5, 1, and 2 h and the ‘control’ samples at 0, 0.5, 1, 2, 3, 4, and 6 h. The DESeq2 results were optimized for padj threshold 0.01. A gene was considered to be upregulated during early dedifferentiation if lFC ≥ 2 and padj ≤ 0.01. Gene expression scaling and gene ordering for the heatmaps were performed as above. We used a hypergeometric test to determine whether the disaggregation genes significantly overlap with the dedifferentiation genes.

### Gene-set enrichment analysis

We performed Gene-set enrichment analysis using the Orange Bioinformatics (v4.0.0) Python library (Demsar et al. 2013). We used Generic GO-slims, KEGG Pathways, and custom gene sets (Supplemental File S8) with sizes between 5 and 500 genes as functional ontology terms. We used all protein-coding genes, except those that had all zero RPKUM values in our samples, as the reference set (‘Reference’) and filtered out genes without Entrez ID from the ‘Reference’ and query group (‘Group’). To account for different proportions of genes annotated with a gene set between the reference and the query groups we used only genes annotated with at least one term. We compared the frequency of individual annotations in the ‘Group’ list with that of the ‘Reference’ list to calculate individual fold enrichments. Enrichment was calculated using hypergeometric test with Benjamini-Hochberg correction. The results were filtered to display only ‘Terms’ with ‘FDR’ ≤ 0.25 and overlap with ‘Group’ ≥ 2. Additional details are provided in Supplemental File S5 (Computational methods).

## Data Access

All raw and processed sequencing data generated in this study have been submitted to the NCBI Gene Expression Omnibus (GEO; https://www.ncbi.nlm.nih.gov/geo/) under accession number GSE152851.

## Acknowledgements

We thank Adam Kuspa for insightful discussions. This work was supported by grant number R35 GM118016 from the National Institutes of Health. Bulk RNA-Seq was performed at the Department of Molecular and Human Genetics Functional Genomics Core at Baylor College of Medicine, partially supported by an NIH shared instrument grant S10OD023469 to Rui Chen.

## References

Abe K, Yanagisawa K. 1983. A new class of rapidly developing mutants in Dictyostelium discoideum: Implications for cyclic AMP metabolism and cell differentiation. Dev Biol 95: 200–210.

Alberts B. 1998. The cell as a collection of protein machines: preparing the next generation of molecular biologists. Cell 92: 291–294.

Basu S, Fey P, Pandit Y, Dodson R, Kibbe WA, Chisholm RL. 2013. DictyBase 2013: integrating multiple Dictyostelid species. Nucleic Acids Res 41: D676–683.

Benabentos R, Hirose S, Sucgang R, Curk T, Katoh M, Ostrowski EA, Strassmann JE, Queller DC, Zupan B, Shaulsky G et al. 2009. Polymorphic members of the lag gene family mediate kin discrimination in Dictyostelium. Current biology : CB 19: 567–572.

Blumenthal T. 2004. Operons in eukaryotes. Brief Funct Genomic Proteomic 3: 199–211.

Booth EO, Van Driessche N, Zhuchenko O, Kuspa A, Shaulsky G. 2005. Microarray phenotyping in Dictyostelium reveals a regulon of chemotaxis genes. Bioinformatics 21: 4371–4377.

Cai H, Katoh-Kurasawa M, Muramoto T, Santhanam B, Long Y, Li L, Ueda M, Iglesias PA, Shaulsky G, Devreotes PN. 2014. Nucleocytoplasmic shuttling of a GATA transcription factor functions as a development timer. Science 343: 1249531.

Chen G, Shaulsky G, Kuspa A. 2004. Tissue-specific G1-phase cell-cycle arrest prior to terminal differentiation in Dictyostelium. Development 131: 2619–2630.

Demsar J, Curk T, Erjavec A, Gorup C, Hocevar T, Milutinovic M, Mozina M, Polajnar M, Toplak M, Staric A et al. 2013. Orange: Data Mining Toolbox in Python. Journal of Machine Learning Research 14: 2349–2353.

Dong W, Moses C, Li K. 2011. Efficient k-nearest neighbor graph construction for generic similarity measures. In Proceedings of the 20th international conference on World wide web, doi:10.1145/1963405.1963487, pp. 577–586. Association for Computing Machinery, Hyderabad, India.

Dynes JL, Clark AM, Shaulsky G, Kuspa A, Loomis WF, Firtel RA. 1994. LagC is required for cell-cell interactions that are essential for cell-type differentiation in Dictyostelium. Genes & development 8: 948–958.

Early VE, Williams JG. 1988. A Dictyostelium prespore-specific gene is transcriptionally repressed by DIF in vitro. Development 103: 519–524.

Eichinger L, Pachebat JA, Glockner G, Rajandream MA, Sucgang R, Berriman M, Song J, Olsen R, Szafranski K, Xu Q et al. 2005. The genome of the social amoeba Dictyostelium discoideum. Nature 435: 43–57.

Fey P, Dodson RJ, Basu S, Hartline EC, Chisholm RL. 2019. dictyBase and the Dicty Stock Center (version 2.0) - a progress report. Int J Dev Biol 63: 563–572.

Fischer DS, Theis FJ, Yosef N. 2018. Impulse model-based differential expression analysis of time course sequencing data. Nucleic Acids Res 46: e119.

Glockner G, Lawal HM, Felder M, Singh R, Singer G, Weijer CJ, Schaap P. 2016. The multicellularity genes of dictyostelid social amoebas. Nat Commun 7: 12085.

Gomer RH, Jang W, Brazill D. 2011. Cell density sensing and size determination. Development, growth & differentiation 53: 482–494.

Good JR, Cabral M, Sharma S, Yang J, Van Driessche N, Shaw CA, Shaulsky G, Kuspa A. 2003. TagA, a putative serine protease/ABC transporter of Dictyostelium that is required for cell fate determination at the onset of development. Development 130: 2953–2965.

Gu Z, Eils R, Schlesner M. 2016. Complex heatmaps reveal patterns and correlations in multidimensional genomic data. Bioinformatics 32: 2847–2849.

Haase SB, Wittenberg C. 2014. Topology and control of the cell-cycle-regulated transcriptional circuitry. Genetics 196: 65–90.

Hirose S, Santhanam B, Katoh-Kurosawa M, Shaulsky G, Kuspa A. 2015. Allorecognition, via TgrB1 and TgrC1, mediates the transition from unicellularity to multicellularity in the social amoeba Dictyostelium discoideum. Development 142: 3561–3570.

Katoh M, Shaw C, Xu Q, Van Driessche N, Morio T, Kuwayama H, Obara S, Urushihara H, Tanaka Y, Shaulsky G. 2004. An orderly retreat: Dedifferentiation is a regulated process. Proceedings of the National Academy of Sciences of the United States of America 101: 7005–7010.

Katoh-Kurasawa M, Santhanam B, Shaulsky G. 2016. The GATA transcription factor gene gtaG is required for terminal differentiation in Dictyostelium. J Cell Sci 129: 1722–1733.

Kessin RH. 2001. Dictyostelium - Evolution, cell biology, and the development of multicellularity. Cambridge Univ. Press, Cambridge, UK.

Knecht DA, Cohen SM, Loomis WF, Lodish HF. 1986. Developmental regulation of Dictyostelium discoideum actin gene fusions carried on low-copy and high-copy transformation vectors. Mol Cell Biol 6: 3973–3983.

Kundert P, Sarrion-Perdigones A, Gonzalez Y, Katoh-Kurasawa M, Hirose S, Lehmann P, Venken KJT, Shaulsky G. 2020. A GoldenBraid cloning system for synthetic biology in social amoebae. Nucleic Acids Res 48: 4139–4146.

Kuspa A. 2006. Restriction enzyme-mediated integration (REMI) mutagenesis. Methods in molecular biology 346: 201–209.

Loomis WF. 1975. Dictyostelium discoideum. A developmental system. Ac. Press, New York.

Loomis WF. 1978. The number of developmental genes in Dictyostelium. Birth Defects: Original Article Series 14 no.2: 497–505.

Loomis WF. 1998. Role of PKA in the timing of developmental events in Dictyostelium cells. Microbiol Mol Biol Rev 62: 684.

Love MI, Huber W, Anders S. 2014. Moderated estimation of fold change and dispersion for RNA-seq data with DESeq2. Genome biology 15: 550.

Manstein DJ, Schuster HP, Morandini P, Hunt DM. 1995. Cloning vectors for the production of proteins in Dictyostelium discoideum. Gene 162: 129–134.

Miranda ER, Rot G, Toplak M, Santhanam B, Curk T, Shaulsky G, Zupan B. 2013. Transcriptional profiling of Dictyostelium with RNA sequencing. Methods in molecular biology 983: 139–171.

Muramoto T, Chubb JR. 2008. Live imaging of the Dictyostelium cell cycle reveals widespread S phase during development, a G2 bias in spore differentiation and a premitotic checkpoint. Development 135: 1647–1657.

Nichols JM, Antolovic V, Reich JD, Brameyer S, Paschke P, Chubb JR. 2020. Cell and molecular transitions during efficient dedifferentiation. Elife 9.

Parikh A, Miranda ER, Katoh-Kurasawa M, Fuller D, Rot G, Zagar L, Curk T, Sucgang R, Chen R, Zupan B et al. 2010. Conserved developmental transcriptomes in evolutionarily divergent species. Genome biology 11: R35.

Paschke P, Knecht DA, Silale A, Traynor D, Williams TD, Thomason PA, Insall RH, Chubb JR, Kay RR, Veltman DM. 2018. Rapid and efficient genetic engineering of both wild type and axenic strains of Dictyostelium discoideum. PLoS One 13: e0196809.

Pedregosa F, Varoquaux G, Gramfort A, Michel V, Thirion B, Grisel O, Blondel M, Prettenhofer P, Weiss R, Dubourg V et al. 2011. Scikit-learn: Machine Learning in Python. J Mach Learn Res 12: 2825–2830.

Ritchie AV, van Es S, Fouquet C, Schaap P. 2008. From drought sensing to developmental control: evolution of cyclic AMP signaling in social amoebas. Mol Biol Evol 25: 2109–2118.

Rosengarten RD, Santhanam B, Fuller D, Katoh-Kurasawa M, Loomis WF, Zupan B, Shaulsky G. 2015. Leaps and lulls in the developmental transcriptome of Dictyostelium discoideum. BMC genomics 16: 294.

Rosengarten RD, Santhanam B, Kokosar J, Shaulsky G. 2017. The Long Noncoding RNA Transcriptome of Dictyostelium discoideum Development. G3 (Bethesda) 7: 387–398.

Rubin M, Miller AD, Katoh-Kurasawa M, Dinh C, Kuspa A, Shaulsky G. 2019. Cooperative predation in the social amoebae Dictyostelium discoideum. PLoS One 14: e0209438.

Santhanam B, Cai H, Devreotes PN, Shaulsky G, Katoh-Kurasawa M. 2015. The GATA transcription factor GtaC regulates early developmental gene expression dynamics in Dictyostelium. Nat Commun 6: 7551.

Schilde C, Lawal HM, Noegel AA, Eichinger L, Schaap P, Glockner G. 2016. A set of genes conserved in sequence and expression traces back the establishment of multicellularity in social amoebae. BMC genomics 17: 871.

Servén D, Brummitt C. 2018. pyGAM: Generalized Additive Models in Python. DOI: 105281/ZENODO1208724 doi:10.5281/ZENODO.1208724.

Shaulsky G, Loomis WF. 1995. Mitochondrial DNA replication but no nuclear DNA replication during development of Dictyostelium. Proceedings of the National Academy of Sciences of the United States of America 92: 5660–5663.

Stajdohar M, Rosengarten RD, Kokosar J, Jeran L, Blenkus D, Shaulsky G, Zupan B. 2017. dictyExpress: a web-based platform for sequence data management and analytics in Dictyostelium and beyond. BMC Bioinformatics 18: 291.

Strasser K, Bloomfield G, MacWilliams A, Ceccarelli A, MacWilliams H, Tsang A. 2012. A retinoblastoma orthologue is a major regulator of S-phase, mitotic, and developmental gene expression in Dictyostelium. PLoS One 7: e39914.

Sussman M. 1952. An analysis of the aggregation stage in the development of the slime molds, Dictyosteliaceae. II. Aggregative center formation by mixtures of Dictyostelium discoideum wild type and aggregateless variants. Biol Bull 103: 446–457.

Sussman M, Sussman RR. 1962. Ploidal inheritance in Dictyostelium discoideum: stable haploid, stable diploid and metastable strains. J Gen Microbiol 28: 417–429.

Sussman RR, Sussman M. 1960. The dissociation of morphogenesis from cell division in the cellular slime mould, Dictyostelium discoideum. J Gen Microbiol 23: 287–293.

Van Driessche N, Demsar J, Booth EO, Hill P, Juvan P, Zupan B, Kuspa A, Shaulsky G. 2005. Epistasis analysis with global transcriptional phenotypes. Nature genetics 37: 471–477.

Veltman DM, Akar G, Bosgraaf L, Van Haastert PJ. 2009. A new set of small, extrachromosomal expression vectors for Dictyostelium discoideum. Plasmid 61: 110–118.

Zimmerman W, Weijer CJ. 1993. Analysis of cell cycle progression during the development of Dictyostelium and its relationship to differentiation. Dev Biol 160: 178–185.

