## Supplemental Materials for "Transcriptional milestones in *Dictyostelium* development"

#### Supplemental Figures

##### Supplemental_Fig_S1 Precocious culmination in the pkaC-overexpressor strain

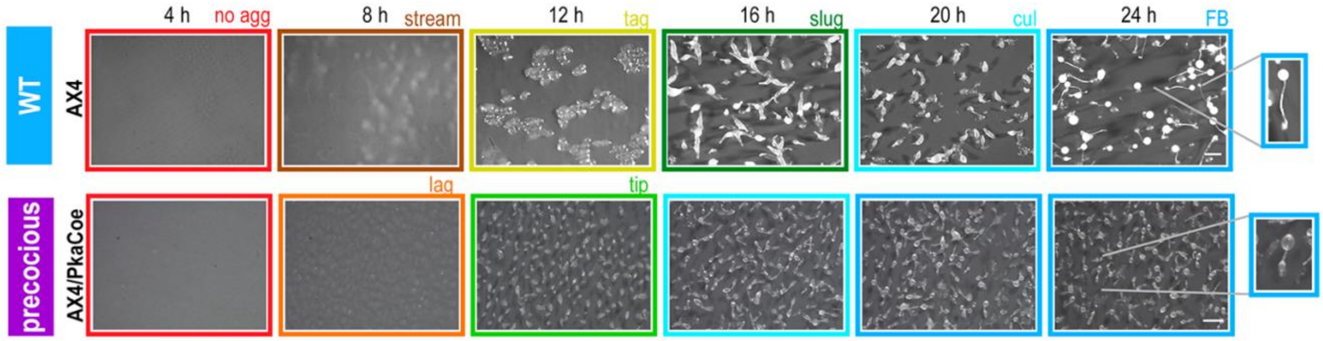

**Supplemental_Fig_S1 Precocious culmination in the pkaC-overexpressor strain.** Comparison between the wild type (AX4) and the *pkaC*$^{oe}$ developmental morphologies. The AX4 images, all the experimental details, and the color codes are identical to those shown in Fig. 1 in the main text.

Supplemental_Fig_S2 MDS and PCA plots of individual strains

A.

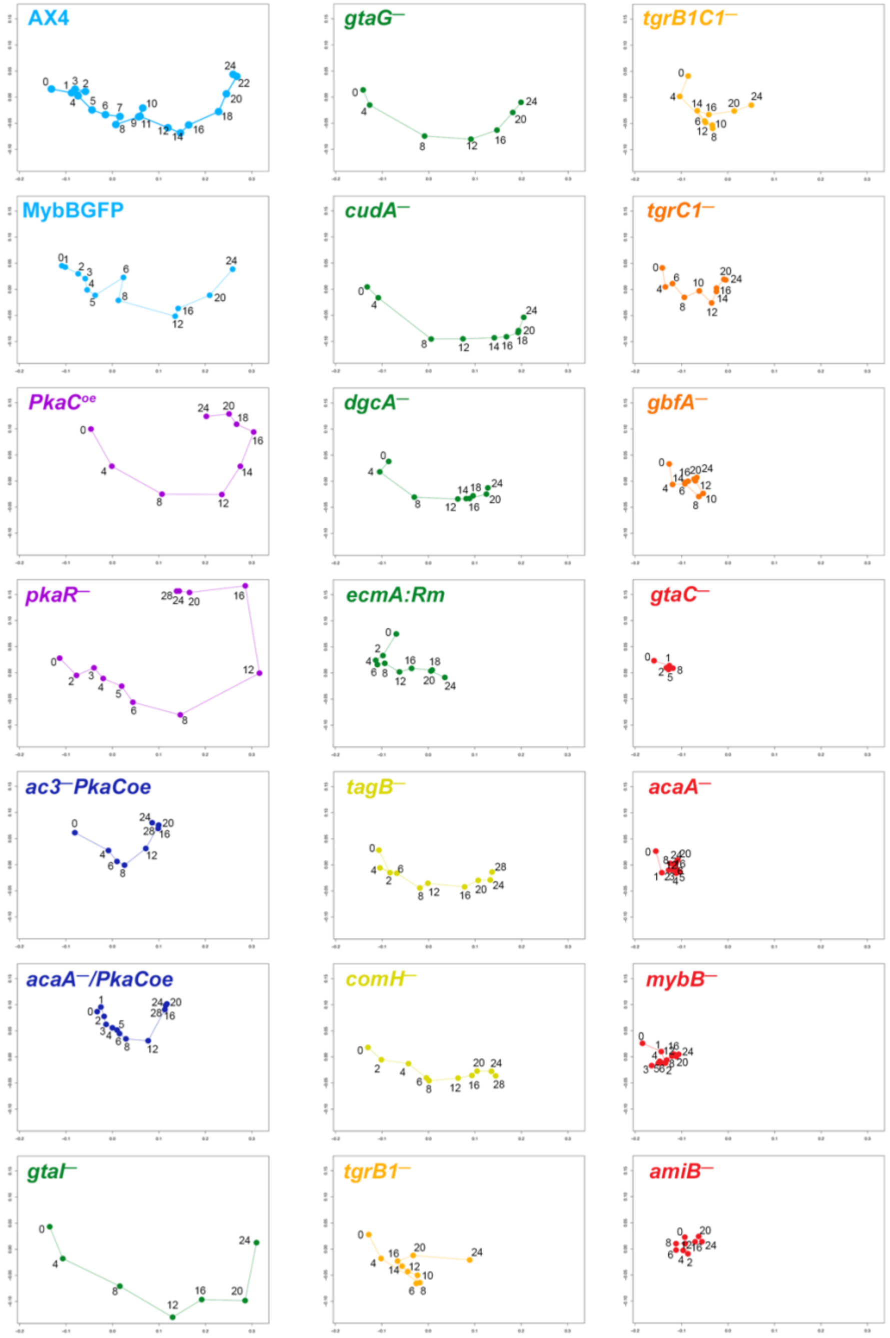

**B.**

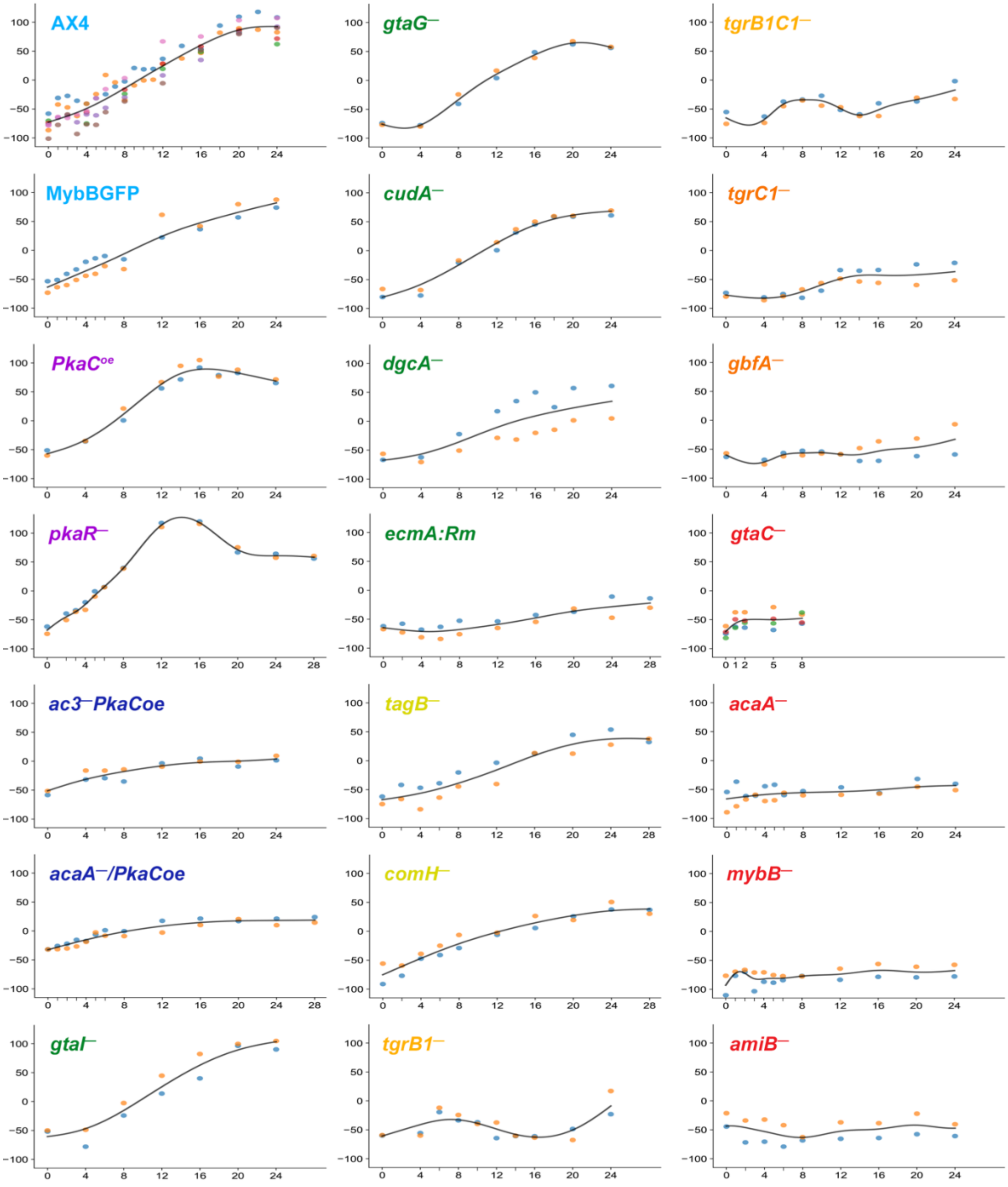

**Supplemental_Fig_S2 MDS and PCA plots of individual strains.** We analyzed the transcriptomes of the developing cells across time by RNA-seq and performed MDS (A) and PCA (B) using expression data from 2-7 replicates of each strain. The plots are identical to the respective analyses shown in aggregate in Figure 2, to provide better resolution. The PCA plots here include individual replicates of each strain plotted as different color circles to illustrate reproducibility.

**Supplemental_Fig_S3 Precocious spore and stalk differentiation in the *pkaR*⁻ strain**

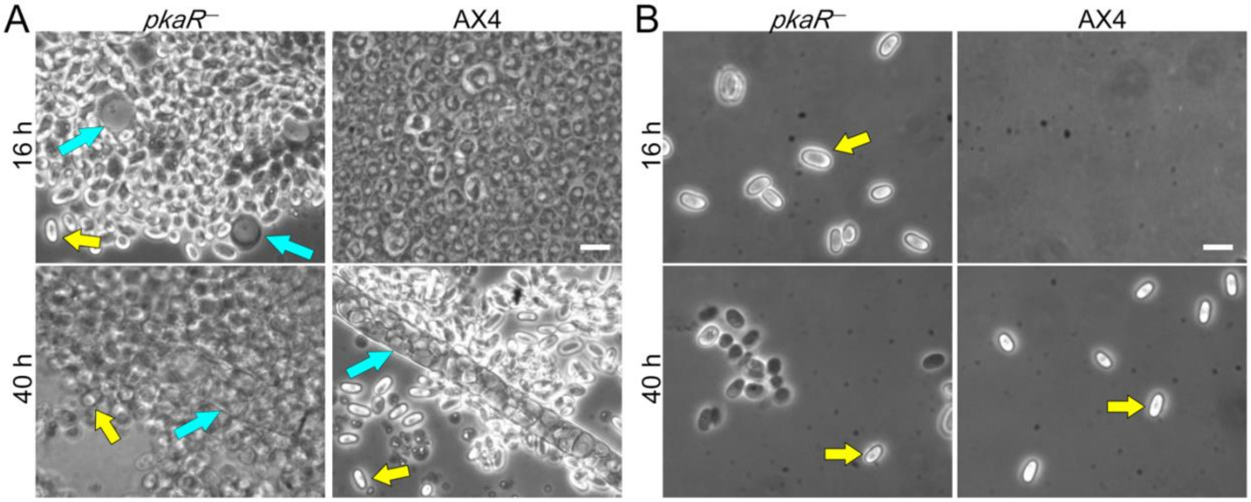

**Supplemental_Fig_S3 Precocious spore and stalk differentiation in the *pkaR*⁻ strain.** We developed AX4 and *pkaR*⁻ cells on nitrocellulose filters for 16 h and 40 h, as indicated. (A) We placed whole mounds on microscope slides, squashed them gently under a cover-slip and examined the cell morphology with phase-contrast microscopy. Blue arrows indicate stalk cells (16 h) or stalk tubes (40 h) and yellow arrows indicate spores. Bars = 20 μm. (B) We treated the developing cells with detergent to eliminate amoebae and imaged them with phase-contrast microscopy. Yellow arrows indicate spores. Bars = 20 μm.

**Supplemental_Fig_S4 Stage information on PCA plots**

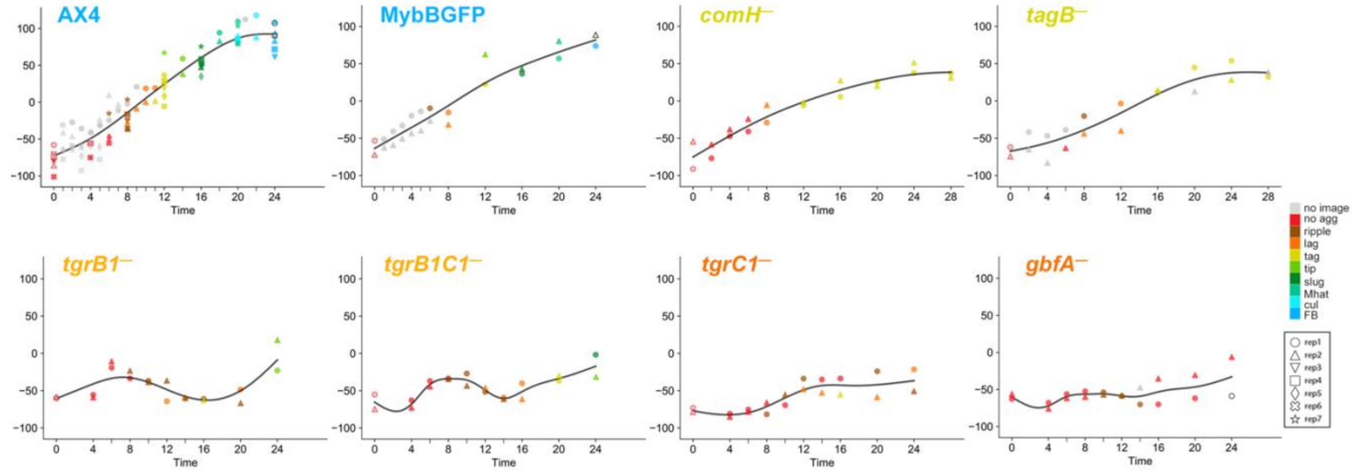

**Supplemental_Fig_S4  Stage information on PCA plots.** We added the representative stage information of each sample to the PC1 vs. time plots of the wild type, tight aggregate, and disaggregation phenotype groups (a subset of Supplemental_Fig_S2). PC1 (y-axis, arbitrary units) of each strain was plotted against time (x-axis, hours). The strain names are indicated in the plot and the color of the strain name represents the phenotype group: wild type (light blue), tight aggregate arrest (dark yellow), tight aggregate disaggregation (light orange), and loose aggregate disaggregation (dark orange). Representative morphological stages are plotted in different colors as indicated in the legend on the right. Samples at the zero time points were assigned a no agg stage. If the image was not captured for the samples at 0 h, red borders were added to gray symbols. White symbols with black outlines indicate samples for which representative stages were undetermined due to mixed morphologies.

**Supplemental Files provided in this section**
Supplemental_File_S1   Gene-set enrichment among milestone genes
Supplemental_File_S2   Gene-set enrichment among the regulons
Supplemental_File_S3   Gene-set enrichment in *tgr*-disaggregation and dedifferentiation
Supplemental_File_S4   Introductions to dictyExpress and Orange
Supplemental_File_S5   Computational methods
Supplemental_File_S6  Standard experimental methods
==============================================================

#### Supplemental_File_S1: Gene-set enrichment among milestone genes

All genes (except for non-expressed genes): 12,431 genes
Genes with Entrez ID (EID): 12,339 genes
Annotated genes: 4,510 genes (36.55%)

Milestone genes: 1,371 milestone genes at 8 stage transitions

**“Term”:** Enriched GO-term, KEGG-pathway, custom gene sets(Custom)
**“Ontology”:** GO:Biological process(BP), Cellular component(CC), Molecular function(MF), KE GG:pathway(Path.), Custom
**“Group”:** The number of genes with the term in the milestone gene set
**”Reference”:** The number of genes with the term in the reference (with EID)
**”FDR”:** false discovery rate, hypergeometric test
The bar graph size shows the fold enrichment and the color (see scale) represents the FDR.

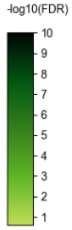

Selected Enrichment terms: FDR ≤ 0.25, gene number in the group ≥ 2

**1)  “no agg” to “ripple/stream” stage**
**Down-regulated**:
all genes: 294, with EID: 292, term-annotated genes: **132 genes**
**Enrichment annotation list:**

| Term | Ontology | Group | Reference | FDR |
| --- | --- | --- | --- | --- |
| Ascorbate and aldarate metabolism | KEGG: Path. | 2 (1.52%) | 5 (0.11%) | 7.65e-02 |
| Lysine biosynthesis | KEGG: Path. | 2 (1.52%) | 6 (0.13%) | 9.76e-02 |
| Folate biosynthesis | KEGG: Path. | 4 (3.03%) | 19 (0.42%) | 4.48e-02 |
| translation factor activity | GO: MF | 3 (2.27%) | 19 (0.42%) | 1.31e-01 |
| Pyruvate metabolism | KEGG: Path. | 4 (3.03%) | 29 (0.64%) | 8.36e-02 |
| Glycine | KEGG: Path. | 3 (2.27%) | 24 (0.53%) | 2.18e-01 |
| Glutathione metabolism | KEGG: Path. | 3 (2.27%) | 25 (0.55%) | 2.30e-01 |
| lysosome | GO: CC | 8 (6.06%) | 69 (1.53%) | 3.35e-02 |
| methyltransferase activity | GO: MF | 6 (4.55%) | 56 (1.24%) | 6.58e-02 |
| homeostatic process | GO: BP | 7 (5.30%) | 70 (1.55%) | 6.58e-02 |
| cofactor metabolic process | GO: BP | 7 (5.30%) | 72 (1.60%) | 6.58e-02 |
| oxidoreductase activity | GO: MF | 19 (14.39%) | 207 (4.59%) | 8.46e-04 |
| peptidase activity | GO: MF | 8 (6.06%) | 91 (2.02%) | 6.58e-02 |
| lipid metabolic process | GO: BP | 12 (9.09%) | 159 (3.53%) | 4.48e-02 |
| extracellular region | GO: CC | 11 (8.33%) | 157 (3.48%) | 6.58e-02 |
| Biosynthesis of secondary metabolites | KEGG: Path. | 18 (13.64%) | 281 (6.23%) | 3.78e-02 |
| vacuole | GO: CC | 8 (6.06%) | 127 (2.82%) | 2.18e-01 |
| response to stress | GO: BP | 22 (16.67%) | 358 (7.94%) | 3.35e-02 |
| small molecule metabolic process | GO: BP | 19 (14.39%) | 353 (7.83%) | 6.67e-02 |

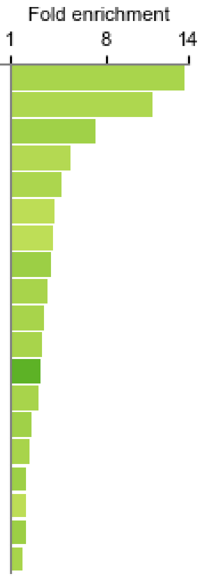

**Up-regulated:**
all genes: 247, with EID: 246, term-annotated genes: **72 genes**
**Enrichment annotation list**:

| Term | Ontology | Group | Reference | FDR |
| --- | --- | --- | --- | --- |
| pst gene | Custom | 17 (23.61%) | 495 (10.98%) | 1.09e-01 |

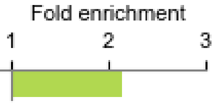

**2) “ripple/stream” to “lag” stage**
**Down-regulated:**
all genes: 1, with EID: 1, term-annotated genes: **0 genes**

**Up-regulated:**
all genes: 71, with EID: 70, term-annotated genes: **26 genes**
**Enrichment annotation list:** None

**3) “lag” to “tag” stage**
**Down-regulated:**
all genes: 11, with EID: 11, term-annotated genes: **2 genes**
**Enrichment annotation list**: None

**Up-regulated:**
all genes: 260, with EID: 259, term-annotated genes: **133 genes**
**Enrichment annotation list:**

| Term | Ontology | Group | Reference | FDR | Fold enrichment (1 – 12 – 22) |
|---|---|---|---|---|---|
| DNA replication | KEGG: Path. | 21 (15.79%) | 33 (0.73%) | 3.41e-23 | |
| chromosome segregation | GO: BP | 9 (6.77%) | 18 (0.40%) | 8.90e-09 | |
| Mismatch repair | KEGG: Path. | 9 (6.77%) | 19 (0.42%) | 1.37e-08 | |
| mitotic nuclear division | GO: BP | 6 (4.51%) | 15 (0.33%) | 1.91e-05 | |
| nuclear chromosome | GO: CC | 6 (4.51%) | 17 (0.38%) | 3.81e-05 | |
| chromosome | GO: CC | 22 (16.54%) | 70 (1.55%) | 4.34e-16 | |
| Base excision repair | KEGG: Path. | 8 (6.02%) | 28 (0.62%) | 7.80e-06 | |
| Non-homologous end-joining | KEGG: Path. | 2 (1.50%) | 7 (0.16%) | 6.65e-02 | |
| sig and sigN genes | Custom | 16 (12.03%) | 57 (1.26%) | 6.10e-11 | |
| Nucleotide excision repair | KEGG: Path. | 10 (7.52%) | 37 (0.82%) | 6.98e-07 | |
| chromosome organization | GO: BP | 14 (10.53%) | 74 (1.64%) | 2.43e-07 | |
| Pyrimidine metabolism | KEGG: Path. | 5 (3.76%) | 29 (0.64%) | 6.87e-03 | |
| chromatin/centromere | Custom | 3 (2.26%) | 18 (0.40%) | 6.58e-02 | |
| cell cycle | GO: BP | 26 (19.55%) | 167 (3.70%) | 2.69e-11 | |
| DNA metabolic process | GO: BP | 15 (11.28%) | 105 (2.33%) | 2.89e-06 | |
| cAMP pulse induced | Custom | 8 (6.02%) | 56 (1.24%) | 1.18e-03 | |
| microtubule organizing center | GO: CC | 7 (5.26%) | 50 (1.11%) | 3.00e-03 | |
| cell division | GO: BP | 14 (10.53%) | 115 (2.55%) | 3.81e-05 | |
| Homologous recombination | KEGG: Path. | 3 (2.26%) | 25 (0.55%) | 1.28e-01 | |
| mitotic cell cycle | GO: BP | 12 (9.02%) | 100 (2.22%) | 1.92e-04 | |
| helicase activity | GO: MF | 5 (3.76%) | 51 (1.13%) | 6.65e-02 | |
| Ubiquitin mediated proteolysis | KEGG: Path. | 5 (3.76%) | 61 (1.35%) | 1.22e-01 | |
| DNA binding | GO: MF | 17 (12.78%) | 221 (4.90%) | 1.25e-03 | |
| ATPase activity | GO: MF | 12 (9.02%) | 168 (3.73%) | 1.71e-02 | |
| cytoskeleton | GO: CC | 11 (8.27%) | 185 (4.10%) | 7.64e-02 | |
| psp gene | Custom | 17 (12.78%) | 391 (8.67%) | 2.25e-01 | |

#### 4) “tag” to “tip” stage

**Down-regulated**:

all genes: 20, with EID: 20, term-annotated genes: **9 genes**

**Enrichment annotation list:**

| Term | Ontology | Group | Reference | FDR | Fold enrichment (1 – 7 – 13) |
|---|---|---|---|---|---|
| cell adhesion | GO: BP | 2 (22.22%) | 83 (1.84%) | 1.22e-01 | |
| plasma membrane | GO: CC | 4 (44.44%) | 268 (5.94%) | 2.66e-02 | |
| cytoskeleton | GO: CC | 2 (22.22%) | 185 (4.10%) | 1.74e-01 | |
| cytoskeleton organization | GO: BP | 2 (22.22%) | 196 (4.35%) | 1.74e-01 | |
| vesicle-mediated transport | GO: BP | 2 (22.22%) | 233 (5.17%) | 2.07e-01 | |
| signal transduction | GO: BP | 3 (33.33%) | 350 (7.76%) | 1.73e-01 | |

**Up-regulated:**

all genes: 35, with EID: 35, term-annotated genes: **21 genes**

**Enrichment annotation list:**

| Term | Ontology | Group | Reference | FDR | Fold enrichment (1 – 48 – 94) |
|---|---|---|---|---|---|
| 57-aa protein family | Custom | 10 (47.62%) | 23 (0.51%) | 4.13e-18 | |
| gtaG-dependent short protein | Custom | 6 (28.57%) | 49 (1.09%) | 1.92e-07 | |
| cAMP pulse induced | Custom | 2 (9.52%) | 56 (1.24%) | 5.47e-02 | |
| hssA/2C/7E family | Custom | 3 (14.29%) | 95 (2.11%) | 2.29e-02 | |
| pst gene | Custom | 13 (61.90%) | 495 (10.98%) | 1.28e-07 | |
| extracellular region | GO: CC | 3 (14.29%) | 157 (3.48%) | 5.79e-02 | |

#### 5) “tip” to “slug” stage

**Down-regulated:**

No selected genes.

**Up-regulated:**
all genes: 12, with EID: 12, term-annotated genes: **12 genes**
**Enrichment annotation list:**

| Term | Ontology | Group | Reference | FDR | Fold enrichment (1 – 19 – 36) |
|---|---|---|---|---|---|
| hssA/2C/7E family | Custom | 9 (75.00%) | 95 (2.11%) | 8.14e-13 | |

##### 6) “slug” to “Mexican hat” stage
**Down-regulated:**
No selected genes.

**Up-regulated:**
all genes: 209, with EID: 209, term-annotated genes: **96 genes**
**Enrichment annotation list:**

| Term | Ontology | Group | Reference | FDR | Fold enrichment (1 – 3 – 5) |
|---|---|---|---|---|---|
| gtaG-dependent short protein | Custom | 5 (5.21%) | 49 (1.09%) | 1.06e-01 | |
| psp gene | Custom | 38 (39.58%) | 391 (8.67%) | 5.44e-15 | |
| extracellular region | GO: CC | 10 (10.42%) | 157 (3.48%) | 7.48e-02 | |

##### 7) “Mexican hat” to “culmination” stage
**Down-regulated:**
all genes: 9, with EID: 9, term-annotated genes: **7 genes**
**Enrichment annotation list:**

| Term | Ontology | Group | Reference | FDR | Fold enrichment (1 – 8 – 14) |
|---|---|---|---|---|---|
| hssA/2C/7E family | Custom | 2 (28.57%) | 95 (2.11%) | 1.29e-02 | |
| psp gene | Custom | 5 (71.43%) | 391 (8.67%) | 2.60e-04 | |

**Up-regulated**:
all genes: 45, with EID: 45, term-annotated genes: **13 genes**
**Enrichment annotation list:**

| Term | Ontology | Group | Reference | FDR | Fold enrichment (1 – 15 – 29) |
|---|---|---|---|---|---|
| gtaG-dependent short protein | Custom | 4 (30.77%) | 49 (1.09%) | 2.13e-04 | |
| sig and sigN genes | Custom | 3 (23.08%) | 57 (1.26%) | 6.51e-03 | |

##### 8) “culmination” to “fruiting body” stage
**Down-regulated:**
all genes: 3, with EID: 3, term-annotated genes: **2 genes**
**Enrichment annotation list**: None

**Up-regulated:**

all genes: 176, with EID: 175, term-annotated genes: **50 genes**

**Enrichment annotation list:**

| Term | Ontology | Group | Reference | FDR |
|---|---|---|---|---|
| Starch and sucrose metabolism | KEGG: Path. | 5 (10.00%) | 31 (0.69%) | 3.92e-04 |
| anatomical structure formation involved in morphogenesis | GO: BP | 8 (16.00%) | 74 (1.64%) | 3.48e-05 |
| Fructose and mannose metabolism | KEGG: Path. | 2 (4.00%) | 19 (0.42%) | 1.64e-01 |
| transferase activity | GO: MF | 2 (4.00%) | 24 (0.53%) | 2.24e-01 |
| gtaG-dependent short protein | Custom | 4 (8.00%) | 49 (1.09%) | 2.47e-02 |
| extracellular region | GO: CC | 12 (24.00%) | 157 (3.48%) | 5.07e-06 |
| hydrolase activity | GO: MF | 3 (6.00%) | 45 (1.00%) | 1.38e-01 |
| cell differentiation | GO: BP | 8 (16.00%) | 145 (3.22%) | 2.53e-03 |

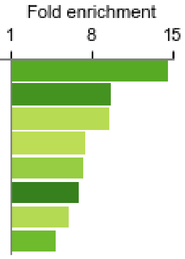

#### Supplemental_File_S2: Gene set enrichment among the regulons

All genes (except for non-expressed genes): 12,431 genes
Genes with Entrez ID (EID): 12,339 genes
Annotated genes: 4, 510 genes (36.55%)

Regulon clusters: 21 clusters containing 1099 selected genes

**"Term":** Enriched GO-term, KEGG-pathway, custom gene sets (Custom)
**"Ontology":** GO:Biological process(BP), Cellular component(CC), Molecular function(MF), KE GG:pathway(Path.), Custom
**"Group":** The number of genes with the term in the regulon cluster
**"Reference":** The number of genes with the term in the reference (with EID)
**"FDR":** false discovery rate, hypergeometric test
The bar graph size shows the fold enrichment and the color (see scale) represents the FDR.

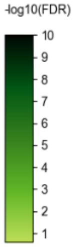

Selected Enrichment terms: FDR ≤ 0.25, gene number in the group ≥ 2

**Cluster 1**:
all genes: 30, with EID: 30, term-annotated genes: **22 genes**
**Enrichment annotation list:**

| Term | Ontology | Group | Reference | FDR |
|---|---|---|---|---|
| ribonucleoprotein complex assembly | GO: BP | 4 (18.18%) | 18 (0.40%) | 5.59e-06 |
| Ribosome biogenesis in eukaryotes | KEGG: Path. | 12 (54.55%) | 68 (1.51%) | 5.18e-16 |
| ribosome biogenesis | GO: BP | 10 (45.45%) | 61 (1.35%) | 3.25e-13 |
| rRNA binding | GO: MF | 2 (9.09%) | 13 (0.29%) | 5.15e-03 |
| nucleolus | GO: CC | 11 (50.00%) | 72 (1.60%) | 4.30e-14 |
| helicase activity | GO: MF | 2 (9.09%) | 51 (1.13%) | 5.01e-02 |
| methyltransferase activity | GO: MF | 2 (9.09%) | 56 (1.24%) | 5.37e-02 |
| RNA binding | GO: MF | 6 (27.27%) | 169 (3.75%) | 4.13e-04 |
| protein-containing complex assembly | GO: BP | 4 (18.18%) | 133 (2.95%) | 9.00e-03 |
| cellular component assembly | GO: BP | 4 (18.18%) | 201 (4.46%) | 3.35e-02 |

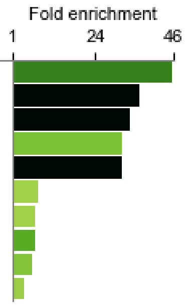

**Cluster 2**:

all genes: 66, with EID: 65, term-annotated genes: **40 genes**

**Enrichment annotation list**

| Term | Ontology | Group | Reference | FDR |
|---|---|---|---|---|
| Oxidative phosphorylation | KEGG: Path. | 9 (22.50%) | 54 (1.20%) | 3.49e-08 |
| Citrate cycle (TCA cycle) | KEGG: Path. | 4 (10.00%) | 30 (0.67%) | 2.94e-03 |
| Various types of N-glycan biosynthesis | KEGG: Path. | 2 (5.00%) | 22 (0.49%) | 1.16e-01 |
| cell morphogenesis | GO: BP | 2 (5.00%) | 23 (0.51%) | 1.16e-01 |
| protein folding | GO: BP | 3 (7.50%) | 37 (0.82%) | 5.03e-02 |
| Phagosome | KEGG: Path. | 4 (10.00%) | 52 (1.15%) | 1.77e-02 |
| unfolded protein binding | GO: MF | 2 (5.00%) | 27 (0.60%) | 1.43e-01 |
| N-Glycan biosynthesis | KEGG: Path. | 2 (5.00%) | 30 (0.67%) | 1.59e-01 |
| Propanoate metabolism | KEGG: Path. | 2 (5.00%) | 32 (0.71%) | 1.66e-01 |
| generation of precursor metabolites and energy | GO: BP | 4 (10.00%) | 77 (1.71%) | 5.03e-02 |
| Ribosome | KEGG: Path. | 4 (10.00%) | 97 (2.15%) | 8.51e-02 |
| mitochondrion | GO: CC | 9 (22.50%) | 229 (5.08%) | 2.94e-03 |
| Carbon metabolism | KEGG: Path. | 3 (7.50%) | 102 (2.26%) | 2.39e-01 |
| cytoplasmic vesicle | GO: CC | 6 (15.00%) | 211 (4.68%) | 8.51e-02 |
| transmembrane transporter activity | GO: MF | 4 (10.00%) | 149 (3.30%) | 1.86e-01 |
| transmembrane transport | GO: BP | 4 (10.00%) | 164 (3.64%) | 2.34e-01 |
| Biosynthesis of secondary metabolites | KEGG: Path. | 6 (15.00%) | 281 (6.23%) | 1.70e-01 |

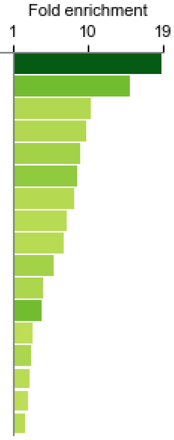

**Cluster 3**:

all genes: 68, with EID: 68, term-annotated genes: **66 genes**

**Enrichment annotation list:**

| Term | Ontology | Group | Reference | FDR |
|---|---|---|---|---|
| ribosome | GO: CC | 59 (89.39%) | 93 (2.06%) | 1.50e-100 |
| structural constituent of ribosome | GO: MF | 53 (80.30%) | 85 (1.88%) | 5.40e-87 |
| Ribosome | KEGG: Path. | 60 (90.91%) | 97 (2.15%) | 1.41e-101 |
| rRNA binding | GO: MF | 7 (10.61%) | 13 (0.29%) | 7.76e-10 |
| structural molecule activity | GO: MF | 53 (80.30%) | 117 (2.59%) | 1.08e-76 |
| translation | GO: BP | 52 (78.79%) | 169 (3.75%) | 4.18e-64 |
| RNA binding | GO: MF | 16 (24.24%) | 169 (3.75%) | 4.99e-09 |
| ribosome biogenesis | GO: BP | 3 (4.55%) | 61 (1.35%) | 1.84e-01 |
| cytosol | GO: CC | 8 (12.12%) | 203 (4.50%) | 3.12e-02 |

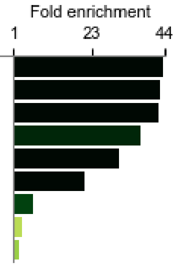

**Cluster 4**:

all genes: 55, with EID: 55, term-annotated genes: **23 genes**

**Enrichment annotation list:**

| Term | Ontology | Group | Reference | FDR |
|---|---|---|---|---|
| Endocytosis | KEGG: Path. | 5 (21.74%) | 75 (1.66%) | 4.29e-04 |
| cytoskeletal protein binding | GO: MF | 10 (43.48%) | 158 (3.50%) | 4.70e-08 |
| protein-containing complex assembly | GO: BP | 5 (21.74%) | 133 (2.95%) | 2.19e-03 |
| cytoskeleton organization | GO: BP | 6 (26.09%) | 196 (4.35%) | 2.19e-03 |
| cellular component assembly | GO: BP | 6 (26.09%) | 201 (4.46%) | 2.19e-03 |
| cytoskeleton | GO: CC | 5 (21.74%) | 185 (4.10%) | 7.34e-03 |
| plasma membrane | GO: CC | 7 (30.43%) | 268 (5.94%) | 2.19e-03 |
| cytosol | GO: CC | 4 (17.39%) | 203 (4.50%) | 5.80e-02 |
| response to stress | GO: BP | 7 (30.43%) | 358 (7.94%) | 6.23e-03 |
| cytoplasmic vesicle | GO: CC | 4 (17.39%) | 211 (4.68%) | 5.93e-02 |

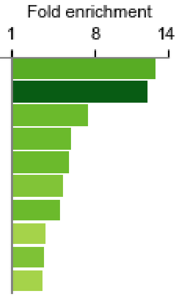

**Cluster 5**:
all genes: 41, with EID: 41, term-annotated genes: **40 genes**
**Enrichment annotation list:**

| Term | Ontology | Group | Reference | FDR |
|---|---|---|---|---|
| Proteasome | KEGG: Path. | 29 (72.50%) | 37 (0.82%) | 2.81e-57 |
| peptidase activity | GO: MF | 21 (52.50%) | 91 (2.02%) | 3.31e-25 |
| catabolic process | GO: BP | 25 (62.50%) | 250 (5.54%) | 2.40e-21 |

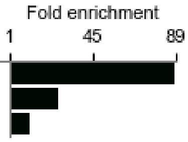

**Cluster 6**:
all genes: 74, with EID: 74, term-annotated genes: **41 genes**
**Enrichment annotation list:**

| Term | Ontology | Group | Reference | FDR |
|---|---|---|---|---|
| Glycosaminoglycan degradation | KEGG: Path. | 2 (4.88%) | 6 (0.13%) | 1.48e-02 |
| cAMP pulse induced | Custom | 9 (21.95%) | 56 (1.24%) | 4.65e-08 |
| cell adhesion | GO: BP | 5 (12.20%) | 83 (1.84%) | 1.39e-02 |
| extracellular region | GO: CC | 6 (14.63%) | 157 (3.48%) | 2.65e-02 |
| anatomical structure development | GO: BP | 15 (36.59%) | 438 (9.71%) | 7.49e-05 |
| cell differentiation | GO: BP | 4 (9.76%) | 145 (3.22%) | 1.88e-01 |
| locomotion | GO: BP | 5 (12.20%) | 189 (4.19%) | 1.50e-01 |
| signal transduction | GO: BP | 8 (19.51%) | 350 (7.76%) | 8.52e-02 |
| plasma membrane | GO: CC | 6 (14.63%) | 268 (5.94%) | 1.62e-01 |
| response to stress | GO: BP | 8 (19.51%) | 358 (7.94%) | 8.52e-02 |
| pst gene | Custom | 10 (24.39%) | 495 (10.98%) | 8.52e-02 |

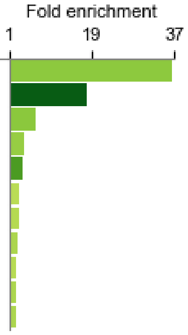

**Cluster 7**:
all genes: 40, with EID: 40, term-annotated genes: **14 genes**
**Enrichment annotation list:**

| Term | Ontology | Group | Reference | FDR |
|---|---|---|---|---|
| cell death | GO: BP | 2 (14.29%) | 25 (0.55%) | 1.93e-02 |
| RNA degradation | KEGG: Path. | 2 (14.29%) | 48 (1.06%) | 4.65e-02 |
| anatomical structure formation involved in morphogenesis | GO: BP | 3 (21.43%) | 74 (1.64%) | 1.93e-02 |
| cell differentiation | GO: BP | 3 (21.43%) | 145 (3.22%) | 4.65e-02 |
| regulatory transcription factor | Custom | 3 (21.43%) | 160 (3.55%) | 5.12e-02 |
| chemotaxis | Custom | 2 (14.29%) | 110 (2.44%) | 1.33e-01 |
| locomotion | GO: BP | 3 (21.43%) | 189 (4.19%) | 6.24e-02 |
| kinase activity | GO: MF | 5 (35.71%) | 327 (7.25%) | 1.93e-02 |
| anatomical structure development | GO: BP | 6 (42.86%) | 438 (9.71%) | 1.93e-02 |
| cytoskeleton | GO: CC | 2 (14.29%) | 185 (4.10%) | 2.36e-01 |
| transcriptional regulation and chromatin organization | Custom | 3 (21.43%) | 289 (6.41%) | 1.40e-01 |

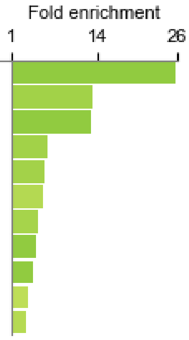

**Cluster 8**:
all genes: 23, with EID: 23, term-annotated genes: **10 genes**
**Enrichment annotation list:**

| Term | Ontology | Group | Reference | FDR |
|---|---|---|---|---|
| kinase activity | GO: MF | 3 (30.00%) | 327 (7.25%) | 1.71e-01 |
| psp gene | Custom | 3 (30.00%) | 391 (8.67%) | 1.71e-01 |

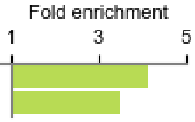

**Cluster 9**:
all genes: 20, with EID: 20, term-annotated genes: **18 genes**
**Enrichment annotation list:**

| Term | Ontology | Group | Reference | FDR | Fold enrichment (1, 32, 62) |
|---|---|---|---|---|---|
| sig and sigN genes | Custom | 14 (77.78%) | 57 (1.26%) | 5.61e-24 | |
| cAMP pulse induced | Custom | 7 (38.89%) | 56 (1.24%) | 1.77e-09 | |
| gtaG-dependent short protein | Custom | 3 (16.67%) | 49 (1.09%) | 8.77e-04 | |
| pst gene | Custom | 11 (61.11%) | 495 (10.98%) | 5.18e-07 | |

**Cluster 10**:
all genes: 36, with EID: 36, term-annotated genes: **19 genes**
**Enrichment annotation list:**

| Term | Ontology | Group | Reference | FDR | Fold enrichment (1, 41, 80) |
|---|---|---|---|---|---|
| chromosome segregation | GO: BP | 6 (31.58%) | 18 (0.40%) | 6.51e-10 | |
| mitotic nuclear division | GO: BP | 5 (26.32%) | 15 (0.33%) | 2.27e-08 | |
| DNA replication | KEGG: Path. | 5 (26.32%) | 33 (0.73%) | 1.03e-06 | |
| chromosome | GO: CC | 6 (31.58%) | 70 (1.55%) | 1.15e-06 | |
| chromosome organization | GO: BP | 6 (31.58%) | 74 (1.64%) | 1.41e-06 | |
| cell cycle | GO: BP | 13 (68.42%) | 167 (3.70%) | 1.07e-13 | |
| mitotic cell cycle | GO: BP | 7 (36.84%) | 100 (2.22%) | 6.70e-07 | |
| cell division | GO: BP | 7 (36.84%) | 115 (2.55%) | 1.15e-06 | |
| microtubule organizing center | GO: CC | 3 (15.79%) | 50 (1.11%) | 3.09e-03 | |
| cytoskeleton | GO: CC | 7 (36.84%) | 185 (4.10%) | 1.99e-05 | |
| cytoskeleton organization | GO: BP | 5 (26.32%) | 196 (4.35%) | 3.09e-03 | |
| cytoskeletal protein binding | GO: MF | 4 (21.05%) | 158 (3.50%) | 9.62e-03 | |
| DNA metabolic process | GO: BP | 2 (10.53%) | 105 (2.33%) | 1.57e-01 | |

**Cluster 11**:
all genes: 77, with EID: 77, term-annotated genes: **24 genes**
**Enrichment annotation list:**

| Term | Ontology | Group | Reference | FDR | Fold enrichment (1, 8, 14) |
|---|---|---|---|---|---|
| cAMP pulse induced | Custom | 4 (16.67%) | 56 (1.24%) | 4.62e-03 | |
| Starch and sucrose metabolism | KEGG: Path. | 2 (8.33%) | 31 (0.69%) | 1.13e-01 | |
| cell differentiation | GO: BP | 7 (29.17%) | 145 (3.22%) | 3.29e-04 | |
| cell adhesion | GO: BP | 3 (12.50%) | 83 (1.84%) | 1.13e-01 | |
| anatomical structure development | GO: BP | 7 (29.17%) | 438 (9.71%) | 1.01e-01 | |

**Cluster 12**:
all genes: 57, with EID: 57, term-annotated genes: **35 genes**
**Enrichment annotation list:**

| Term | Ontology | Group | Reference | FDR | Fold enrichment (1, 9, 17) |
|---|---|---|---|---|---|
| cell-cell signaling | GO: BP | 2 (5.71%) | 16 (0.35%) | 7.65e-02 | |
| psp gene | Custom | 18 (51.43%) | 391 (8.67%) | 2.10e-09 | |
| extracellular region | GO: CC | 7 (20.00%) | 157 (3.48%) | 2.80e-03 | |

**Cluster 13**:
all genes: 64, with EID: 64, term-annotated genes: **51 genes**
**Enrichment annotation list:**

| Term | Ontology | Group | Reference | FDR |
|---|---|---|---|---|
| 57-aa protein family | Custom | 17 (33.33%) | 23 (0.51%) | 7.13e-29 |
| gtaG-dependent short protein | Custom | 17 (33.33%) | 49 (1.09%) | 1.27e-21 |
| hssA/2C/7E family | Custom | 23 (45.10%) | 95 (2.11%) | 1.84e-25 |
| cAMP pulse induced | Custom | 4 (7.84%) | 56 (1.24%) | 1.24e-02 |
| pst gene | Custom | 28 (54.90%) | 495 (10.98%) | 5.22e-14 |

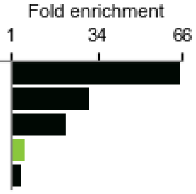

**Cluster 14:**
all genes: 19, with EID: 19, term-annotated genes: **16 genes**
**Enrichment annotation list:**

| Term | Ontology | Group | Reference | FDR |
|---|---|---|---|---|
| hssA/2C/7E family | Custom | 14 (87.50%) | 95 (2.11%) | 7.32e-22 |
| gtaG-dependent short protein | Custom | 2 (12.50%) | 49 (1.09%) | 3.15e-02 |

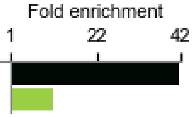

**Cluster 15:**
all genes: 91, with EID: 91, term-annotated genes: **67 genes**
**Enrichment annotation list:**

| Term | Ontology | Group | Reference | FDR |
|---|---|---|---|---|
| cell wall | GO: CC | 7 (10.45%) | 13 (0.29%) | 4.08e-09 |
| external encapsulating structure | GO: CC | 7 (10.45%) | 13 (0.29%) | 4.08e-09 |
| cell wall organization or biogenesis | GO: BP | 5 (7.46%) | 15 (0.33%) | 2.20e-05 |
| Starch and sucrose metabolism | KEGG: Path. | 4 (5.97%) | 31 (0.69%) | 8.54e-03 |
| anatomical structure formation involved in morphogenesis | GO: BP | 9 (13.43%) | 74 (1.64%) | 1.73e-05 |
| psp gene | Custom | 47 (70.15%) | 391 (8.67%) | 6.90e-34 |
| cAMP pulse induced | Custom | 6 (8.96%) | 56 (1.24%) | 1.46e-03 |
| cell differentiation | GO: BP | 10 (14.93%) | 145 (3.22%) | 4.91e-04 |
| carbohydrate metabolic process | GO: BP | 4 (5.97%) | 88 (1.95%) | 2.45e-01 |
| extracellular region | GO: CC | 6 (8.96%) | 157 (3.48%) | 1.85e-01 |
| anatomical structure development | GO: BP | 13 (19.40%) | 438 (9.71%) | 8.03e-02 |

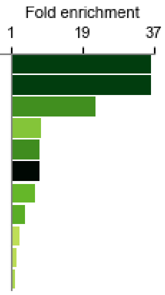

**Cluster 16**:
all genes: 74, with EID: 74, term-annotated: **39 genes**
**Enrichment annotation list:**

| Term | Ontology | Group | Reference | FDR |
|---|---|---|---|---|
| gtaG-dependent short protein | Custom | 4 (10.26%) | 49 (1.09%) | 7.65e-03 |
| hssA/2C/7E family | Custom | 5 (12.82%) | 95 (2.11%) | 9.14e-03 |
| extracellular region | GO: CC | 8 (20.51%) | 157 (3.48%) | 6.64e-04 |
| psp gene | Custom | 17 (43.59%) | 391 (8.67%) | 1.62e-07 |
| pst gene | Custom | 9 (23.08%) | 495 (10.98%) | 1.35e-01 |

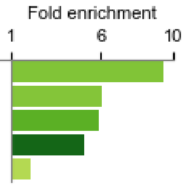

**Cluster 17**:
all genes: 61, with EID: 59, term-annotated genes: **22 genes**
**Enrichment annotation list:**

| Term | Ontology | Group | Reference | FDR |
|---|---|---|---|---|
| sig and sigN genes | Custom | 5 (22.73%) | 57 (1.26%) | 9.05e-05 |
| gtaG-dependent short protein | Custom | 4 (18.18%) | 49 (1.09%) | 7.79e-04 |
| psp gene | Custom | 11 (50.00%) | 391 (8.67%) | 1.60e-05 |

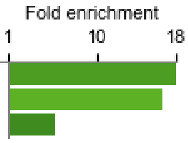

**Cluster 18**:
all genes: 46, with EID: 45, term-annotated genes: **19 genes**
**Enrichment annotation list:**

| Term | Ontology | Group | Reference | FDR |
|---|---|---|---|---|
| anatomical structure formation involved in morphogenesis | GO: BP | 3 (15.79%) | 74 (1.64%) | 2.72e-02 |
| extracellular region | GO: CC | 6 (31.58%) | 157 (3.48%) | 7.23e-04 |
| cell differentiation | GO: BP | 3 (15.79%) | 145 (3.22%) | 1.30e-01 |
| psp gene | Custom | 7 (36.84%) | 391 (8.67%) | 8.41e-03 |
| oxidoreductase activity | GO: MF | 3 (15.79%) | 207 (4.59%) | 1.84e-01 |

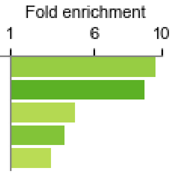

**Cluster 19**:
all genes: 58, with EID: 58, term-annotated genes: **18 genes**
**Enrichment annotation list:**

| Term | Ontology | Group | Reference | FDR |
|---|---|---|---|---|
| hydrolase activity | GO: MF | 2 (11.11%) | 45 (1.00%) | 6.52e-02 |
| gtaG-dependent short protein | Custom | 2 (11.11%) | 49 (1.09%) | 6.52e-02 |
| cell adhesion | GO: BP | 3 (16.67%) | 83 (1.84%) | 4.22e-02 |
| anatomical structure formation involved in morphogenesis | GO: BP | 2 (11.11%) | 74 (1.64%) | 9.00e-02 |
| extracellular region | GO: CC | 4 (22.22%) | 157 (3.48%) | 4.22e-02 |
| hssA/2C/7E family | Custom | 2 (11.11%) | 95 (2.11%) | 1.26e-01 |
| cell differentiation | GO: BP | 3 (16.67%) | 145 (3.22%) | 6.52e-02 |
| regulatory transcription factor | Custom | 3 (16.67%) | 160 (3.55%) | 7.25e-02 |
| pst gene | Custom | 6 (33.33%) | 495 (10.98%) | 6.52e-02 |
| transcriptional regulation and chromatin organization | Custom | 3 (16.67%) | 289 (6.41%) | 1.99e-01 |

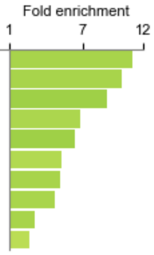

**Cluster 20:**
all genes: 54, with EID: 54, term-annotated genes: **20 genes**
**Enrichment annotation list:**

| Term | Ontology | Group | Reference | FDR |
|---|---|---|---|---|
| gtaG-dependent short protein | Custom | 3 (15.00%) | 49 (1.09%) | 3.50e-02 |
| Phagosome | KEGG: Path. | 2 (10.00%) | 52 (1.15%) | 2.10e-01 |
| hssA/2C/7E family | Custom | 3 (15.00%) | 95 (2.11%) | 1.15e-01 |

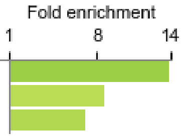

**Cluster 21**:
all genes: 45, with EID: 45, term-annotated genes: **12 genes**
**Enrichment annotation list:**

| Term | Ontology | Group | Reference | FDR |
|---|---|---|---|---|
| sig and sigN genes | Custom | 2 (16.67%) | 57 (1.26%) | 1.24e-01 |
| psp gene | Custom | 8 (66.67%) | 391 (8.67%) | 2.81e-05 |

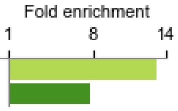

### Supplemental_File_S3: Gene-set enrichment in *tgr*-disaggregation and dedifferentiation

All genes (except for non-expressed genes): 12,431 genes
Genes with Entrez ID (EID): 12,339 genes
Annotated genes: 4,510 genes (36.55%)

**“Term”:** Enriched GO-term, KEGG-pathway, custom gene sets (Custom)
**“Ontology”:** GO:Biological process(BP), Cellular component(CC), Molecular function(MF), KEGG:pathway(Path.), Custom
**“Group”:** The number of genes with the term in the disagg gene set
**”Reference”:** The number of genes with the term in the reference (with EID)
**”FDR”:** false discovery rate, hypergeometric test
The bar graph size shows the fold enrichment and the color (see scale) represents the FDR.

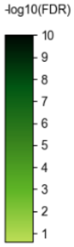

Selected Enrichment terms: FDR ≤ 0.25, gene number in the group ≥ 2

##### *tgr*-disagg 8 hr vs 6hr (padj ≦ 0.01, FoldChange ≧ 2.5)

all genes: 72, with EID: 70, term-annotated genes: **31 genes** (44.0%)

**Enrichment annotation list:**

| Term | Ontology | Group | Reference | FDR |
|---|---|---|---|---|
| Lysine degradation | KEGG: Path. | 2 (6.45%) | 22 (0.49%) | 1.47e-01 |
| Tryptophan metabolism | KEGG: Path. | 2 (6.45%) | 24 (0.53%) | 1.47e-01 |
| lipid binding | GO: MF | 4 (12.90%) | 59 (1.31%) | 4.79e-02 |
| membrane organization | GO: BP | 3 (9.68%) | 53 (1.18%) | 1.47e-01 |
| homeostatic process | GO: BP | 3 (9.68%) | 70 (1.55%) | 1.47e-01 |
| anatomical structure development | GO: BP | 8 (25.81%) | 438 (9.71%) | 1.47e-01 |
| pst gene | Custom | 8 (25.81%) | 495 (10.98%) | 1.71e-01 |

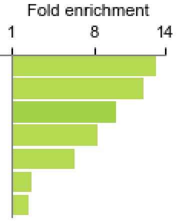

##### *tgr*-disagg 12hr vs 8hr (padj ≦ 0.01, FoldChange ≧ 2.5)

all genes: 218, with EID: 217, term-annotated genes: **102 genes** (47.0%)

**Enrichment annotation list:**

| Term | Ontology | Group | Reference | FDR |
|---|---|---|---|---|
| ribosome biogenesis | GO: BP | 34 (33.33%) | 61 (1.35%) | 2.18e-40 |
| nucleolus | GO: CC | 34 (33.33%) | 72 (1.60%) | 2.32e-37 |
| Ribosome biogenesis in eukaryotes | KEGG: Path. | 30 (29.41%) | 68 (1.51%) | 1.10e-31 |
| ribonucleoprotein complex assembly | GO: BP | 5 (4.90%) | 18 (0.40%) | 4.97e-04 |
| rRNA binding | GO: MF | 3 (2.94%) | 13 (0.29%) | 2.48e-02 |
| helicase activity | GO: MF | 8 (7.84%) | 51 (1.13%) | 2.43e-04 |
| nucleocytoplasmic transport | GO: BP | 4 (3.92%) | 26 (0.58%) | 2.48e-02 |
| RNA binding | GO: MF | 21 (20.59%) | 169 (3.75%) | 1.57e-09 |
| methyltransferase activity | GO: MF | 6 (5.88%) | 56 (1.24%) | 1.76e-02 |
| nucleotidyltransferase activity | GO: MF | 3 (2.94%) | 29 (0.64%) | 1.83e-01 |
| nucleoplasm | GO: CC | 7 (6.86%) | 91 (2.02%) | 3.23e-02 |
| ATPase activity | GO: MF | 10 (9.80%) | 168 (3.73%) | 3.23e-02 |

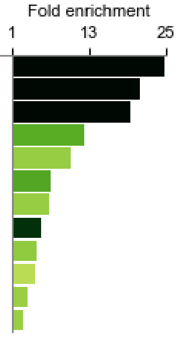

**dediff_medium 0.5-2 hr vs buf allT (padj ≤ 0.01, FoldChange ≥ 4)**
all genes: 360, with EID: 359, term-annotated genes: **162 genes** (45.0%)
**Enrichment annotation list:**

| Term | Ontology | Group | Reference | FDR |
|---|---|---|---|---|
| Ascorbate and aldarate metabolism | KEGG: Path. | 2 (1.23%) | 5 (0.11%) | 1.03e-01 |
| Pentose and glucuronate interconversions | KEGG: Path. | 2 (1.23%) | 6 (0.13%) | 1.42e-01 |
| Ribosome biogenesis in eukaryotes | KEGG: Path. | 17 (10.49%) | 68 (1.51%) | 1.61e-08 |
| Purine metabolism | KEGG: Path. | 10 (6.17%) | 50 (1.11%) | 2.35e-04 |
| nucleolus | GO: CC | 13 (8.02%) | 72 (1.60%) | 8.41e-05 |
| ribosome biogenesis | GO: BP | 11 (6.79%) | 61 (1.35%) | 2.35e-04 |
| Alanine | KEGG: Path. | 4 (2.47%) | 23 (0.51%) | 9.26e-02 |
| methyltransferase activity | GO: MF | 9 (5.56%) | 56 (1.24%) | 3.44e-03 |
| RNA polymerase | KEGG: Path. | 4 (2.47%) | 25 (0.55%) | 1.03e-01 |
| Folate biosynthesis | KEGG: Path. | 3 (1.85%) | 19 (0.42%) | 1.92e-01 |
| 2-Oxocarboxylic acid metabolism | KEGG: Path. | 3 (1.85%) | 19 (0.42%) | 1.92e-01 |
| Pyrimidine metabolism | KEGG: Path. | 4 (2.47%) | 29 (0.64%) | 1.45e-01 |
| Biosynthesis of amino acids | KEGG: Path. | 8 (4.94%) | 62 (1.37%) | 2.20e-02 |
| isomerase activity | GO: MF | 6 (3.70%) | 50 (1.11%) | 9.26e-02 |
| homeostatic process | GO: BP | 8 (4.94%) | 70 (1.55%) | 4.38e-02 |
| lyase activity | GO: MF | 7 (4.32%) | 66 (1.46%) | 9.26e-02 |
| Carbon metabolism | KEGG: Path. | 9 (5.56%) | 102 (2.26%) | 1.02e-01 |
| small molecule metabolic process | GO: BP | 31 (19.14%) | 353 (7.83%) | 8.91e-05 |
| ligase activity | GO: MF | 7 (4.32%) | 80 (1.77%) | 1.74e-01 |
| oxidoreductase activity | GO: MF | 18 (11.11%) | 207 (4.59%) | 7.92e-03 |
| mitochondrion | GO: CC | 18 (11.11%) | 229 (5.08%) | 2.05e-02 |
| Biosynthesis of secondary metabolites | KEGG: Path. | 21 (12.96%) | 281 (6.23%) | 1.74e-02 |
| cellular amino acid metabolic process | GO: BP | 9 (5.56%) | 124 (2.75%) | 2.11e-01 |

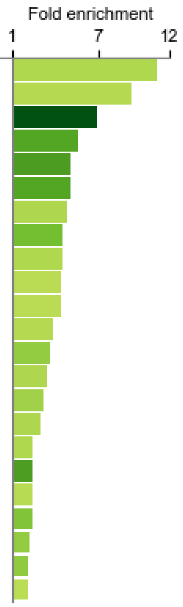

| | disaggregation_8to12 | | dedifferentiation | |
|---|---|---|---|---|
| | **gene #** | **/list** | **gene #** | **/list** |
| Milestone_noAgg_down (294 genes) | 10 | 3% | 48 | 16% |
| Regulon 1 (30 genes) | 30 | 100% | 15 | 50% |
| U3 related_rRNA processing (56 genes) | 45 | 80% | 21 | 38% |
| G-protein-coupled receptor (28 genes) | 4 | 14.3% | 1 | 3.6% |
| Overlap between two sets | 70 | 19% | 70 | 32% |

DE on data from Nichols, et al. (2020): dediff_set
dediff_set overlap with all genes: 2.9%

overlap between disaggregation_8to12 group and dediff_set: 32.1%
p-val: 9.25E-55

overlap between disaggregation_6to8 group and dediff_set: 2.8%
p-val: 6.21E-01

#### Supplemental_File_S4 Introduction to data mining in dictyExpress and Orange

#### A. An introduction to dictyExpress

Open dictyExpress in a web browser: https://dictyexpress.research.bcm.edu, and press "Run dictyExpress". If you are new to dictyExpress, follow the brief online tutorial before you proceed with the following suggestion.

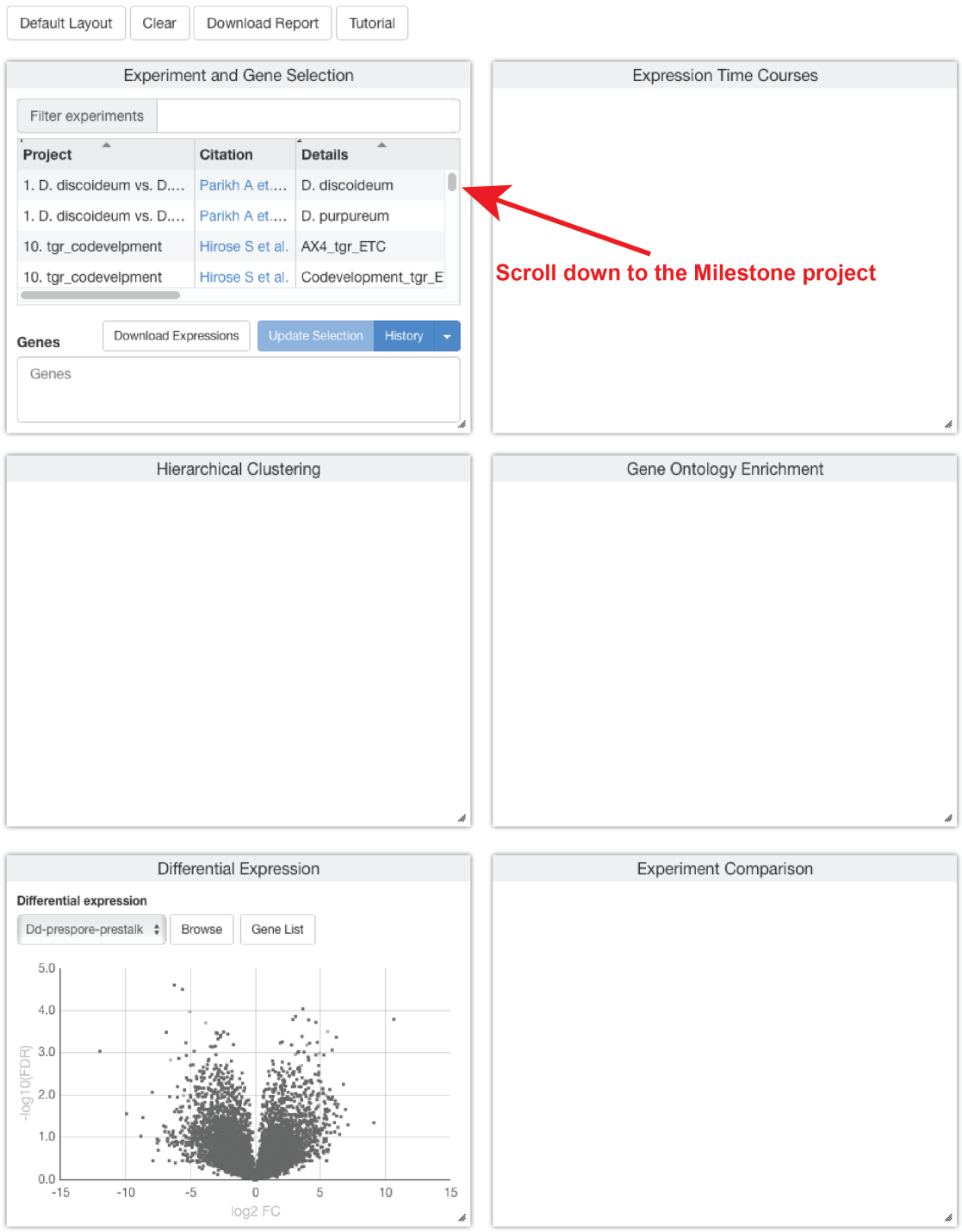

Select one of the Milestone project experiments in the 'Experiment and Gene Selection' panel. We recommend starting with AX4.

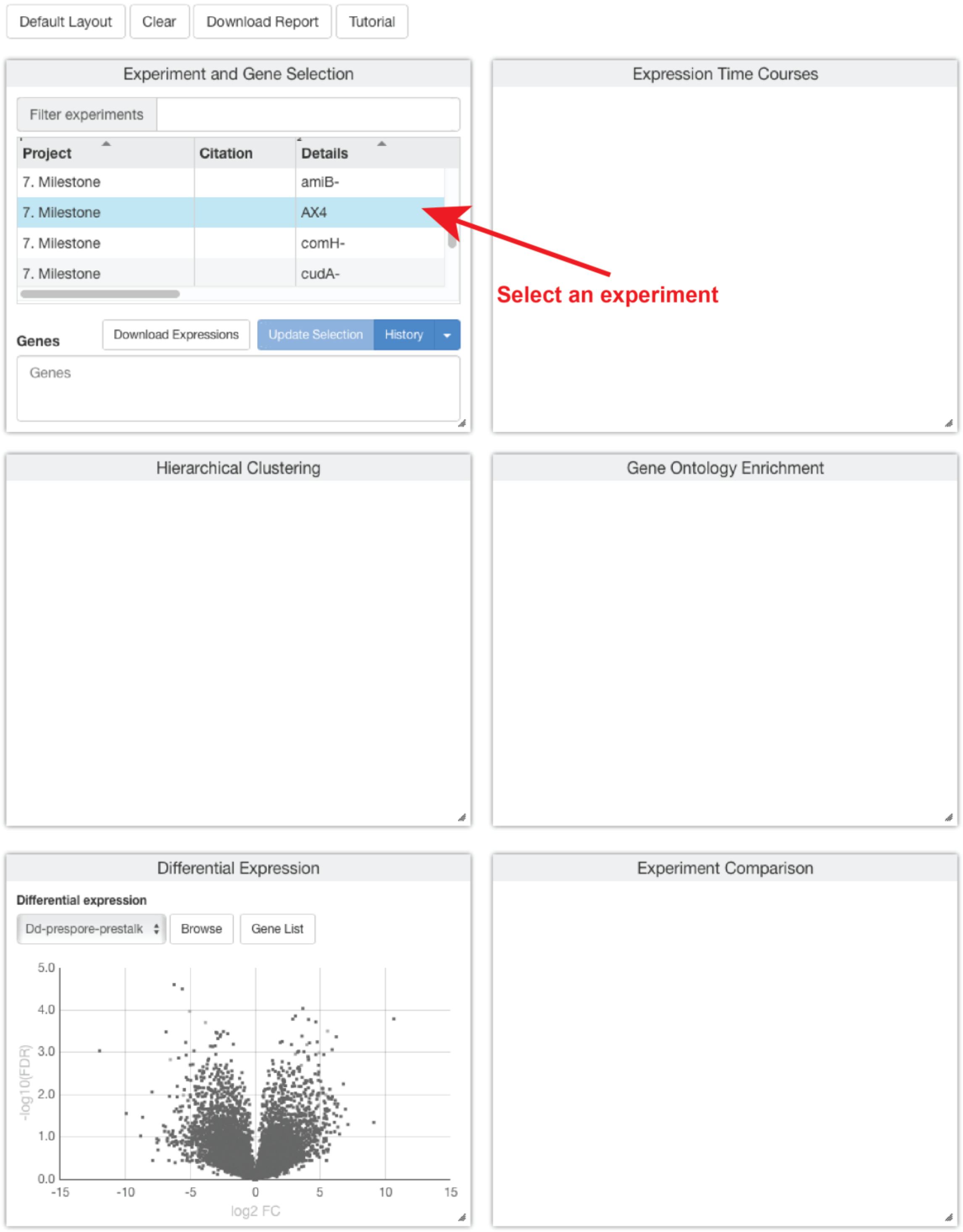

Type a gene name in the ‘Genes’ box. In this example we chose the actin gene Act6 by typing ‘act’ and selecting a gene from the ensuing drop-down menu.

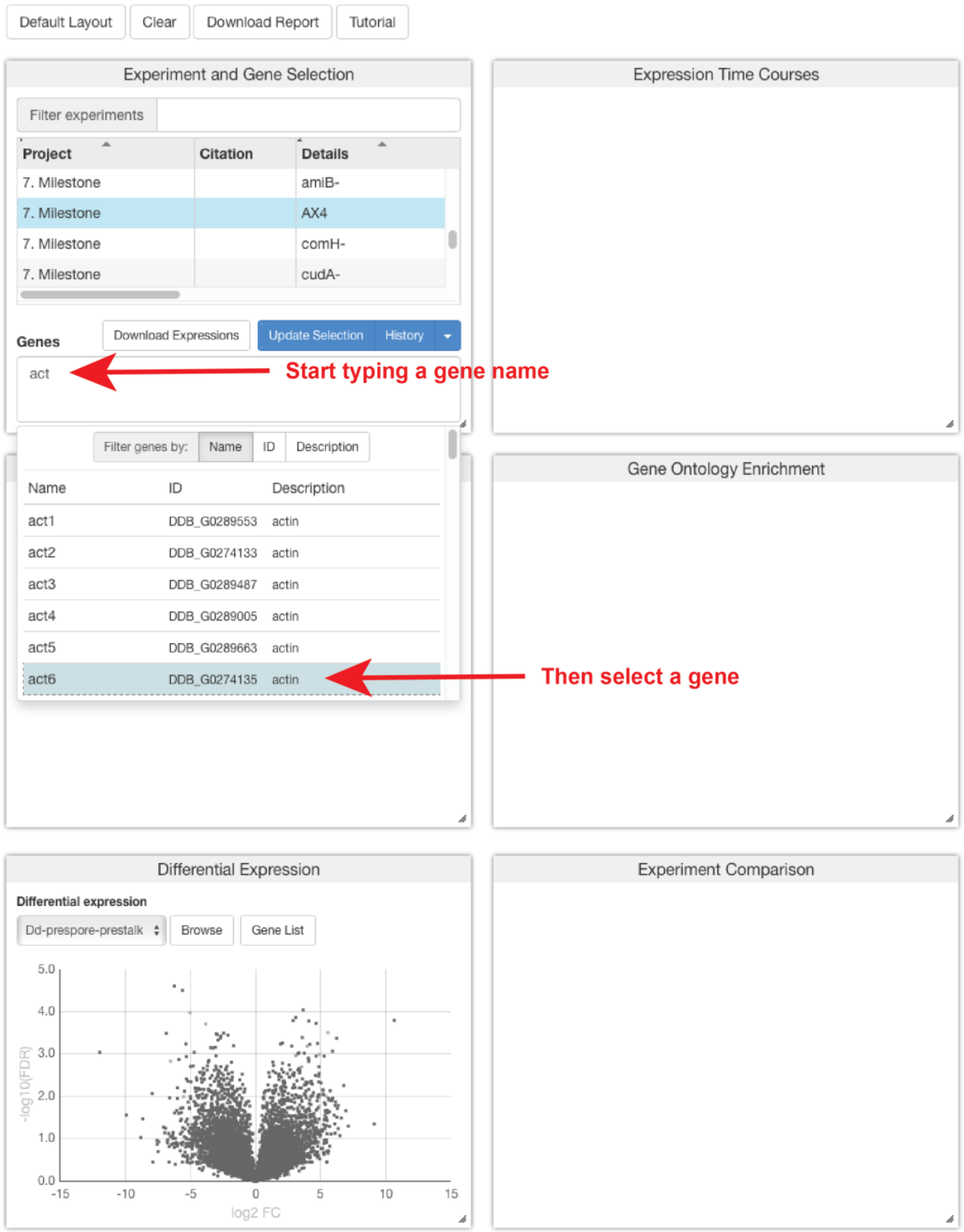

Pressing the ‘Update Selection’ button or hitting ‘return’ on your keyboard propagates the gene selection in all the other panels.

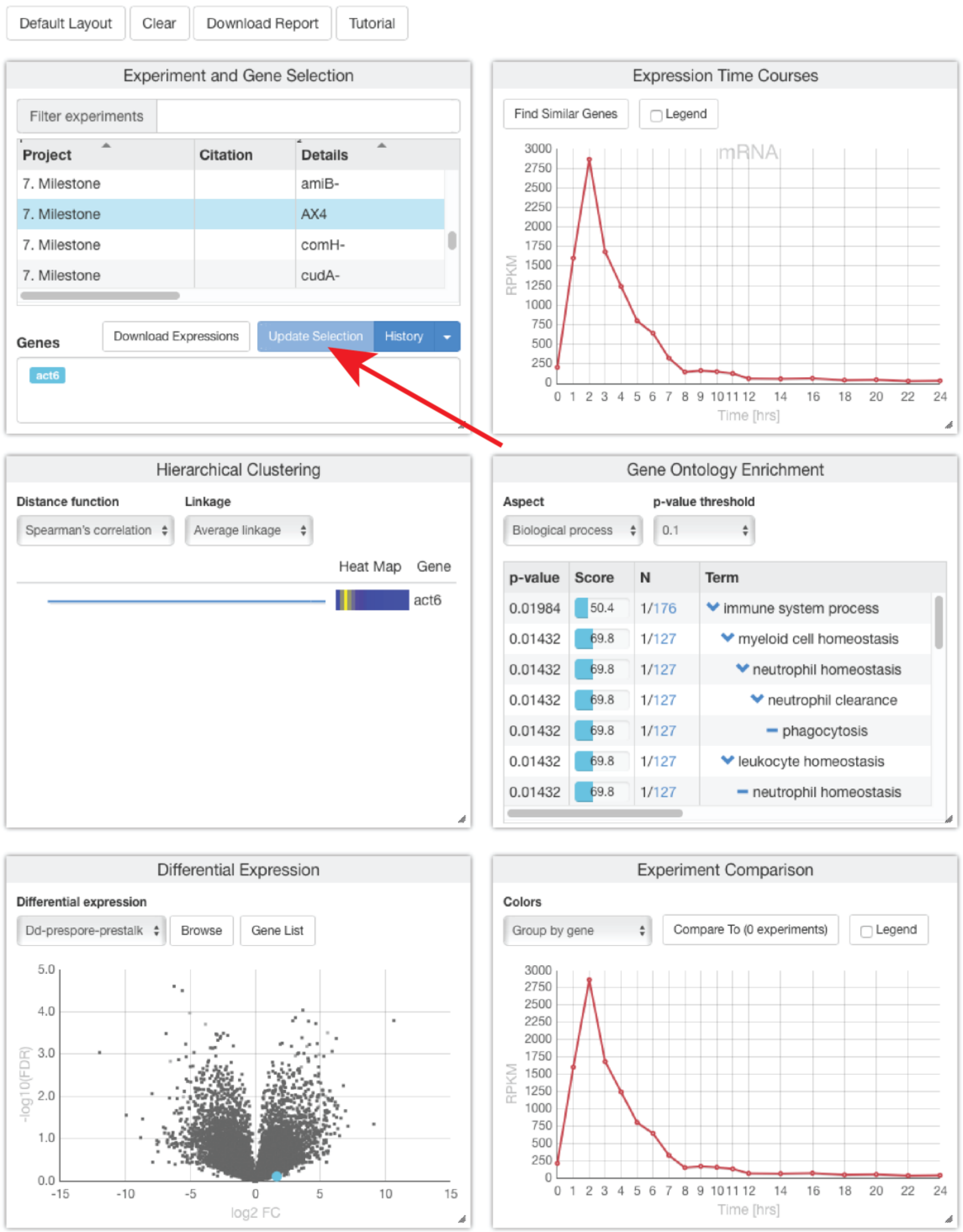

To compare the temporal and developmental expression patterns of actin 6 between several strains, click the ‘Compare To’ button in the ‘Experiment Comparison’ panel.

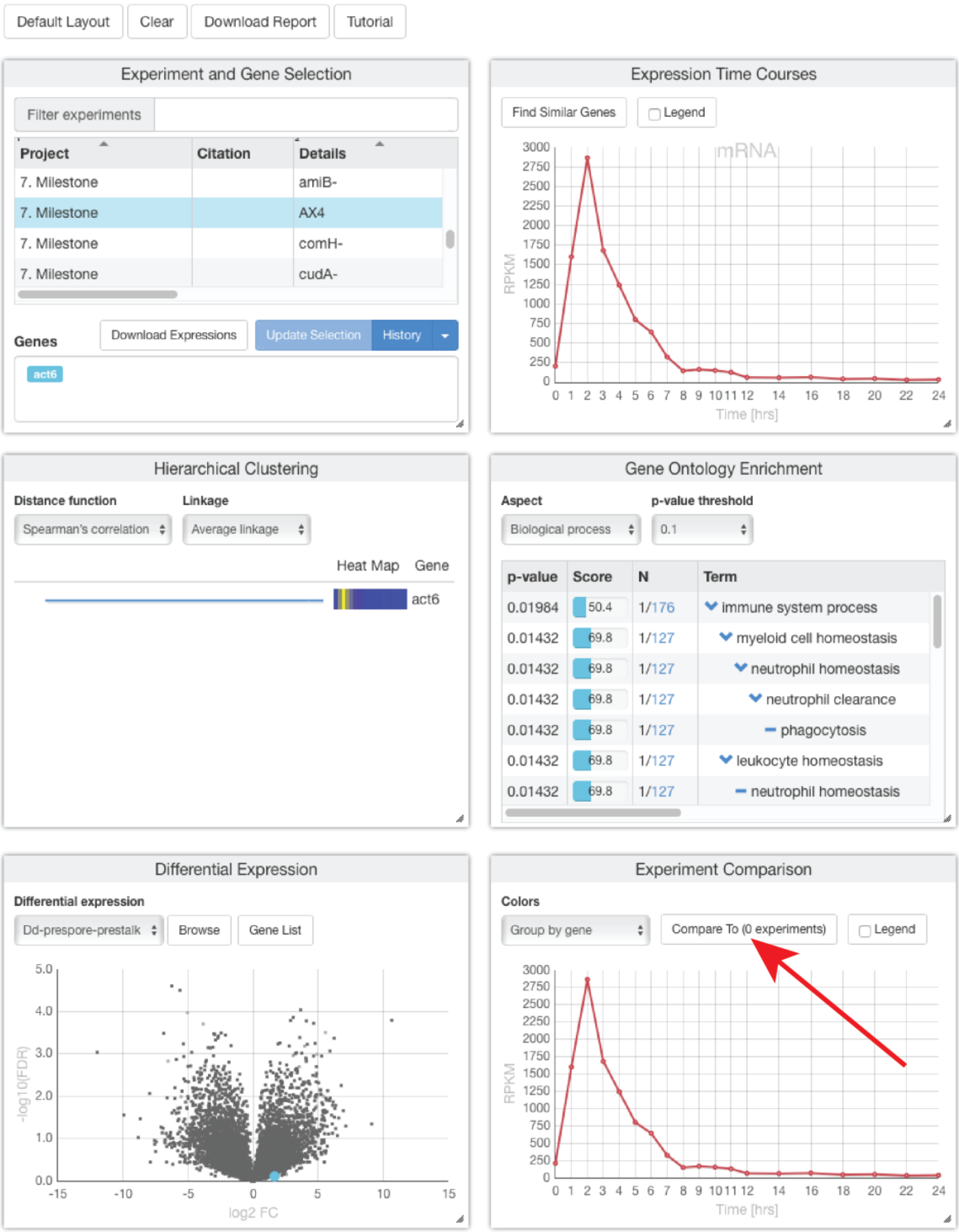

Select the desired experiments from the pop-up menu. Here, we selected the aggregationless mutant *acaA*⁻ and the precocious developer *pkaC*$^{OE}$.

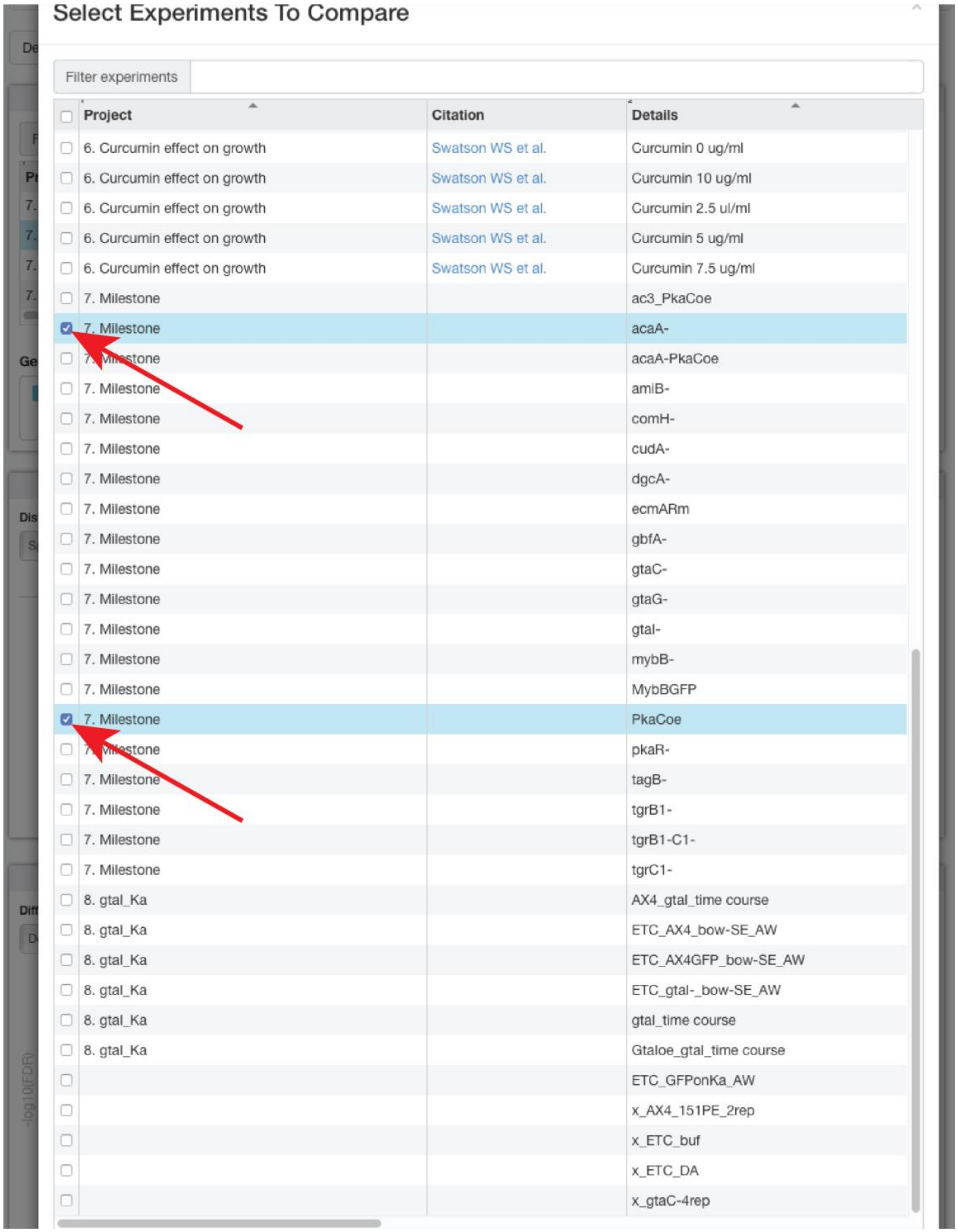

Close the selection menu to observe the comparison.

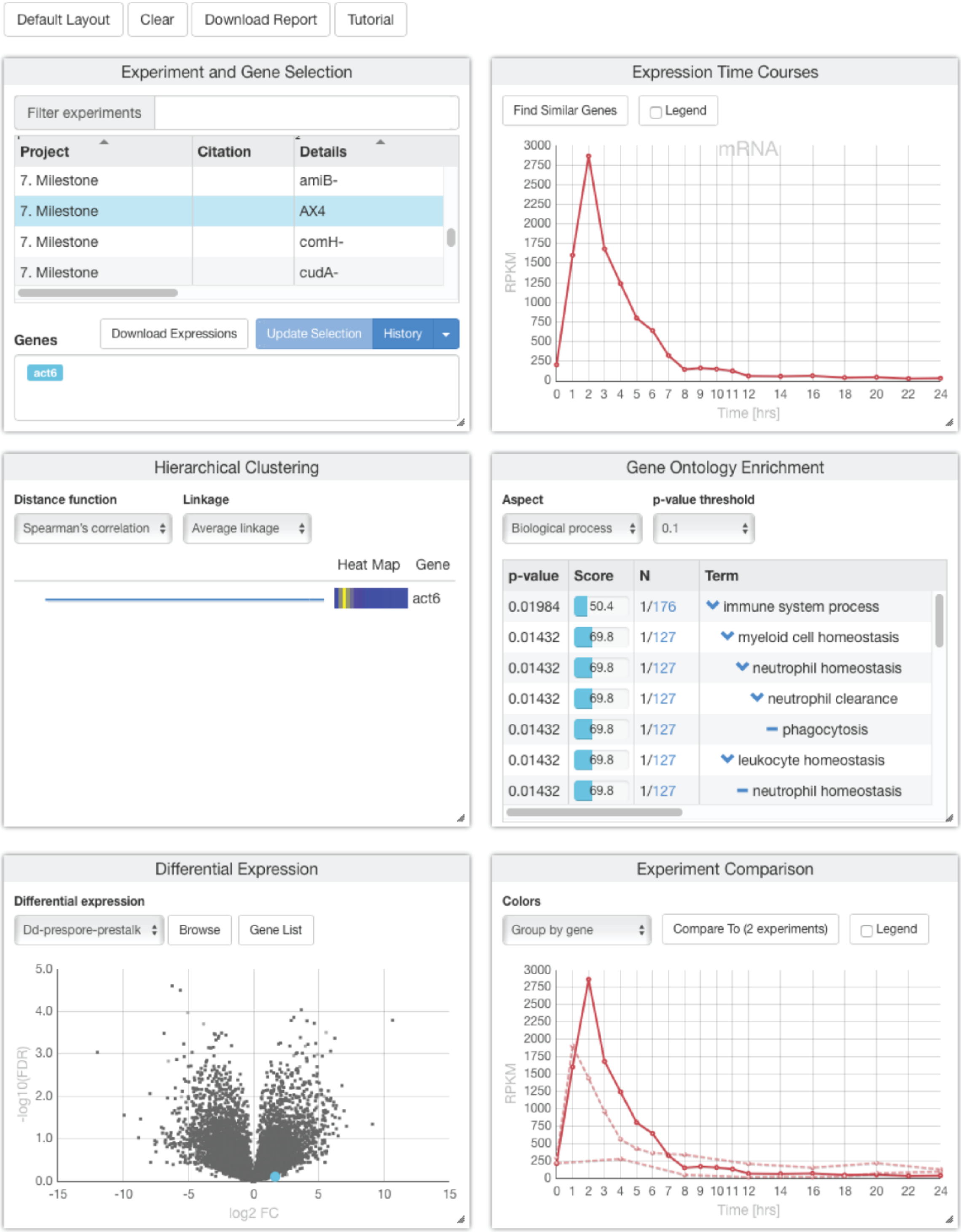

Change the color selection by selecting the desired grouping in the ‘Group by’ button. Add a legend as needed. Mouse over the legend to view details (not shown).
It is easy to see that the actin 6 mRNA abundance is reduced to about 60% in the aggregationless *acaA*$^{-}$ strain (tan) and to about 10% in the precocious *pkaC*$^{OE}$ strain (purple).

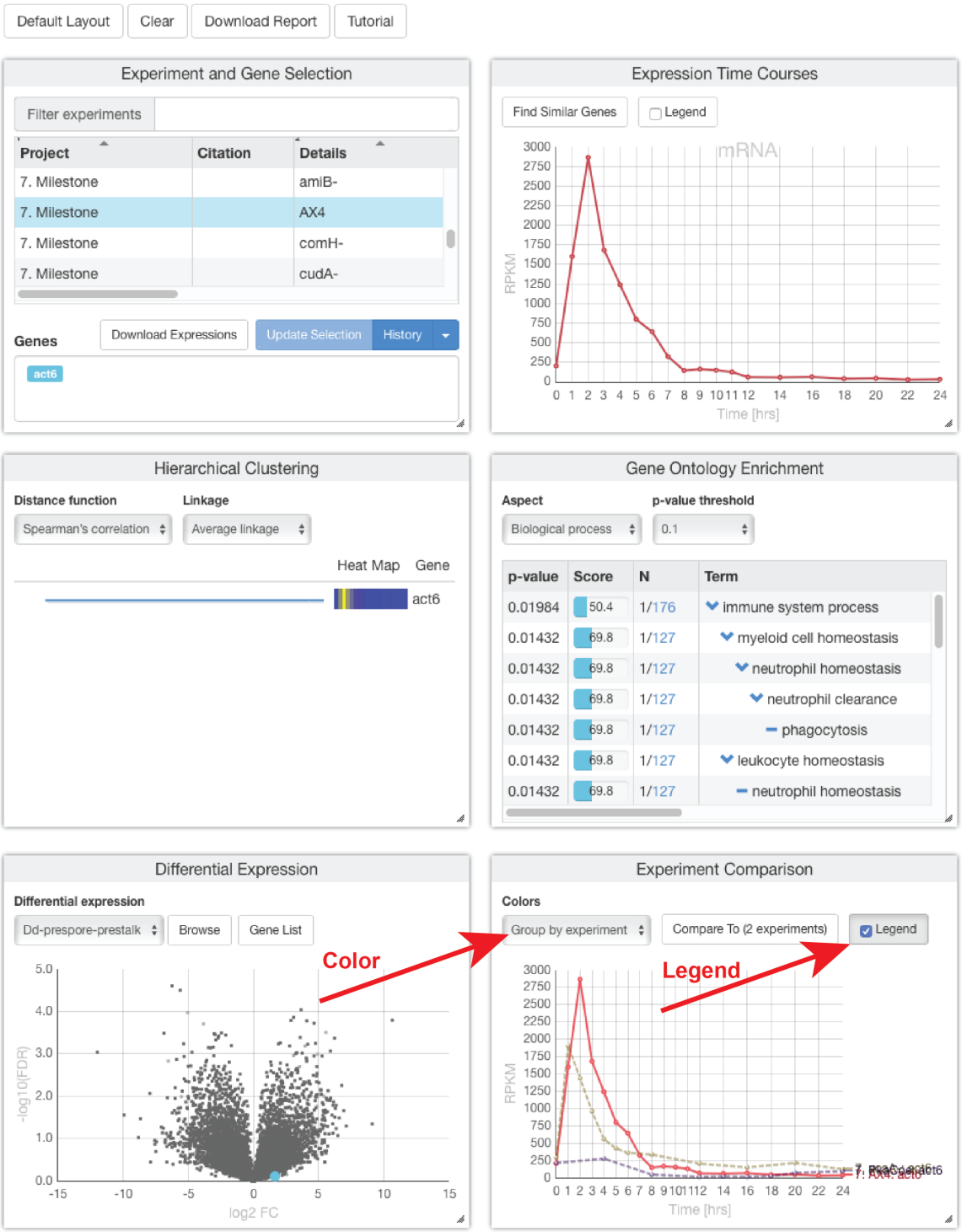

#### B. An introduction to Orange

Install Orange on your computer https://orangedatamining.com. If you are new to Orange, we strongly recommend following the 'Getting Started with Orange' tutorials at https://www.youtube.com/channel/UClKKWBe2SCAEyv7ZNGhIe4g
Then, add the Bioinformatics add-on as described in https://www.youtube.com/watch?v=OANsA6fMJKg. In the following introduction, we used Orange 3.27.1 with the Bioinformatics add-on 4.3.1

First, select two data sets for comparison. Start by opening a new Orange canvas.

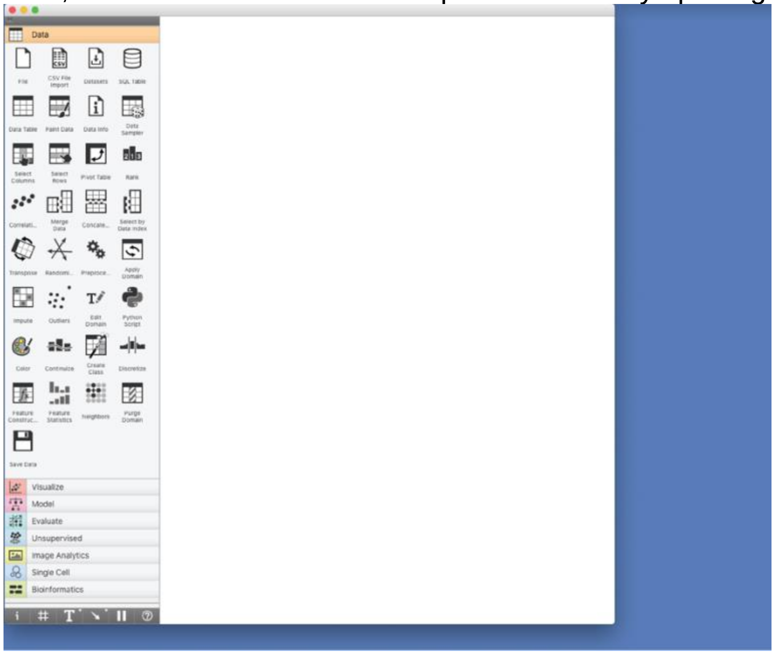

Select the dictyExpress widget from the Bioinformatics menu.

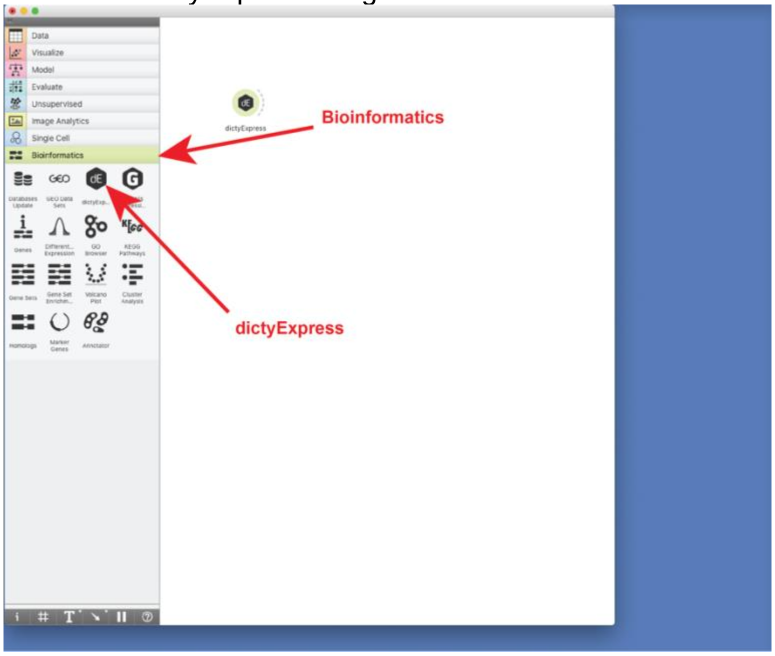

Open the dictyExpress widget (double click).

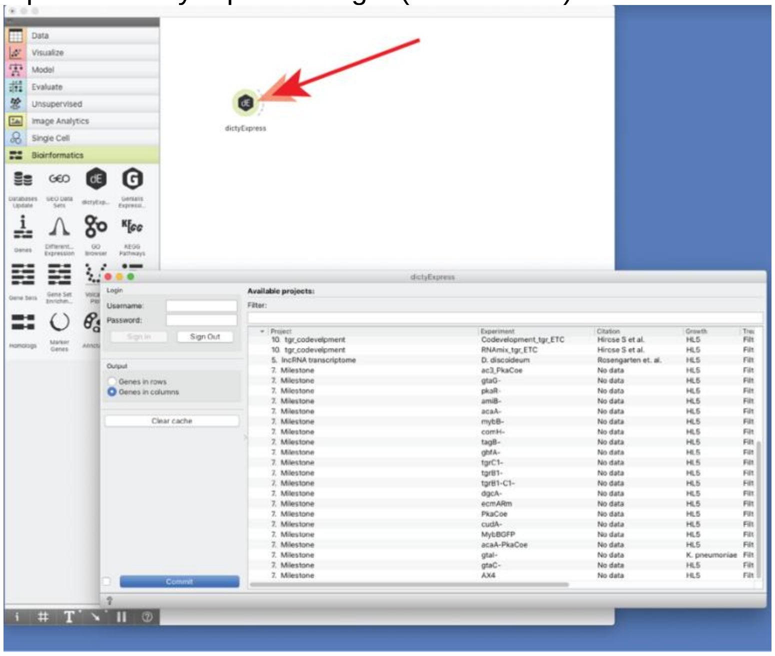

Select the ‘Genes in columns’ output and one of the Milestone datasets – a dot will appear next to the selection. Here, we selected AX4. Press ‘Commit’ to confirm your selection and close the dictyExpress window.

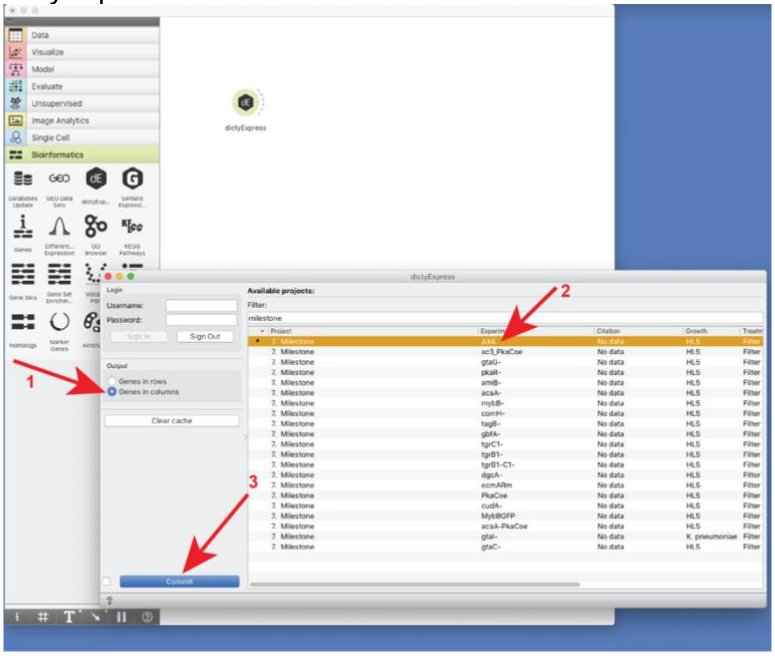

Select a second dictyExpress widget from the Bioinformatics menu and repeat the above process but select a different dataset for comparison. Here, we selected mybB-.

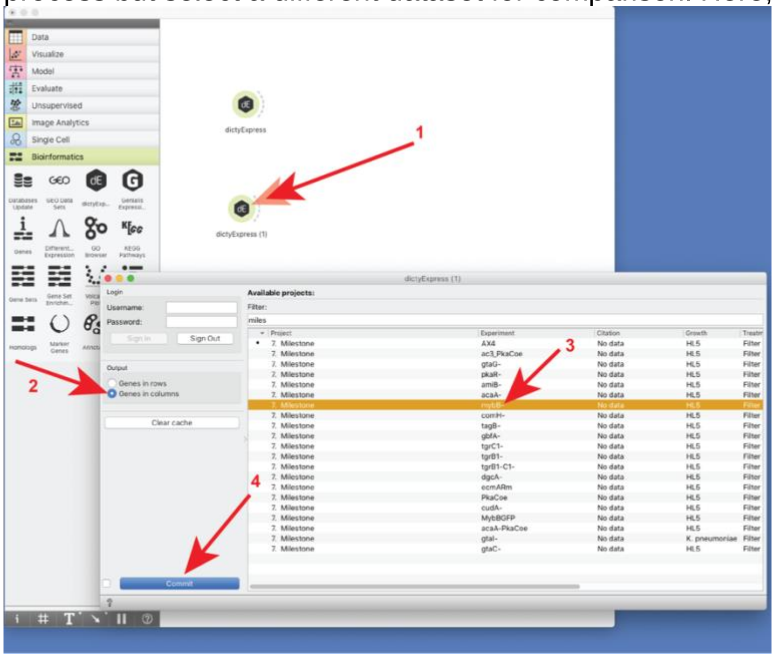

Next, we will combine the two datasets. Select the Concatenate widget from the Data menu.

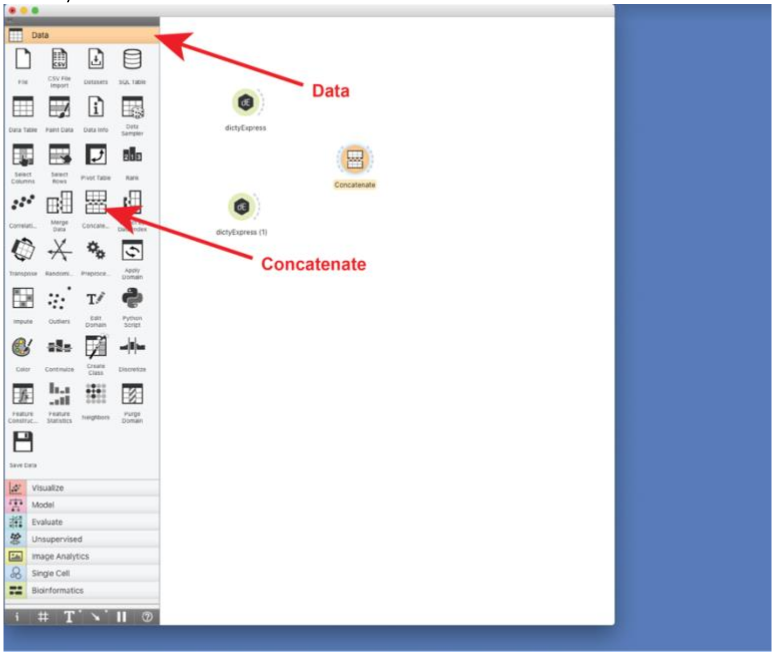

Connect the outputs of the two dictyExpress widgets to the Concatenate widget and open the widget (double click).

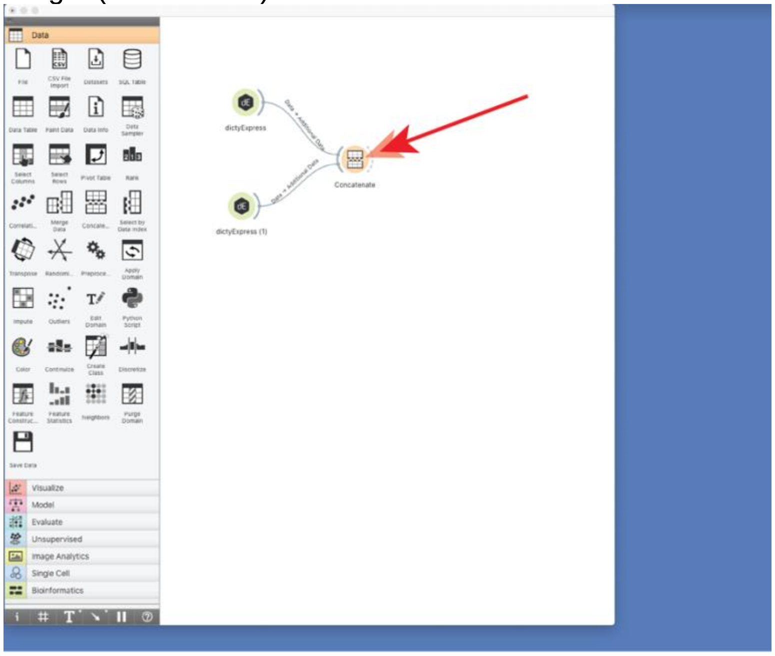

Select the variable merging and source identification options as shown. Here, we changed the source ID by typing ‘Genotype’ in the feature name box. Notice that this widget is set to apply the selections automatically by default.

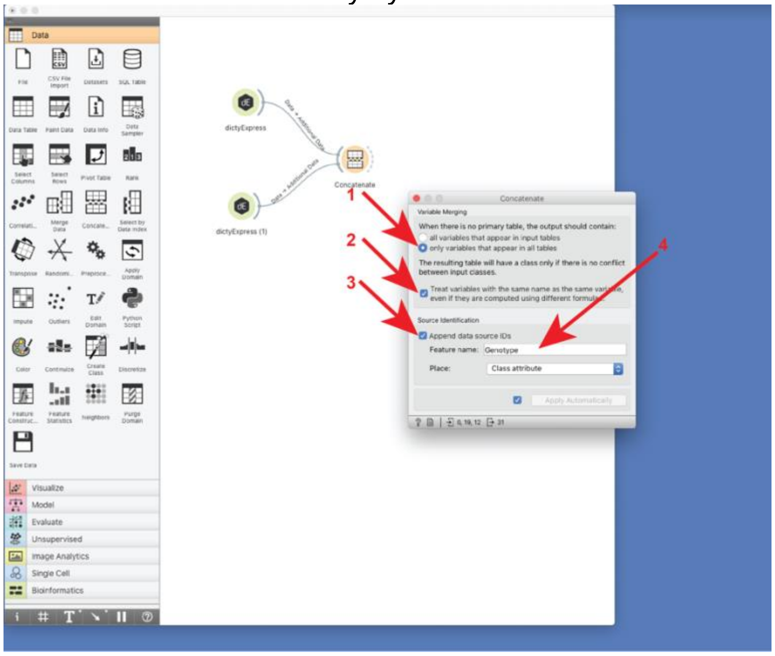

The next 5 steps will change the genotype labels to indicate the strain names in the output data. Add an Edit Domain widget from the Data menu.

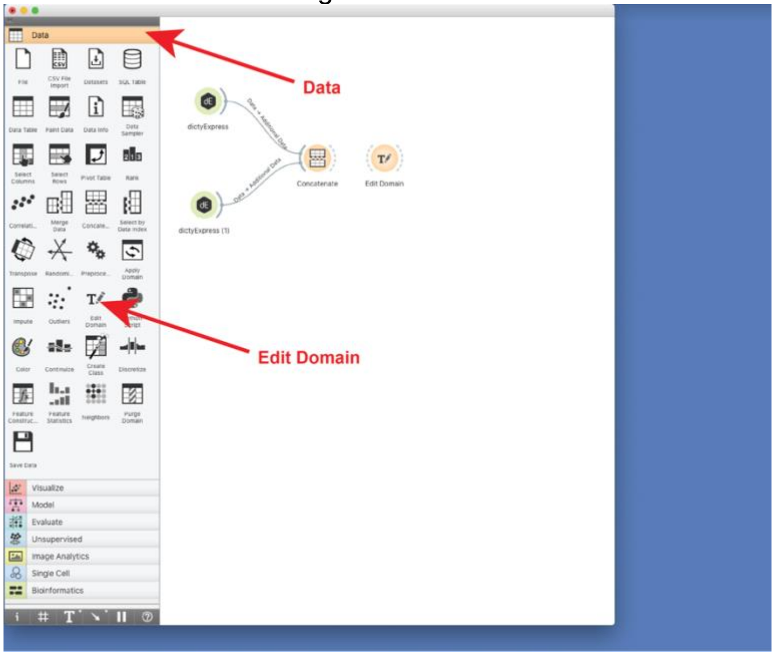

Connect the output of the Concatenate widget to the Edit Domain widget and open the Edit Domain widget (double click).

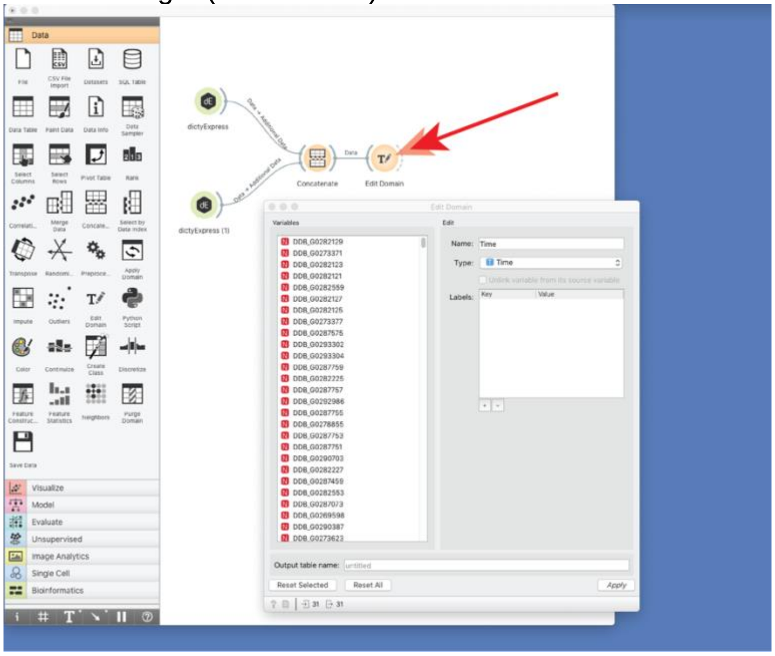

Scroll down the ‘Variables’ list and click the ‘Genotype’ variable.

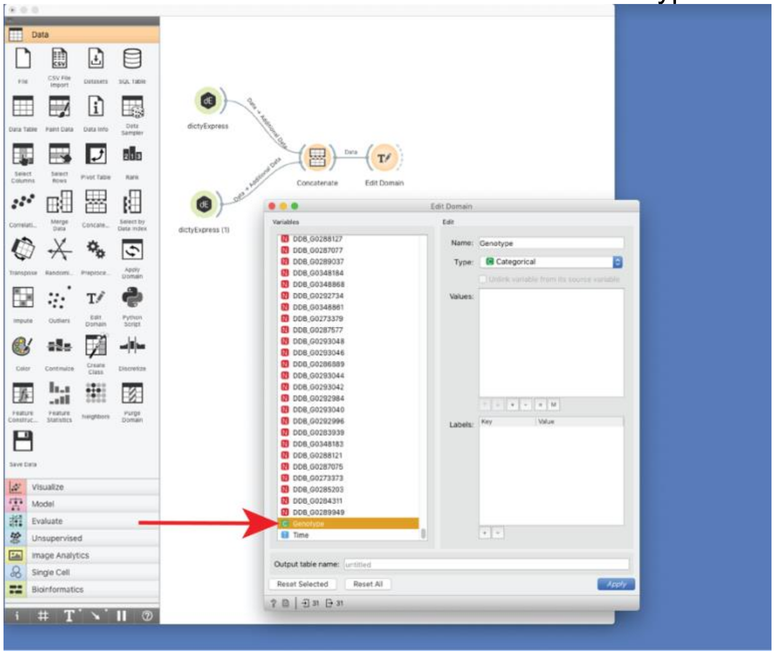

Click inside the ‘Values’ box. Text will appear if it is not already there.

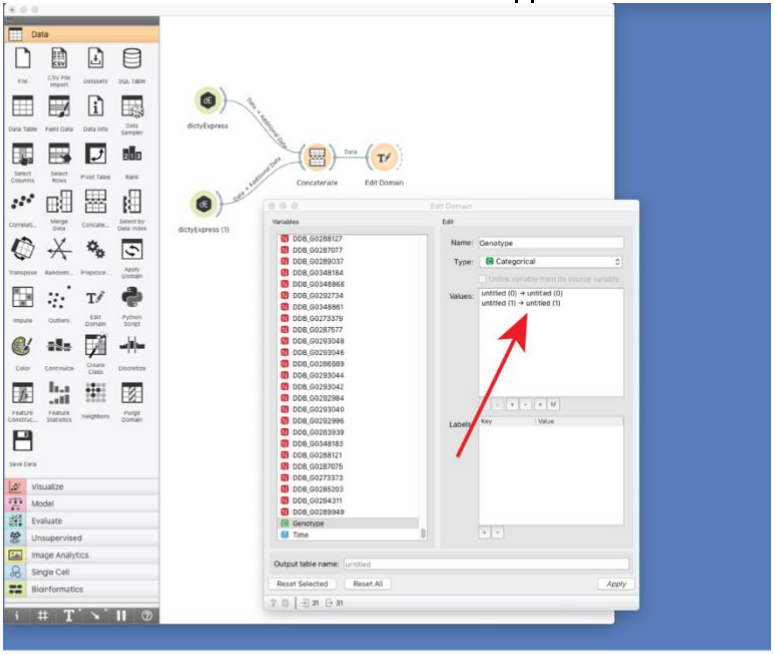

Double click the first value and type ‘AX4’.

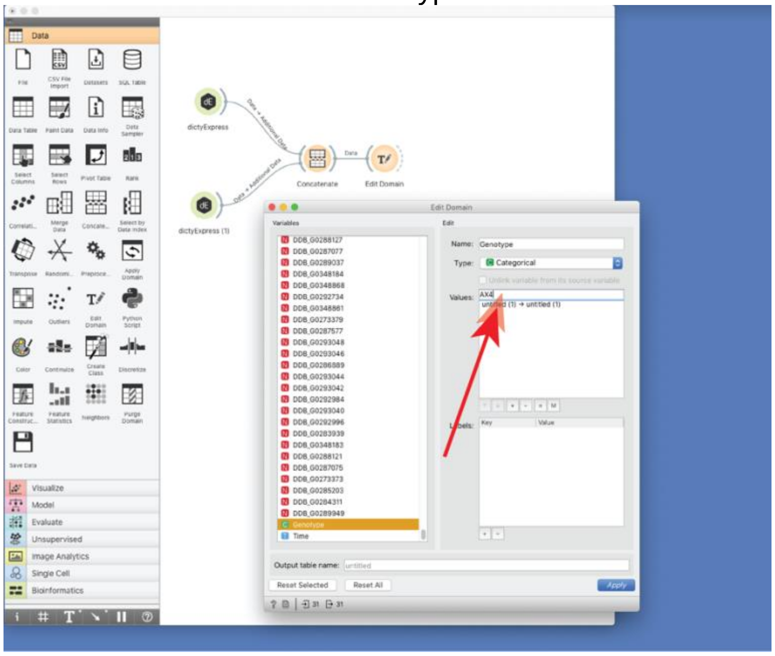

Double click the second value and type ‘mybB-‘. Press ‘Apply’ to propagate the two changes and close the Edit Domain window (not shown).

To view your combined data, connect a Data Table widget to the Edit Domain output and double click to view the data. Here we show the first few columns that list the Genotype, Time and the first three of the 12828 genes (features) in the dataset.

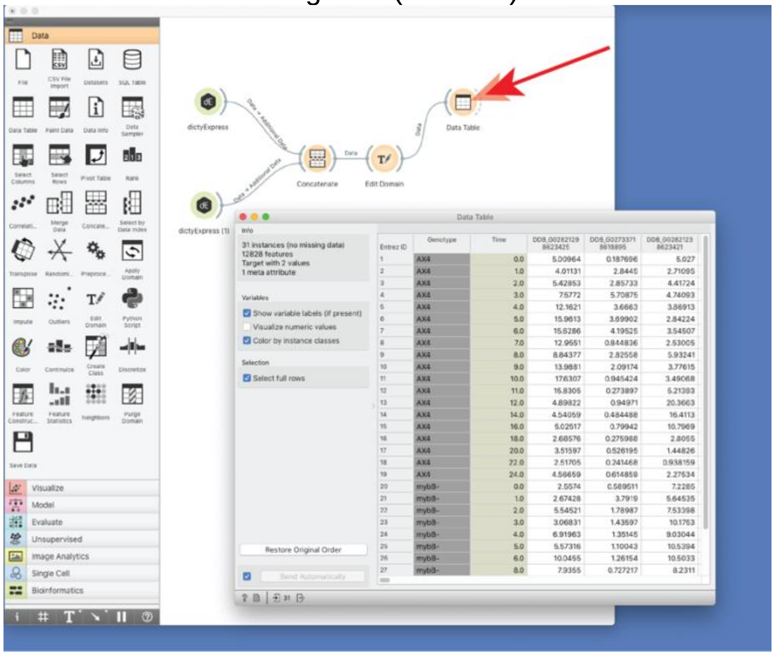

To compare the two datasets, select the MDS (multidimensional scaling) widget from the Unsupervised menu.

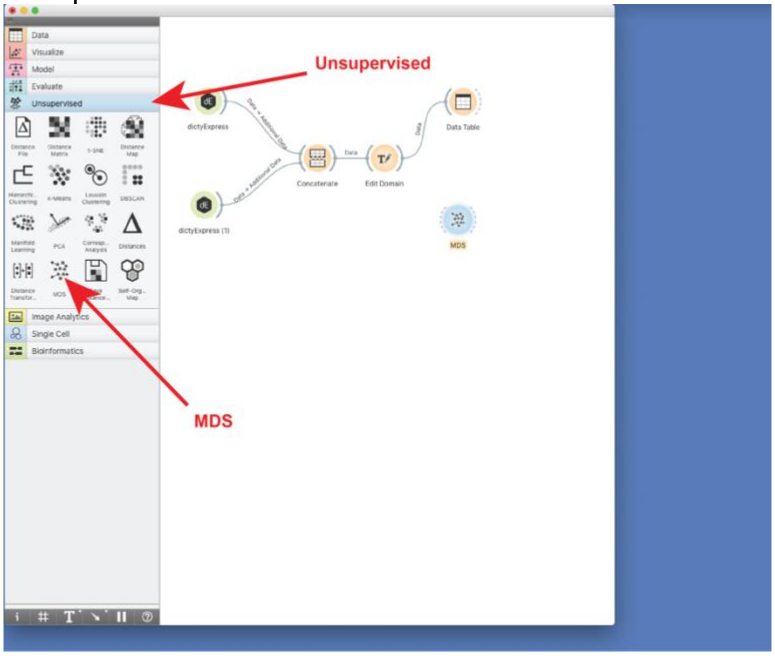

Connect the output of the Edit Domain widget to the MDS widget. Red dots will appear at the ends of the connecting line, indicating data processing. Open the MDS widget (double click). Using the interactive menu on the left, 'Color' the points by 'Genotype' and to 'Label' them by 'Time'. Reduce the scale of the "Show similar pairs" to produce an MDS plot as shown.

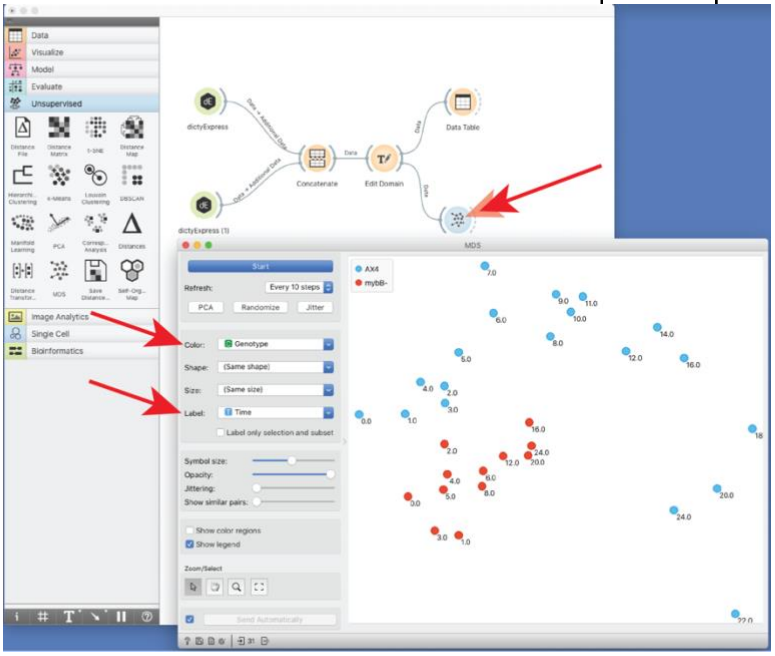

The wild type (AX4) temporal progression is quite different from that of the aggregationless mutant (mybB-), similar to the data shown in Figure 2A in the main manuscript. The projection is not identical to Figure 2A because the latter contains additional datasets that affect the rendering. You could test your skills by changing the mutant dataset from mybB- to another mutant (suggestion: tagB-; not shown).

**Supplemental_File_S5: Computational methods**

**Abbreviations:**
lFC – log fold change
DE – differential expression/differentially expressed
padj – adjusted p-value
pval – p-value
RPKUM – Read counts Per Kilobase of exon model per Uniquely mapped Million reads
tt – transition time
agg – aggregation
FB – fruiting body
GAM – generalized additive model

#### 1 Milestones

To find genes that change their expression relatively strongly between two consecutive developmental stages, we filtered the genes based on DE between two stages and also the shapes of their expression profiles. This approach ensured that the milestone gene exhibited a strong change between two stages and did not fluctuate much during the rest of development. For this analysis, we used AX4 samples annotated with developmental stages based on images that captured the morphologies of developmental structures. When an image contained multiple morphological stages, the majority morphology was used.

##### 1.1 Genes with marked expression changes between stages

First, DE analysis was performed with the DESeq2 (v1.26.0) R library (Love et al. 2014) for every pair of neighboring stages using the AX4 samples. The later stage was used as the case and the earlier as the control. The padj was re-calculated over all the tests for all neighboring stage pairs using Benjamini-Hochberg correction on the pval from DESeq2. A gene was considered DE if the absolute lFC ≥ 2 and padj ≤ 0.01.

##### 1.2 Genes differentially expressed across stages

The shape of the expression profile was analyzed with the ImpulseDE2 (v1.10.0) R library (Fischer et al. 2018) to find DE genes whose expression profiles change monotonously or transiently during development. The ImpulseDE2 model was fit to the AX4 data using ordered majority stages converted to consecutive integers as time-points. Transition times were defined as the x-coordinate values of the sigmoid midpoints. The analysis was run in case-only mode with identification of transient genes whose expression profiles are better fit by single or double sigmoid model compared to constant model. Genes were considered significantly DE between two stages based on padj threshold = 0.001 and with a tt between those stages.

The fitted ImpulseDE2 models were parsed to obtain neighboring stages that had tt between them, indicating a change in the expression level. The tt values were extracted from the appropriate model based on genes that were termed as monotonously or transiently changed across stages by ImpulseDE2. In some cases, this would lead to an inappropriate assignment of tt owing to the input timescale range (stages) and model complexity (single or double sigmoid). For each type of tt that would lead to an inappropriate expression change assignment (described below), changes were made to the tt values based on visual evaluation of example genes with the same tt value inconsistency.

We extracted a single tt for monotonously DE genes according to the following steps even when the double sigmoid impulse model (i.e. having two tt values) was used. If the padj of the impulse model was lower than that of the sigmoid model (single sigmoid), the impulse tt closer to the tt of the sigmoid model was chosen. Otherwise, the tt of the sigmoid model was used. For border reassignment, if a tt value was smaller than the x value of the first stage (no agg), it was readjusted to be immediately after the first stage. If a tt was larger than the x-value of the last stage (FB), it was readjusted to be immediately before the last stage.

For transiently DE genes, two tt values were extracted from the double sigmoid model. If both tt values were between the same two stages, they were reassigned to be right before the first neighboring stage and right after the second neighboring stage. If tt values were smaller than the first stage (no agg) or larger than the last one (FB), they were reassigned as in the monotonous model. If this procedure set the two tt values to be between the same two stages, a single tt value between these two stages was extracted.

#### 1.3 Selection of milestone genes

A gene was determined to be a milestone between two neighboring stages by two criteria; 1) the gene was significantly DE between these two stages based on DESeq2 (Section 1.1) and 2) it was significantly DE based on ImpulseDE2 with tt between the two stages (Section 1.2). These milestone genes were then separated based on being up- or down-regulated between the two stages.

#### 1.4 Milestone genes - expression heatmaps

The expression profiles of milestone genes were visualized with the ComplexHeatmap (v2.3.3) R library (Gu et al. 2016). RPKUM data were averaged across the multiple samples, which were annotated as the same majority stage in each strain, so that the expression data of each gene were summarized by a single averaged value for each stage of a strain. The expression was scaled based on the following formula:

$$Gist\ scaled = \frac{Gist - p_{99}(Gi)}{p_{99}(Gi)}$$

where *Gi* represents all the averaged expression values of a given gene (*i*) across stages and strains, *Gist* represents *Gi* in a certain strain (*s*) at a certain stage (*t*), and $p_{99}$ represents the 99$^{th}$ percentile. This formula linearly scales the majority of the *Gist* values to the interval [-1,0] with extremely high *Gist* being given a value above zero. Values were then capped at 0.1 to reduce the effect of extreme values on the color scale.

The heatmaps of the milestone genes were prepared separately for each pair of neighboring stages and for up- and down-regulation. Gene ordering was based on hierarchical clustering of the scaled averaged AX4 data with the Ward algorithm (ward.D2) in hclust (R v3.6.3) using Euclidean distances, followed by visual reordering with the seriation (v1.2-8) R library (Hahsler et al. 2008). The two strains that did not have stage annotations (*ac3*$^{–}$*/pkaCoe* and *gtaC*$^{–}$) were not included in the milestones heatmap.

#### 2 Regulons

##### 2.1 Selection of regulon candidate genes

Co-regulated gene pairs were extracted from individual strain data to avoid biasing in favor of strains with more samples, according to the following steps. First, we excluded genes whose RPKUM was all-zero in a strain. The RPKUM values of each gene were transformed by adding a pseudocount (+1) followed by $log_2$ transformation and scaling to a mean = 0 and standard deviation = 1. We used the Python nearest neighbor descent, PyNNDescent (v0.3.3) library (https://libraries.io/pypi/pynndescent/0.3.3), to obtain the 300 nearest neighbors of each gene based on cosine similarity. Then, we chose all the genes that have at least one nearest neighbor that exhibited a similarity equal to or higher than the strain-specific threshold (the $30^{th}$ percentile of the similarities in each strain; see 2.2 Strain-specific similarity threshold). For each gene, we counted the number of strains in which the gene was found to have some co-expressed neighbor(s) and compared the number with the gene-specific N threshold. The gene-specific N was specified by the number of strains in which the gene was deemed as expressed highly enough (see details below). If the gene had closest neighbor(s) present in at least N strains, it was considered a regulon candidate. Some genes were strongly co-regulated only in a few strains and exhibited mainly low or no expression in most strains, where they were not counted as co-regulated. When the expression level of a gene pair is very low, cosine similarity often becomes lower than the true similarity due to amplifying noise. Therefore, even if the genes are co-regulated, they would be counted as negative. To avoid such false negatives, we lowered the gene-specific N threshold by not taking into account strains in which the gene was expressed at very low levels. Thus, we first determined the H percent of the $99^{th}$ percentile expression (*He*) in all samples and then defined the gene-specific N threshold as the number of strains in which the gene's expression reached its *He* value at any timepoint. We also tested different H values (0, 10, 30, or 50) for the expression level. By increasing H (resulting in lowering N) more genes would be included as regulon candidates, because genes can be co-expressed even in strains that exhibit relatively low expression (empirically, we found H=0, 365 genes; H=10, 1099 genes; H=30, 1974 genes; H=50, 3182 genes). We chose H=10 based on a visual inspection of the regulons obtained with different values of H. Moreover, we also set an upper limit of N=18 instead of 21 (all strains) to avoid more false negatives in the extraction process for the co-expressed gene pairs. The cap N=18 ensures that a gene would have co-expressed close neighbor(s) in at least one strain among the aggregation minus group in which regulons were frequently disrupted.

##### 2.2 Strain-specific similarity threshold

A gene profile similarity threshold was selected to classify genes as co-expressed or not. The gene profile similarities depend on the strain-specific number of samples and data quality. A different similarity threshold was thus selected for each strain. These similarities to the top closest neighbors obtained for each strain displayed a left-skewed distribution. A strain-specific similarity threshold was set to the $30^{th}$ percentile of the similarities to the closest neighbors. This threshold approximately separated the closest neighbor similarity distribution to a bulk of genes with close neighbors and a tail of genes that had relatively low similarity to the closest neighbor.

##### 2.3 Clustering of selected genes into regulons

The selected regulon candidate genes were clustered based on their expression profiles. Clustering and data preprocessing were performed in Orange (v3.26) (Demsar et al. 2013). We used two methods for data preprocessing: 1) We added a pseudocount (+1) to the RPKUM data, $log_2$-transformed and scaled to mean = 0 and standard deviation = 1. We used the scaled expression of samples as features for clustering. 2) We scaled the RPKUM data of each gene to interval [0,1], followed by PCA dimensionality reduction. We used the first 30 PCA components as features for clustering. Based on visual evaluation of the clustering results, we selected the first method for AX4-based regulons. Most of the regulon candidate genes are expressed in AX4 and thus the first method performed better on the AX4 data as it was able to capture more subtle changes in expression. On the other hand, the second method gives more importance to higher expression profiles, mostly due to peaks, and less importance to lower expression values than the first method. Thus, the second method performed better on strain-wide data where many strains do not express a gene leading to relatively more noise in their low expression values. We selected the second method for all-strains-based regulons. Louvain clustering was performed with resolution 0.8 when AX4-based regulons were extracted from the AX4 data only and with resolution 0.4 for all-strains-based regulon extraction.

##### 2.4 Regulons expression heatmap

Regulon heatmap were prepared as for Milestone genes with the following changes. RPKUM data were averaged across timepoints of the replicates of each strain. Regulons were ordered based on the developmental time of the gene expression peak in AX4. Peak times of individual regulon genes were obtained from the averaged non-scaled AX4 data. Regulons were ordered first by the median of the peak times, followed by the mean of the peak times of the regulon genes. Genes were ordered within each AX4 regulon separately. The ordering was based on hierarchical clustering of the scaled averaged AX4 data as for the milestone genes. The genes in the heatmap of the 'all-strains' regulons were ordered based on the AX4 order, first by regulons and then by ordering within the regulons.

#### 3 Disaggregation genes

We performed DEseq2 analysis to select genes that are related to the disaggregation process in the *tgr* mutant strains. The relevant timepoints were selected based on visual evaluation of the differences in PC1 values between *tgrB1*⁻ and *tgrB1*⁻*C1*⁻ and the AX4, *tagB*⁻ and *comH*⁻ strains.

##### 3.1 Selection of disaggregation genes

Genes that were upregulated during disaggregation, but not upregulated at the same time in normal development were extracted with the following method. Genes DE in individual strains between time points: 6 and 8 hrs, and 8 and 12 hrs, were extracted for AX4, *tagB*, *comH*, *tgrB1*⁻, and *tgrB1*⁻*C1*⁻ strains with DESeq2. The design used adjustment for replicates and thus only replicates present at both timepoints were used. The DESeq2 results were optimized for padj threshold 0.01. A gene was considered significantly upregulated if lFC ≥ 1.32 and padj ≤ 0.01. For each time comparison, the genes upregulated during disaggregation in both *tgrB1*⁻ and *tgrB1*⁻*C1*⁻, but not in AX4, *tagB*⁻, or *comH*⁻ were selected as disaggregation genes.

#### 3.2 Comparison between the disaggregation and dedifferentiation genes

To characterize the selected disaggregation genes, they were compared with genes that are upregulated during early dedifferentiation. The dedifferentiation genes were obtained from published dedifferentiation RNA-seq data (Nichols et al. 2020). The published data included an experiment in which cells were disaggregated and incubated in nutrient medium to induce dedifferentiation, and a control in which the disaggregated cells were incubated in non-nutrient buffer. We downloaded the RNA-seq fastq files (GSE144892) and prepared the RPKUM data by the same procedure as ours through the Genialis platform. Dedifferentiation genes were selected based on a DESeq2 comparison between the pooled ‘medium’ samples at 0.5, 1, and 2 hrs and the pooled ‘buffer’ samples at 0, 0.5, 1, 2, 3, 4, and 6 hrs. We tested for upregulation with DeSeq2 and optimized the results for padj threshold 0.01. A gene was considered to be upregulated during dedifferentiation if lFC ≥ 2 and padj ≤ 0.01. Gene expression scaling and gene ordering for the heatmaps were performed as for the milestone genes. We used a hypergeometric test to determine whether the disaggregation genes significantly overlap with the dedifferentiation genes. For the reference group in the test we used all genes expressed in the data published with this paper.

### 4 Developmental stage annotation

We prepared two types of stage annotations: 1) all stages annotations and 2) representative stages annotations. First, we manually annotated developmental stages from the microscopic images showing developmental morphological structures. When an image contained multiple morphological stages, we annotated the sample with all the observed structures. If the image was not captured for any sample, it was annotated as “no image”, except for t=0 where it was annotated as no_agg. All stage annotations are shown in the color pallet above the heatmaps of regulons and disaggregation genes as information for each sample. When selecting milestone genes that transcriptionally define each developmental stage boundary, we annotated each sample with a “representative stage annotation” that is characteristic of the most abundant morphology.

### 5 Gene-set enrichment analysis

Datasets used for gene-set enrichment (including dictyBase gene name – entrez ID mapping and gene sets) were obtained on the 5$^{th}$ of April, 2020. The data were collected from the following sources. Gene information and taxonomy data were obtained from the NCBI database (ftp://ftp.ncbi.nlm.nih.gov). Gene ontology and KEGG pathway information was obtained from the official GO (http://geneontology.org/docs/download-ontology/) and KEGG (https://www.genome.jp/kegg/) knowledge-bases, respectively. The gene sets were pre-processed to contain only genes with Entrez IDs. The code used for data retrieval is available at https://github.com/JakaKokosar/bioinformatics-serverfiles and the data used in this study are stored at http://download.biolab.si/datasets/bioinformatics/2020_04_05/.
The analysis was performed using the Orange Bioinformatics (v4.0.0) Python library. Entrez IDs were mapped to the dictyBase gene names. The following gene sets were used: generic GO slims for biological process, molecular function, and cellular

component; KEGG Pathways; and the custom gene sets described below. Only gene sets with size within the interval [5,500] were used. The size was determined based on the number of genes that were contained within the reference set. The reference set contained all the genes present in the RNA-seq data, except those that had all zero expression values. Reference and query sets were filtered to include only genes that had an Entrez ID and were contained within at least one of the used gene sets. The latter was done to account for different proportions of genes annotated with a gene set between the reference and the query groups. All gene sets that had non-zero overlap with the query were tested for enrichment using the Orange Gene Set set_enrichment function. The pval were adjusted with Benjamini-Hochberg correction. Results were filtered to display only gene sets with padj ≤ 0.25 and overlap with query ≥ 2.

##### 5.1 Custom gene sets

We used cell-type specific genes (Prespore and Prestalk genes) (Parikh et al. 2010), cAMP-pulse induced genes (Iranfar et al. 2003), chemotaxis genes (Swaney et al. 2010), *Dictyostelium* short gene families (*hssA/2C/7E* family, 57-aa protein family, *sig* and sigN genes, and *gtaG*-dependent short proteins) (Shimada et al. 2008; Vicente et al. 2008; Katoh-Kurasawa et al. 2016) and transcriptional regulation and chromatin organization (regulatory transcription factor, general transcription factors, mediators, chromatin remodeling/histone modification, histone/histone variants, and chromatin/centromere) (Rosengarten et al. 2013; Forbes et al. 2019). These custom gene sets are provided in Supplemental_File_S8.

**Supplemental_File_S6: Standard experimental methods**

**Cell culture, strain maintenance, development and spore collection**

All the *Dictyostelium discoideum* strains were derivatives of AX4 (Knecht et al. 1986) as detailed in Supplemental_Table_S1. We cultured cells at 22°C in HL5 medium with the necessary supplements and antibiotics. To induce development, we washed exponentially growing cells twice with KK2 buffer (20 mM potassium phosphate, pH6.4) to remove nutrients. Cells of the *gtaI*$^{-}$ strain were grown in association with live *Klebsiella pneumoniae* bacteria on SM plates, collected at the exponential growth phase, and washed at least three times with DDW. In all cases, we deposited the cells at a density of $2.6 \times 10^6$ cells/cm$^2$ on black nitrocellulose filters on top of a paper pad soaked with PDF buffer (20 mM KCl, 9.2 mM $K_2HPO_4$, 13.2 mM $KH_2PO_4$, 5.3 mM $MgCl_2$ and 1 mM $CaCl_2$, pH6.4). The cells were incubated in the dark at 22°C for defined periods of time (Katoh et al. 2004). We collected spores from developing structures and treated with detergent (0.1% NP-40, 1mM EDTA in KK2 buffer) to eliminate amoebae (Shaulsky and Loomis 1993).

**RNA-seq**

We collected the cells from one nitrocellulose filter at each time point of two to seven independent developmental series, extracted total RNA from each sample using 1 ml of Trizol (Invitrogen) and performed poly(A) selection twice as described (Katoh-Kurasawa et al. 2016). We note that fruiting bodies, which contain walled spores and stalk cells, were not broken mechanically prior to RNA extraction, which could have resulted in underrepresentation of RNA species that are found exclusively in these walled cells (Van Driessche et al. 2005). We prepared multiplexed cDNA libraries and performed RNA sequencing using the Illumina sequencing platform as described previously. We mapped the resulting sequences to the *Dictyostelium* reference genome and obtained mRNA abundance values for each gene in the genome (Miranda et al. 2013) through the web applications dictyExpress, GenBoard or Genialis platform. The data were deposited in GEO (accession numbers GSE152851). Unless otherwise stated, we preprocessed RPKUM data by $log_2$-transformation after adding one (pseudocount), and then scaling to mean = 0 and standard deviation = 1.

**Supplemental Files S7 and S8 are provided as separate Excel spreadsheets.**

Supplemental_File_S7 Milestones, regulons and disaggregation gene lists
Supplemental_File_S8 Reference gene lists

#### Supplemental_Table_S1 *D. discoideum* strains used

| Strain name | Phenotype group | Strain Descriptor | Strain summary | Parental strain | Antibiotic resistance | Reference |
|---|---|---|---|---|---|---|
| AX4 | wild type | AX4 | wild type | AX3 | | (Knecht et al. 1986) |
| MybBGFP | wild type | mybB–/[mybB]:mybB:GFP | expressing MybB:GFP under control of *mybB* promoter in *mybB*— | *mybB*$^{-}$ | Blasticidin S, G418 | this study |
| pkaC$^{oe}$ | precocious development | [act15]:pkaC:HA | expressing PkaC under control of *act15* promoter in AX4 | AX4 | G418 | this study |
| *pkaR*— | precocious development | *pkaR*— | *pkaR* null mutant | AX4 | Blasticidin S | (Shaulsky et al. 1998) |
| *acaA*—/ *pkaC*$^{oe}$ | small fruiting body | *acaA*— /[act15]:pkaC | expressing PkaC under control of *act15* promoter in *acaA*$^{-}$ | *acaA*$^{-}$ | Blasticidin S, G418 | this study |
| *ac3*—/ *pkaC*$^{oe}$ | small fruiting body | *acaA—/acgA— /acr A—/* [act15]:pkaC:HA | expressing PkaC under control of *act15* promoter in *acaA—/acgA— /acrA—* | *acaA*$^{-}$ /*acgA*$^{-}$ /*acrA*$^{-}$ | Blasticidin S, G418 | this study |
| *gtaI*— | culmination defective | *gtaI*— | *gtaI* insertional mutant | AX4 | Blasticidin S | this study |
| *gtaG*— | culmination defective | *gtaG*— | *gtaG* insertional mutant | AX4 | Blasticidin S | (Katoh-Kurasawa et al. 2016) |
| *cudA*— | culmination defective | *cudA*— | *cudA* deletion mutant pcudAKO(MF2) | AX4 | Blasticidin S | this study; Plasmid: (Fukuzawa et al. 1997) |
| *dgcA*— | culmination defective | *dgcA*— | *dgcA* null mutant pdgcA_KO_443 | AX4 | Blasticidin S | this study; Plasmid: (Chen and Schaap 2012) |
| *ecmARm* | culmination defective | [ecmA]:pkaR (G135E/G261A) | *gtaG* insertional mutant | AX4 | G418 | this study |
| *tagB*— | tag arrest | *tagB*— | *tagB* null mutant | AX4 | Blasticidin S | (Khare and Shaulsky 2010) |
| *t345* | tag arrest | *comH*— | *comH* insertional REMI mutant | AX4 | Blasticidin S | (Kibler et al. 2003)[g] |
| *tgrB1*— | tag disaggregation | *tgrB1*— | *tgrB1* null mutant | AX4 | Hygromycin B | (Benabentos et al. 2009) |

| | | | | | | |
|---|---|---|---|---|---|---|
| *tgrB1C1*$^{-}$ | tag disaggregation | *tgrB1*$^{-}$/*tgrC1*$^{-}$ | *tgrB1C1* null mutant | pyr5-6- | Uracil | (Hirose et al. 2011) |
| *tgrC1*$^{-}$ | lag disaggregation | *tgrC1*$^{-}$ | *tgrC1* null mutant | AX4 | Hygromycin B | (Benabentos et al. 2009) |
| *gbfA*$^{-}$ | lag disaggregation | *gbfA*$^{-}$ | *gbfA* null mutant | AX4 | BS- | this study |
| *gtaC*$^{-}$ | aggregation minus | *gtaC*$^{-}$ | *gtaC* deletion mutant | AX4 | Blasticidin S | (Keller and Thompson 2008) |
| *acaA*$^{-}$ | aggregation minus | *acaA*$^{-}$ | *acaA* null mutant | AX4 | Blasticidin S | (Stepanovic et al. 2005) |
| *mybB*$^{-}$ | aggregation minus | *mybB*$^{-}$ | *mybB* insertional mutant | AX4 | Blasticidin S | this study |
| *amiB*$^{-}$ | aggregation minus | *amiB*$^{-}$ | *amiB* insertional mutant p82ClaI_amiB-REMI | AX4 | Blasticidin S | this study; Plasmid: (Kon et al. 2000) |

#### References

Benabentos R, Hirose S, Sucgang R, Curk T, Katoh M, Ostrowski EA, Strassmann JE, Queller DC, Zupan B, Shaulsky G et al. 2009. Polymorphic members of the lag gene family mediate kin discrimination in Dictyostelium. *Current biology : CB* **19**: 567-572.

Chen ZH, Schaap P. 2012. The prokaryote messenger c-di-GMP triggers stalk cell differentiation in Dictyostelium. *Nature* **488**: 680-683.

Demsar J, Curk T, Erjavec A, Gorup C, Hocevar T, Milutinovic M, Mozina M, Polajnar M, Toplak M, Staric A et al. 2013. Orange: Data Mining Toolbox in Python. *Journal of Machine Learning Research* **14**: 2349-2353.

Fischer DS, Theis FJ, Yosef N. 2018. Impulse model-based differential expression analysis of time course sequencing data. *Nucleic Acids Res* **46**: e119.

Forbes G, Chen ZH, Kin K, Lawal HM, Schilde C, Yamada Y, Schaap P. 2019. Phylogeny-wide conservation and change in developmental expression, cell-type specificity and functional domains of the transcriptional regulators of social amoebas. *BMC genomics* **20**: 890.

Fukuzawa M, Hopper N, Williams J. 1997. cudA: A Dictyostelium gene with pleiotropic effects on cellular differentiation and slug behaviour. *Development* **124**: 2719-2728.

Gu Z, Eils R, Schlesner M. 2016. Complex heatmaps reveal patterns and correlations in multidimensional genomic data. *Bioinformatics* **32**: 2847-2849.

Hahsler M, Kurt H, Christian B. 2008. Getting Things in Order: An Introduction to the R Package seriation. *Journal of Statistical Software* **25**: 1-34.

Hirose S, Benabentos R, Ho HI, Kuspa A, Shaulsky G. 2011. Self-recognition in social amoebae is mediated by allelic pairs of tiger genes. *Science* **333**: 467-470.

Iranfar N, Fuller D, Loomis WF. 2003. Genome-wide expression analyses of gene regulation during early development of Dictyostelium discoideum. *Euk Cell* **2**: 664-670.

Katoh M, Shaw C, Xu Q, Van Driessche N, Morio T, Kuwayama H, Obara S, Urushihara H, Tanaka Y, Shaulsky G. 2004. An orderly retreat: Dedifferentiation is a regulated process. *Proceedings of the National Academy of Sciences of the United States of America* **101**: 7005-7010.

Katoh-Kurasawa M, Santhanam B, Shaulsky G. 2016. The GATA transcription factor gene gtaG is required for terminal differentiation in Dictyostelium. *J Cell Sci* **129**: 1722-1733.

Keller T, Thompson CR. 2008. Cell type specificity of a diffusible inducer is determined by a GATA family transcription factor. *Development* **135**: 1635-1645.

Khare A, Shaulsky G. 2010. Cheating by exploitation of developmental prestalk patterning in Dictyostelium discoideum. *PLoS genetics* **6**: e1000854.

Kibler K, Nguyen TL, Svetz J, van Driessche N, Ibarra M, Thompson C, Shaw C, Shaulsky G. 2003. A novel developmental mechanism in Dictyostelium revealed in a screen for communication mutants. *Dev Biol* **259**: 193-208.

Knecht DA, Cohen SM, Loomis WF, Lodish HF. 1986. Developmental regulation of Dictyostelium discoideum actin gene fusions carried on low-copy and high-copy transformation vectors. *Mol Cell Biol* **6**: 3973-3983.

Kon T, Adachi H, Sutoh K. 2000. amiB, a novel gene required for the growth/differentiation transition in Dictyostelium. *Genes to Cells* **5**: 43-55.

Love MI, Huber W, Anders S. 2014. Moderated estimation of fold change and dispersion for RNA-seq data with DESeq2. *Genome biology* **15**: 550.

Miranda ER, Rot G, Toplak M, Santhanam B, Curk T, Shaulsky G, Zupan B. 2013. Transcriptional profiling of Dictyostelium with RNA sequencing. *Methods in molecular biology* **983**: 139-171.

Nichols JM, Antolovic V, Reich JD, Brameyer S, Paschke P, Chubb JR. 2020. Cell and molecular transitions during efficient dedifferentiation. *Elife* **9**.

Parikh A, Miranda ER, Katoh-Kurasawa M, Fuller D, Rot G, Zagar L, Curk T, Sucgang R, Chen R, Zupan B et al. 2010. Conserved developmental transcriptomes in evolutionarily divergent species. *Genome biology* **11**: R35.

Rosengarten RD, Santhanam B, Katoh-Kurasawa M. 2013. Transcriptional Regulators: Dynamic Drivers of Multicellular Formation, Cell Differentiation and Development. In *Dictyostelids: evolution, genomics and cell biology*, (ed. M Romeralo, et al.), pp. 89-108. Springer, Heidelberg ; New York.

Shaulsky G, Fuller D, Loomis WF. 1998. A cAMP-phosphodiesterase controls PKA-dependent differentiation. *Development* **125**: 691-699.

Shaulsky G, Loomis WF. 1993. Cell type regulation in response to expression of ricin A in Dictyostelium. *Developmental biology* **160**: 85-98.

Shimada N, Kanno-Tanabe N, Minemura K, Kawata T. 2008. GBF-dependent family genes morphologically suppress the partially active Dictyostelium STATa strain. *Devel Genes Evol* **218**: 55-68.

Stepanovic V, Wessels D, Daniels K, Loomis WF, Soll DR. 2005. Intracellular role of adenylyl cyclase in regulation of lateral pseudopod formation during Dictyostelium chemotaxis. *Euk Cell* **4**: 775-786.

Swaney KF, Huang CH, Devreotes PN. 2010. Eukaryotic chemotaxis: a network of signaling pathways controls motility, directional sensing, and polarity. *Annual review of biophysics* **39**: 265-289.

Van Driessche N, Demsar J, Booth EO, Hill P, Juvan P, Zupan B, Kuspa A, Shaulsky G. 2005. Epistasis analysis with global transcriptional phenotypes. *Nature genetics* **37**: 471-477.

Vicente JJ, Galardi-Castilla M, Escalante R, Sastre L. 2008. Structural and functional studies of a family of Dictyostelium discoideum developmentally regulated, prestalk genes coding for small proteins. *BMC Microbiol* **8:1**: 16 pages.
